# TRIM24 Degradation Counteracts Adaptation to Androgen Receptor Inhibition in Prostate Cancer

**DOI:** 10.1101/2025.08.21.671529

**Authors:** Daniela Bossi, Arianna Vallerga, Giuseppe Salfi, Nicolo’ Formaggio, Valentina Ceserani, Zhang Jichang, Tiziano Bernasocchi, Andrea Rinaldi, Lukas Bubendorf, Simone Mosole, Eva Corey, Marco Bolis, Matteo Pecoraro, Roger Geiger, Wouter R Karthaus, Jessica Barizzi, Ricardo Pereira Mestre, Jinhua Wang, Jean-Philippe Theurillat

## Abstract

The androgen receptor (AR) is the primary therapeutic target in prostate cancer. While androgen deprivation therapy (ADT) and androgen receptor signaling inhibitors (ARSi) are effective, the disease eventually progresses to fatal castration-resistant prostate cancer (CRPC). That said, little is known about the mechanisms in residual disease that initiates tumor relapse upon ADT/ARSi. Here, we discover a crucial role for TRIM24 in supporting the survival of residual cell clusters primed for tumor relapse in vivo. Consequently, reducing TRIM24 with bifunctional degraders (dTRIM24) significantly delays or even prevents the emergence of CRPC in the context of AR reactivation and lineage plasticity. dTRIM24 not only inhibits AR signaling but also counteracts adaptive pathways engaged by AR inhibition itself, such as STAT3 activation and EMT. Our findings underscore the potential of TRIM24 as an effective and druggable target for preventing prostate cancer progression under AR inhibition.

**Significance:** Despite advances in targeting AR signaling in prostate cancer, tumor relapse remains a major concern. Here, we provide evidence that more durable responses can be achieved by pharmacologically degrading TRIM24. At the molecular level, TRIM24 degradation inhibits both AR signaling and adaptive pathways that enable tumor relapse.

## Introduction

The androgen receptor (AR) is a critical oncogenic lineage-specific pathway in prostate cancer^1,2^. Upon binding to androgens, AR dimerizes and translocate into the cell nucleus, activating a transcriptional output program that supports tumor cell proliferation^3^. Consequently, androgen deprivation therapy (ADT) and more recent androgen receptor signaling inhibitors (ARSi) (e.g., second-generation anti-androgens such as enzalutamide (ENZA) or the androgen synthesis inhibitor abiraterone) remain the cornerstone of prostate cancer therapy in an early metastatic setting^4–6^. However, in many cases, tumors become resistant to ADT and ARSi and progress to castration-resistant prostate cancer disease (CRPC) with poor prognosis and fatal outcomes.

In the past twenty years, multiple resistance mechanisms to AR inhibition have been identified and can be broadly classified into two categories: (i) the restoration of AR signaling under low availability of androgens and (ii) the reprogramming and loss of the lineage-specific AR signaling through lineage plasticity. The former includes upregulation of intra-tumoral androgen synthesis, glucocorticoid receptor overexpression, genetic amplification and point mutations of the AR, the expression of AR isoforms lacking the steroid hormone-binding domain (e.g., AR-V7), and upregulation of AR coactivators^7–12^. The latter includes a variety of new emerging subtypes that are likely linked to stronger inhibition of the AR signaling through more potent ARSi^13^. Among these, neuroendocrine prostate cancer and double-negative prostate cancers are characterized by either activation of WNT-signaling or a stem-cell-like (SCL) phenotype^14–16^. These subtypes represent likely extremes of a continuum, which involves different degrees of epithelial-to-mesenchymal transition (EMT) and intermediate phenotypes (e.g., AR low)^17,18^.

While resistance mechanisms have been extensively studied by comparing cancer tissues or model organisms before and after the onset of resistance^7,15,19–21^, relatively little is known about the adaptive processes that occur in residual disease, which may subsequently prime cells for tumor relapse. Conceivably, identifying and subsequently targeting critical components may prevent disease progression or render therapeutic responses more durable. In the current study, we discover that residual cancer cells exposed to androgen deprivation require the chromatin interactor TRIM24 for tumor relapse. Moreover, we explore the therapeutic impact of TRIM24 degradation in combination with AR inhibition in castration sensitive models and as monotherapy in CRPC models in various clinically relevant settings. Our findings suggest a critical function of TRIM24 in supporting AR signaling and counteracting adaptive processes leading to disease relapse.

## Results

### TRIM24 is required for the emergence of CRPC in an LNCaP xenograft model

It has been previously demonstrated that TRIM24 supports AR signaling in CRPC by recruiting AR to cell cycle-promoting genes under low availability of androgens^7^. At the protein expression level, TRIM24 is generally associated with features of aggressive disease in primary tumors^7,22^. We further analyzed matched primary tumors and local CRPC, as well as local lymph node metastasis on tissue microarrays (TMA) by immunohistochemistry (IHC) to test if upregulation of TRIM24 is also associated with resistance to ADT or metastatic spread in individual patients. Indeed, the number of TRIM24 positive tumor cells increased in both settings, further reinforcing the link between TRIM24 and disease progression (Supp. Fig. 1A, B and Table S1A, C). Because TRIM24 chromatin binding is not exclusively regulated by protein abundance^7^, we interrogated a TRIM24 activity gene set inferred from chromatin occupancy and gene expression data on a recently generated harmonized prostate cancer RNA sequencing atlas (www.prostatecanceratlas.org). We found that TRIM24 activity steadily increases from normal to primary and CPRC and the activity correlated with a continuous pseudotime progression score (Supp. Fig. 1C, D)^7,21^. The signature was significantly associated with various driver alterations in primary disease in the TCGA cohort (e.g., TP53, FOXA1, SPOP (Supp. Fig. 1E), while we noted at least a trend towards the loss of RB1/TP53/PTEN copies and gains of MYC, in both the TCGA and WCDT CRPC cohort (Supp. Fig. 1F)^23,24^. Despite a significant association with poor survival in the latter cohort, the signature did not emerge as an independent prognostic factor in a multivariate analysis, suggesting that TRIM24 activity may be linked to complex interactions of genomic alterations related to disease progression and androgen independence (Supp. Fig. 1G, H).

Given these findings, we explored the role of TRIM24 in the adaptation process to androgen withdrawal. For this purpose, we used the AR-positive LNCaP xenograft mouse model in which androgen deprivation by surgical castration leads to complete tumor regression in 4 weeks, followed by tumor relapse within 1-2 months (Fig. 1A)^21^. Using single-cell RNA sequencing (scRNA-seq), we longitudinally characterized the pathways in this model. The analysis of epithelial and stromal markers confirmed successful separation of the murine tumor microenvironment from the human tumor compartment. The latter was used for subsequent analyses (Supp. Fig. 2). Tumor relapse was associated with restoration of AR and mTOR signaling, as well as the activation of multiple signaling pathways related to EMT process, such as IL6-Jak-STAT3, TNFA-NFKB, TGFβ, and hypoxia (Fig. 1B & Supp. Fig. 3A)^17,18, 25^. To validate our model, we derived a progression signature from the tumor cells before castration and upon relapse and applied it to single cell data generated in human tumors before and after enzalutamide (ENZA) treatment (Table S2)^26^. Indeed, the progression signature was significantly enriched in the relapsed tumors, indicating the clinical relevance of the model (Supp. Fig. 3B).

**Figure 1.**
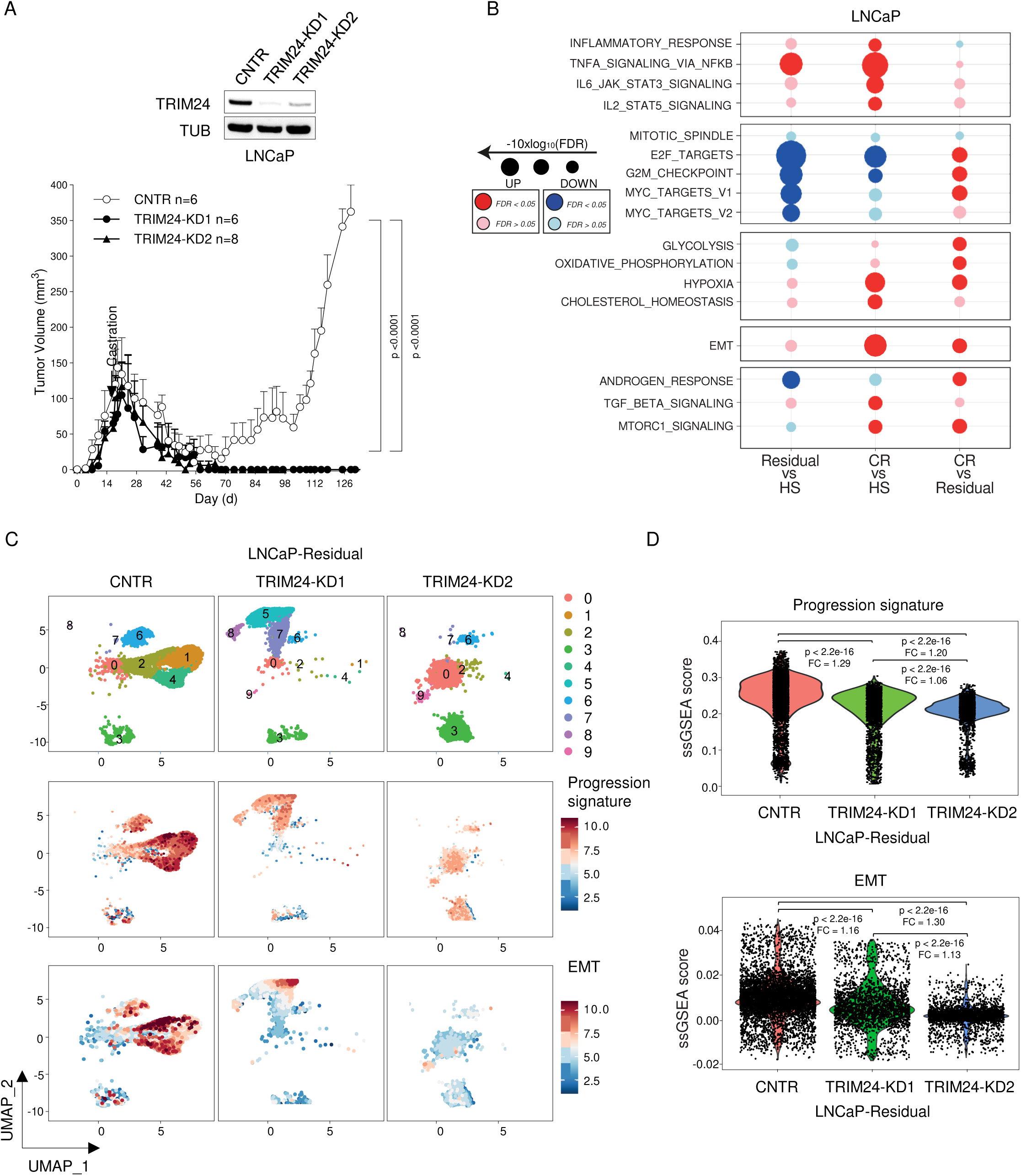
Silencing of TRIM24 prevents tumor relapse after castration. **A** Upper panel, immunoblot analysis of the indicated proteins showing TRIM24 knockdown of LNCAP cells transduced with lentiviral shSCR (control, CNTR), shTRIM24-1 (TRIM24-KD1), and shTRIM24-2 (TRIM24-KD2) Lower panel, growth kinetics of CNTR, TRIM24-KD1, and TRIM24-KD2 xenografts in NRG mice. Castration (arrow) was performed when the tumor reached an average size of 50-75 mm3. Statistical significance is determined using a Two-way ANOVA test. **B** Gene set enrichment analysis (GSEA) of Hallmark signatures using pseudo-bulk scRNA-seq (scRNA-seq) data from LNCaP-HS (Hormone-Sensitive, pre-castration), LNCaP-Residual (3 weeks post-castration), and LNCaP-CR (Castration-Resistant, relapse post-castration). The dot plot illustrates the gene-enriched scores’ false discovery rate (FDR) for all comparisons. **C** Uniform Manifold Approximation and Projection (UMAP) plot showing the distribution of cell clusters (upper panel) and the gene set enrichment scores of the progression signature (middle panel; see methods and Supplementary Figure 3B) and the Epithelial-Mesenchymal-Transition (EMT) signature (lower panel) in LNCaP-Residual tumors sampled 3 weeks after castration, comparing CNTR (left), TRIM24-KD1 (middle, on treatment), and TRIM24-KD2 (right). D Violin plot of the single-sample enrichment score (ssGSEA) for the progression and EMT signature computed for each residual tumor cell. Median pathway scores for each group are indicated, and fold-change values (ratio of medians) are annotated between groups. Statistical significance was assessed using the Wilcoxon rank-sum test.

Next, we generated LNCaP control cells and TRIM24 knockdown cells using two stables short-hairpin RNAs (KD1, KD2, Fig. 1A). Following our previous report, knockdown of TRIM24 just minorly affected tumor growth under androgen-rich condition^7^. In stark contrast, TRIM24 KD completely abolished tumor regrowth after castration, indicating a crucial role in this process (Fig. 1A). To investigate why TRIM24 loss-of-function prevented tumor relapse, we interrogated the scRNA-seq data of the residual disease of control and TRIM24-KD1 and TRIM24-KD2 jointly. The knockdown of TRIM24 had a major influence on the distribution of cell clusters. Most notably, we observed that clusters 1, 2, and 4 largely disappear with both short hairpins (Fig. 1C).

To understand if these clusters may have been critical for tumor relapse, we analyzed the disease progression signature across all clusters. Indeed, cells within clusters 1, 2, and 4 displayed a significantly higher enrichment score, while scores decreased in clusters 5, 7, 8, and 9 unique to TRIM24 KD samples (Fig. 1C, D & Supp. Fig. 4A). Moreover, the analysis of the signatures across clusters revealed that cluster 4 had by far the highest level of AR and EMT signaling – the latter feature had been previously associated with Trim24 over-expression in a genetically engineered mouse model for metaplastic breast cancer (Fig. 1 C, D & Supp. Fig. 4B)^27^. In line with a functional relationship of AR and TRIM24, we also found a good overlap between AR and TRIM24 expression in clusters 1, 2, and 4 (Supp. Fig. 4C, D). In aggregate, the data suggests that TRIM24 depletion abrogates tumor regrowth by preventing the emergence of cell clusters with sustained AR signaling and the engagement of EMT associated markers. Given the importance of TRIM24 for cancer cells to adapt to ADT, we investigated if TRIM24 protein level or the activity gene set may predict response to ADT/ARSi. For this purpose, we compared TRIM24 protein abundance in prostate biopsies of patients with de novo, high volume metastatic disease and either poor or exceptional response to ADT +/- ARSi, defined as ≤ 9 months or more than 38 months to radiographic disease progression (Table S1E). Despite considering only high-grade tumors (i.e., Gleason score of 8-10), the number of TRIM24 positive nuclei were significantly higher among the poor responders (Supp. Fig. 5A). Moreover, a high TRIM24 activity signature showed a trend towards poor survival of CRPC patients of the WCDT cohort prior abiraterone or ENZA treatment (Supp. Fig. 5B, C)^24^, suggesting in aggregate that targeting TRIM24 in prostate cancer patients with TRIM24 protein expression/activity may help prolong responses to ADT/ARSi.

### TRIM24 degradation suppresses AR signaling under ADT/ARSi

The recent development of bifunctional TRIM24 protein degraders by the VHL-EloBC-Cul2 complex enabled us to interrogate whether a pharmacological degradation may have the same effect as the TRIM24 knockdown on tumor relapse^28^. To this end, we tested the published TRIM24 degrader, dTRIM24-1 and a derivate with enhanced solubility dTRIM24-2 in cell culture (Fig. 2A). Both molecules efficiently degraded TRIM24 in cell culture in androgen-dependent LNCaP, VCaP, and LAPC4 cells at the low micromolar (dTRIM24-1) and nanomolar range (dTRIM24-2), respectively (Fig. 2B). In line with our previous study suggesting a critical role for TRIM24 in supporting AR signaling and tumor growth under low availability of androgens^7^, clonogenic growth of all three cell lines was reduced with dTRIM24-1/2. This reduction occurred when LNCaP cells were cultured under androgen-deprived condition with charcoal-stripped serum (CSS) alone or in the presence of enzalutamide but not under substitution with dihydrotestosterone (DHT) (Fig. 2C, D). Consistently, dTRIM24-1/2 also decreased AR signaling, as evidenced by reduced NKX3.1 expression, under CSS and CSS/ENZA but not under DHT (Fig. 2E & Supp. Fig. 6A). Similar results were produced for TRIM24-KD in LNCaP cells (Supp. Fig. 6B, C). The data points to the specific dependency of androgen-dependent prostate cancer cells to TRIM24 degradation when treated with ADT alone or in combination with ARSi.

**Figure 2.**
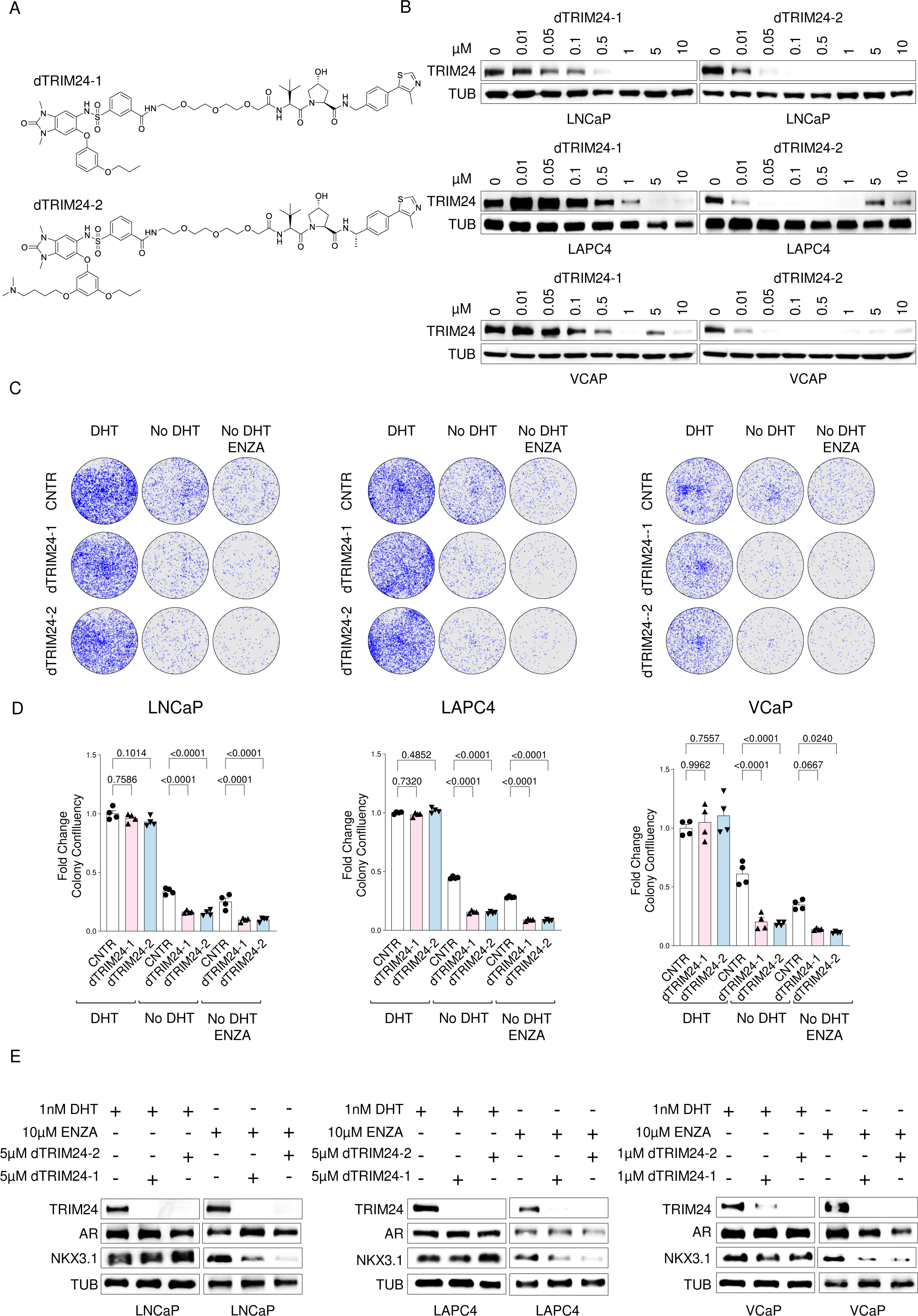
Degradation of TRIM24 reduces the growth of prostate cancer cell lines under AR inhibition. **A** Chemical structure of dTRIM24-1(upper panel) and dTRIM24-2 (lower panel). **B** Immunoblot analysis of the indicated proteins of LNCaP (upper panel), LAPC4 (middle panel), and VCAP (lower panel) cells incubated with the indicated concentrations of dTRIM24-1 and dTRIM24-2 for 48h. **C-D** Colonies formation assay for LNCaP (left), LAPC4 (middle), and VCAP (right) maintained for 21 days in: CSS (Charcoal-Stripped Serum, testosterone-free) medium with 1 nM DHT (DHT), CSS medium alone (no DHT), and CSS medium with 10 µM ENZA (no DHT + ENZA), along with dTRIM24-1 and dTRIM24-2 treatments. The concentrations used for the dTRIM24-1 and-2 were 5 µM for both LNCaP and LAPC4 cells, and 1 µM for VCAP cells. **C**, representative image of colony confluency; **D**, bar plot showing the fold change in colony confluency for the indicated cell lines at the indicated condition. The fold change is calculated based on the average colony confluency observed in the DHT condition. This data is derived from four independent experiments, with the bars representing the mean and the Standard Error of the Mean (SEM). Statistical significance was evaluated using a one-way ANOVA test. **E** Immunoblot analysis of indicated proteins in LNCaP (left panel), LAPC4 (middle panel), and VCAP (right panel) cultures for 3 days in CSS medium with DHT (lane1) or enzalutamide (lane 4) alone or in combination with dTRIM24-1 (lane 2,5) or dTRIM24-2 (lane 3,6) at the indicated concentration.

### TRIM24 degradation delays the onset of CRPC in an LNCaP xenograft model

We next set out to test the activity of dTRIM24-1/2 in vivo. To explore the feasibility of evaluating toxic side effects in vivo, we tested the capacity of the degraders to decrease murine Trim24. In cell culture, dTRIM24-2 demonstrated superior effectiveness across a broader range of concentrations compared to dTRIM24-1, which displayed a strong Hook effect in murine TRAMP-C1 cells (Supp. Fig. 6D). In vivo, the dTRIM24-2 was again more effective. It reduced Trim24 in brain, lung, liver, spleen, and prostate homogenates while we did not observe any effect in the heart, blood, or kidney (Supp. Fig. 6E, F).

Next, we evaluated the in vivo activity of dTRIM24-1 and −2 in comparison to ENZA in the LNCaP xenograft model. Intraperitoneal administration of both compounds reduced TRIM24 protein abundance also in vivo (Fig. 3A), although less pronounced than those observed in cell culture (Fig. 2B) or after TRIM24 knockdown (Fig. 1A). After castration, we treated the mice for 3 weeks and then monitored tumor growth until relapse occurred. While ENZA slightly delayed disease progression, dTRIM24-1 and −2 exhibited both a more pronounced effect and a slower kinetic of tumor relapse (Fig. 3B). In addition, we did not observe any signs of toxicity or weight loss, and there was no effect on morbidity under treatment (data not shown; see below).

**Figure 3.**
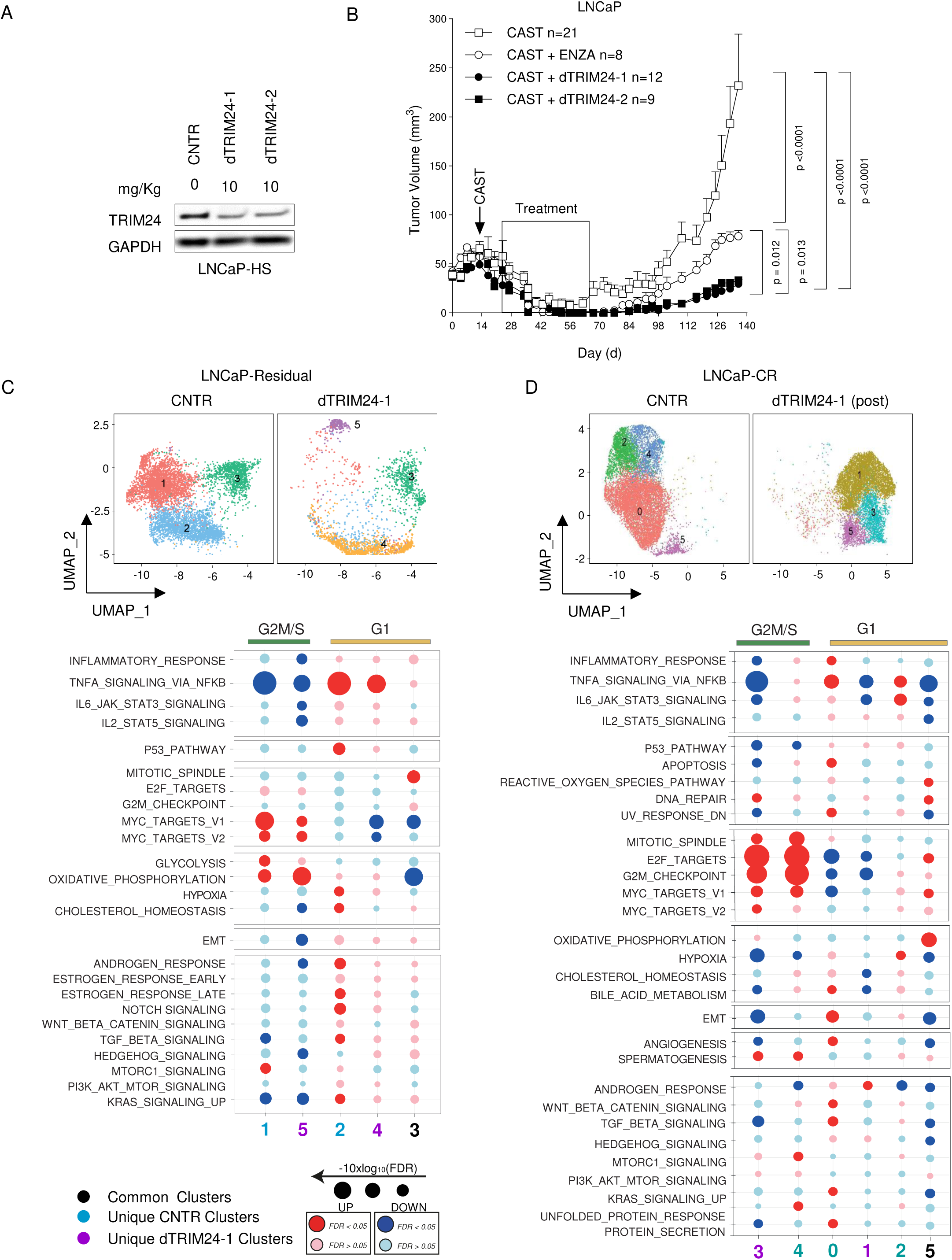
TRIM24 is required to trigger STAT3 signaling and EMT upon AR inhibition. **A** Immunoblot analysis of indicated proteins of LNCaP cells maintained for 3 days in CSS medium alone (lane 2), in combination with DHT (CSS+DHT, lane 1), or with ENZA (CSS+ENZA, lane 3) at the indicated concentration. **B** Corresponding immunoprecipitation (IP) of TRIM24 and IgG control in LNCaP cells followed by immunoblotting for TRIM24 and phospho-STAT3. On the right, immunoblot analysis of input lysates for the indicated proteins. **C** Immunoblot analysis of indicated proteins of LNCaP cells maintained for 3 days in CSS medium with ENZA (CSS+ENZA, lane 1) alone or in combination with dTRIM24-1 (CSS+ENZA+dTRIM24-1, lane 2) or dTRIM24-2 (CSS+ENZA+dTRIM24-2, lane 3) at the indicated concentration. **D** Bar plot showing mRNA expression analysis of the indicated genes in LNCaP cells cultured for 3 days in CSS+DHT, CSS+ENZA (white bar), CSS+ENZA+dTRIM24-1 (light pink bar), or CSS+ENZA+dTRIM24-2 (light blue bar) by qPCR. The fold change is calculated based on the average expression in the DHT condition (dashed gray line). Data reported from at least 3 independent experiments, bars represent mean with Standard Error of Mean (SEM). Statistical significance is determined using an unpaired Student’s t-test. **E** Trans-well invasion assay of LNCaP cells maintained for 3 days in CSS+DHT (CNTR), CSS+ENZA, CSS+ENZA+dTRIM24-1, and CSS+ENZA+dTRIM24-2 and then incubated in a Boyden chamber for 24 hours. Migrated cells were stained with crystal violet, and the cluster numbers were quantified. The bars report the mean (SEM) of 4 independent experiments. Statistical significance is determined using an unpaired Student’s t-test. Representative images of the transwell migration assay of LNCaP cells stained with crystal violet are shown on the right (magnification 10X). **F** Wound healing assay by live cell imaging using the IncuCyte system of LNCaP in different medium conditions. LNCaP cells are maintained for 3 days in CSS+DHT (CNTR), CSS+ENZA, CSS+ENZA+dTRIM24-1, and CSS+ENZA+dTRIM24-2, scratched, and then tested for 120 hours. The graph displays the relative wound density of LNCaP cells at different time points. The bars represent the mean (SEM) of a single representative experiment performed in triplicate. Representative microscope images are shown on the left (magnification 10X). White line: wound at timepoint 0, red line: wound at timepoint 120h. Statistical significance is determined using an unpaired Student’s t-test.

To better understand the similarities and differences induced at the molecular level by ENZA and dTRIM24-1 treatment, we performed scRNA-seq and analyzed the residual disease of the human tumor compartment (Supp. Fig. 7). While no changes were observed in the cell cluster organization (Supp. Fig. 8A), ENZA effectively inhibited AR signaling and cell cycle progression, as expected (Supp. Fig. 8B, C). In addition, it also engaged pathways linked to disease progression (e.g., EMT-related markers, IL6-JAK-STAT3) and metastasis (e.g., hypoxia, angiogenesis) (Supp. Fig. 8B, C), in line with previous reports^17,18,25,29,30^. In comparison, in the dTRIM24-1 treatment, the cell cluster clusters were completely rewound and associated with more pronounced transcriptional changes (Fig. 3C). Specifically, two clusters (4 and 5) emerged, and like ENZA, we observed a reduction of AR signaling and cell cycle at the transcriptional level when comparing the G2M/S cluster (1 versus 5) and the G1 clusters (2 versus 4) (Fig. 3C and Supp. Fig 8D). Moreover, pathways related to disease progression were suppressed, such as mTOR, EMT-related markers, IL6-JAK-STAT3, and TGF-beta (Fig. 3C & Supp. Fig 8D). In agreement, we found in our previous matched IHC analysis that STAT3 phosphorylation significantly increased as well among the group of CRPC with increased TRIM24 expression (Supp. Fig. 1A, Table S1B). Our findings suggest that the degradation of TRIM24 significantly delays disease progression by co-targeting both AR signaling and adaptive mechanisms leading to tumor relapse.

Next, we set out to investigate why the TRIM24 knockdown exhibited more pronounced disease relapse than the degraders. We analyzed the scRNA-seq data from both experiments and identified cluster 1 as the cluster likely primed for disease progression due to its high expression of TRIM24, AR, EMT, TGF-beta, and IL6-JAK-STAT3 (Supp. Figs. 9 & 10A, B). In line, the knockdown of TRIM24 resulted in a higher reduction of cells in this specific cluster compared to dTRIM24-1 (Supp. Fig. 10C). This observation is consistent with the fact that cluster 1 also had a high score for disease progression signatures, which was further elevated when comparing dTRIM24-1 with TRIM24 knockdown (Supp. Fig. 9). Finally, immunoblot analysis of residual disease 3 weeks post castration revealed a stronger reduction of TRIM24 in the context of TRIM24 knockdown that was paralleled by stronger reduction in AR and phosphorylation of STAT3 (Supp. Fig. 10D). The findings may partially explain the distinct outcomes between dTRIM24-1 and TRIM24 knockdown, suggesting that developing more potent degraders can lead to better therapeutic benefits. Because the kinetics of tumor relapse was slower after the treatment with the TRIM24 degraders compared to the control group (castration alone) (Fig. 3B), we hypothesized that the regrowing tumor evolves differently post treatment. Indeed, the scRNA-seq analysis of control and dTRIM24-1 at tumor relapse showed significant differences in tumor cell cluster organization (Fig. 3D). Most importantly, the clusters 3 and 1 specific to dTRIM24-1 treatment displayed much lower activity of the EMT-related markers and associated pathways such as TNFA-NFKB, IL6-JAK-STAT3, TGF-beta, and WNT-beta catenin (Fig. 3D & Supp. Fig. 11A). This highlights the potential of dTRIM24-1 treatment to affect these crucial pathways. In line with this, STAT3 phosphorylation was also reduced in the relapse post dTRIM24-1 and −2 administration (Supp. Fig. 11B). In contrast, we also found that the dTRIM24-1 treatment induces high levels of AR signaling only in the slow-cycling cells of cluster 1 in the relapsed tumor post treatment, suggesting that the tumor cells can have progressed by upregulating AR signaling while failing to engage adequately EMT-related markers and associated pathways such as IL6-JAK-STAT3 (Fig. 3D & Supp Fig. 11A). Resistance mechanisms of critical target inhibition are often directly linked to the target activity itself. In line with this, we found a significant increase in TRIM24 protein levels at the time point of relapse (Supp. Fig. 11B). We observed a change in the transcriptional level of TRIM24 expression during the different stages of tumor progression. However, the change in the mRNA level of TRIM24 in the longitudinal tumor samples did not match the protein level changes (Supp. Fig. 11A), implying other post-transcriptional mechanisms involved in TRIM24 upregulation. Because TRIM28 has been shown to stabilize TRIM24 in prostate cancer^31^, we next interrogated TRIM28 expression in the relapsed tumors. Indeed, TRIM28 mRNA levels increased specifically in the cells with the restoration of AR and mTOR signaling after dTRIM24-1 treatment (Supp. Fig. S11A). In addition, we confirmed increasedTRIM28 protein levels and a decreased STAT3 phosphorylation at relapsed tumors after the treatment with the degraders (Supp. Fig. S11B). These results may indicate that TRIM28 can stabilize TRIM24 to restore AR signaling and promote tumor relapse in the absence of the induction of EMT-related markers.

### TRIM24 counteracts STAT3 signaling and the release of EMT-promoting cytokines

Tumor relapse in the LNCaP xenograft model was associated with an increased activation of the IL6-JAK-STAT3 signaling, an upstream pathway linked to EMT process, disease progression, and lineage plasticity in prostate cancer^18,25,32–35^. Therefore, we investigated whether STAT3-signaling activation is a direct consequence of AR inhibition. Indeed, LNCaP cells cultured in the absence of DHT and the presence of ENZA exhibited increased STAT3 phosphorylation and promoted expression of EMT-associated markers (i.e., upregulation of N-Cadherin and downregulation of E-Cadherin) (Fig. 4A). Because TRIM24 has been shown to interact and activate STAT3 in glioblastoma^36^, we set out to investigate whether androgen deprivation and ENZA also elicit the interaction between TRIM24 and STAT3 in prostate cancer. In agreement, we found that more phosphorylated and total STAT3 were more bound to TRIM24 when cells were cultured in the absence of DHT and the presence of ENZA (Fig. 4B & Supp. Fig. 12A). In line with this, both TRIM24 knockdown or dTRIM24-1/2 reduced STAT3 phosphorylation and decreased N-Cadherin protein levels while over-expression had the opposite effect (Figs. 4C & Supp. Fig. 12B, C). Suppression of STAT3 activation in cell culture was specific to ENZA and not observed under DHT-rich condition (Supp. Fig. 12D, E) while there was no robust effect under CSS only (data not shown). Similar results were obtained in LAPC4, and VCaP cells with dTRIM24-1/2 (Supp. Fig. 12F-I). In line with a reduction with STAT3 activation, we found corresponding changes in mRNAs of canonical STAT3 and EMT-associated cytokines (i.e., IL-6, TGFbeta1, TNFA, IL-10, IL-1beta) with both the knockdown and dTRIM24-1/2 in LNCaP and with dTRIM241/2 in VCaP and LAPC4 cells as well (Figs. 4D & Supp. Fig. 13A-C).

**Figure 4.**
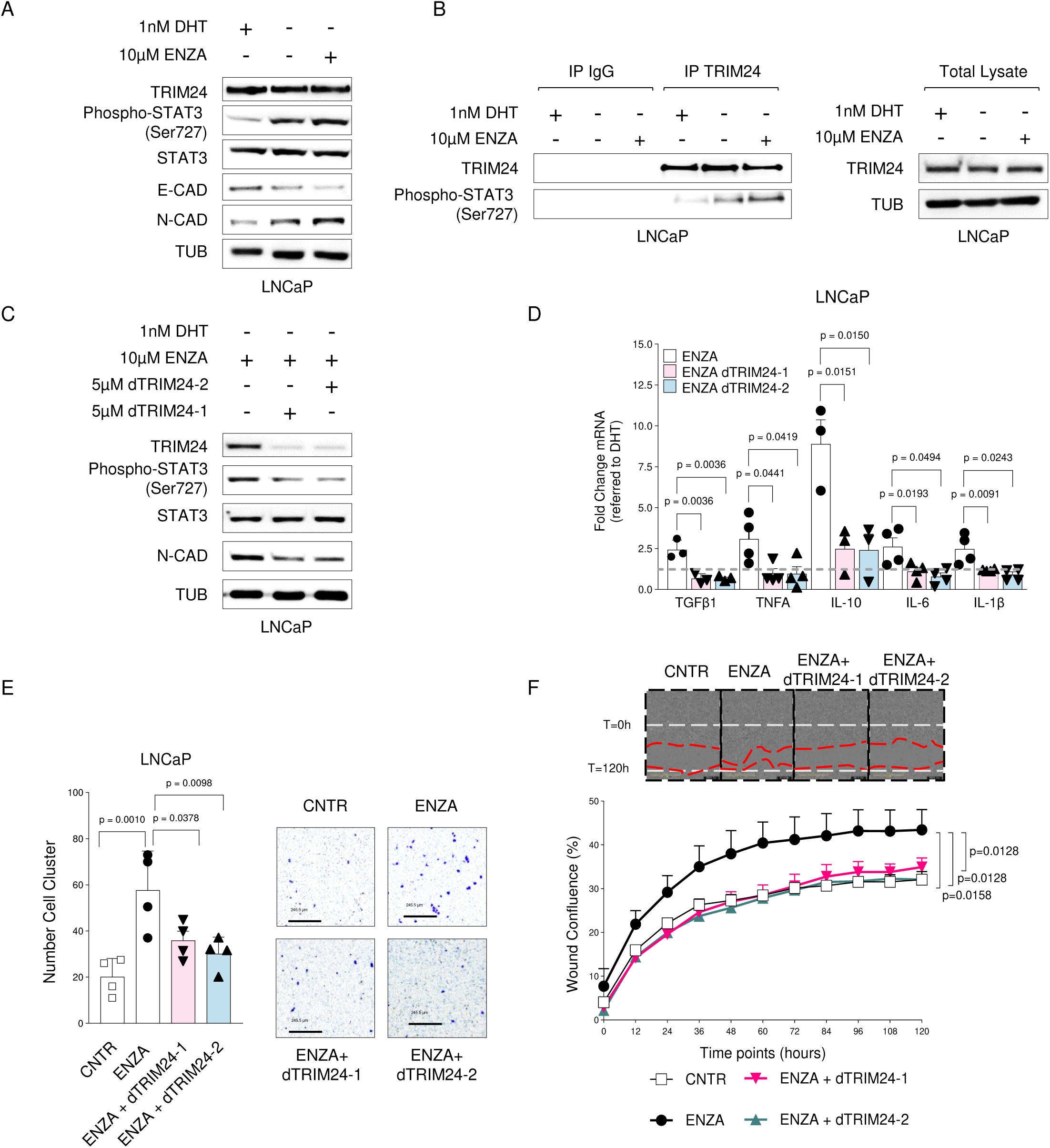
TRIM24 is required to trigger STAT3 signaling and EMT upon AR inhibition. **A** Immunoblot analysis of indicated proteins of LNCaP cells maintained for 3 days in CSS medium alone (lane 2), in combination with DHT (CSS+DHT, lane 1), or with ENZA (CSS+ENZA, lane 3) at the indicated concentration. **B** Corresponding immunoprecipitation (IP) of TRIM24 and IgG control in LNCaP cells followed by immunoblotting for TRIM24 and phospho-STAT3. On the right, immunoblot analysis of input lysates for the indicated proteins. **C** Immunoblot analysis of indicated proteins of LNCaP cells maintained for 3 days in CSS medium with ENZA (CSS+ENZA, lane 1) alone or in combination with dTRIM24-1 (CSS+ENZA+dTRIM24-1, lane 2) or dTRIM24-2 (CSS+ENZA+dTRIM24-2, lane 3) at the indicated concentration. **D** Bar plot showing mRNA expression analysis of the indicated genes in LNCaP cells cultured for 3 days in CSS+DHT, CSS+ENZA (white bar), CSS+ENZA+dTRIM24-1 (light pink bar), or CSS+ENZA+dTRIM24-2 (light blue bar) by qPCR. The fold change is calculated based on the average expression in the DHT condition (dashed gray line). Data reported from at least 3 independent experiments, bars represent mean with Standard Error of Mean (SEM). Statistical significance is determined using an unpaired Student’s t-test. **E** Trans-well invasion assay of LNCaP cells maintained for 3 days in CSS+DHT (CNTR), CSS+ENZA, CSS+ENZA+dTRIM24-1, and CSS+ENZA+dTRIM24-2 and then incubated in a Boyden chamber for 24 hours. Migrated cells were stained with crystal violet, and the cluster numbers were quantified. The bars report the mean (SEM) of 4 independent experiments. Statistical significance is determined using an unpaired Student’s t-test. Representative images of the transwell migration assay of LNCaP cells stained with crystal violet are shown on the right (magnification 10X). **F** Wound healing assay by live cell imaging using the IncuCyte system of LNCaP in different medium conditions. LNCaP cells are maintained for 3 days in CSS+DHT (CNTR), CSS+ENZA, CSS+ENZA+dTRIM24-1, and CSS+ENZA+dTRIM24-2, scratched, and then tested for 120 hours. The graph displays the relative wound density of LNCaP cells at different time points. The bars represent the mean (SEM) of a single representative experiment performed in triplicate. Representative microscope images are shown on the left (magnification 10X). White line: wound at timepoint 0, red line: wound at timepoint 120h. Statistical significance is determined using an unpaired Student’s t-test.

We further investigated effects of ADT/ENZA and dTRIM24 on cytokine secretion. While the cytokines mentioned above were barely detected in the supernatant, androgen deprivation and ENZA robustly increased the levels of cytokines linked to cancer progression, such as FGF-19, GDF-15, MIF, IGFBP2 (Supp. Fig. 13D, E and Table S3)^37–40^. Importantly, the addition of dTRIM24-1/2 to CSS and ENZA largely prevented the induction of cytokines. The observations were further validated by corresponding mRNA expression changes in the tumor cells (Supp. Fig. 13F). Finally, we asked whether dTRIM24-1 and −2 exhibits effects on cell invasion and migration, two processes typically associated with STAT3 signaling and EMT in cell culture. In agreement with our previous findings, androgen deprivation and ENZA induced cell invasion and migration in LNCaP cells, while co-administration with dTRIM24-1 nd −2 or TRIM24 KD largely abrogated the effect (Fig. 4E, F & Supp. Fig. 13G, H). Together, the findings strongly suggest that ADT/ARSi, increases STAT3 signaling and EMT-related markers, at least in part through TRIM24, and these effects may be attenuated by TRIM24 degradation.

### TRIM24 degradation inhibits the growth of AR-V7-driven prostate cancer models

Restoration of AR signaling in CRPC is frequently achieved by alterations of AR itself which in turn sustain its activity when testosterone levels are low. For example, LNCaP cells harbor the AR mutation T877A mutation which opens the binding repertoire of steroid hormones^10^. Another prevalent adaptation mechanism involves the expression of AR isoforms lacking the steroid hormone-binding domain (e.g., AR-V7)^11,41^. To test whether TRIM24 degradation also affect AR-V7 signaling, we investigated effects of TRIM24 KD or dTRIM24-1 and −2 in LNCaP-95, an androgen-independent LNCaP subline, as well as in 22Rv1 cells both of which express AR-V7 (Fig. 5A). Both degraders efficiently lowered TRIM24 protein abundance in both cell lines. The degradation with dTRIM24-2 was superior across a wider range of concentrations, as expected (Supp. Fig. 14A). The decrease in TRIM24 by the degraders was further confirmed by mass-spectrometry for LNCaP-95, where TRIM24 emerged as the by far most downregulated protein (Fig. 5B). In line, the principal component analysis of the corresponding RNA-seq data displayed coordinate changes related to dTRIM24-1/2 (Supp. Fig. 14B and Table S4). Further mining of the RNA-seq data, revealed a reduction in AR signaling for both dTRIM24-1 and −2 in LNCaP-95 cells (Supp. Fig. 14C). Interestingly, we found that AR-V7 was substantially reduced in both LNCaP-95 and 22Rv1 cells upon dTRIM24-1/2 treatment and TRIM24 knockdown. This reduction was observed at both the protein and the mRNA level. Additionally, we noted a reduction in NKX3.1 and the AR-V7 target genes SCL3A2 and NUP210^42^, and mTOR signaling was affected as evidenced by a reduction in phosphorylation of S6K1 and 4-EBP1 (Figs. 5C-E & Supp. Fig. 14D-F). In addition, TRIM24 loss-of-function was consistently associated with the downmodulation MYC protein abundance and corresponding target genes in the RNA sequencing data (Figs. 5C & Supp. Fig. 14C, D). The latter finding was not recapitulated in LNCaP cells by dTRIM24-1/2, suggesting a potential specific link of MYC regulation in AR-V7-driven models (Supp. Fig. 14G). Conversely, we did not note a negative regulation of STAT3 phosphorylation or signaling in LNCaP-95 cells upon dTRIM24-1/2 treatment, previously noted upon adaptation of androgen-dependent prostate cancer cell lines to CSS/ENZA (Figs. 4A & Supp. Figs. 14C, H). In agreement with the suppression of key oncogenic pathways in LNCaP-95 and 22Rv1 cells, both dTRIM24-1/2 or TRIM24 knockdown reduced clonogenic growth (Figs. 5F, G & Supp. Fig. 14I). To further substantiate the translational potential of dTRIM24-1 and −2, we compared the effects with ENZA in vivo in LNCaP-95 xenografted tumors. After five days of continuous intraperitoneal administration, both compounds reduced TRIM24 levels (Fig. 5H). We then treated the mice for two weeks. We monitored tumor growth during the exponential growth phases. ENZA had a weak effect, as expected, but dTRIM24-1 and −2 efficiently reduced tumor growth without any signs of toxicity (Figs. 5H & Supp. Fig. 14J). These results may indicate that dTRIM24-1 and −2 may also counteract the restoration of AR signaling driven AR-V7 when combined upfront with ADT/ARSi.

**Figure 5.**
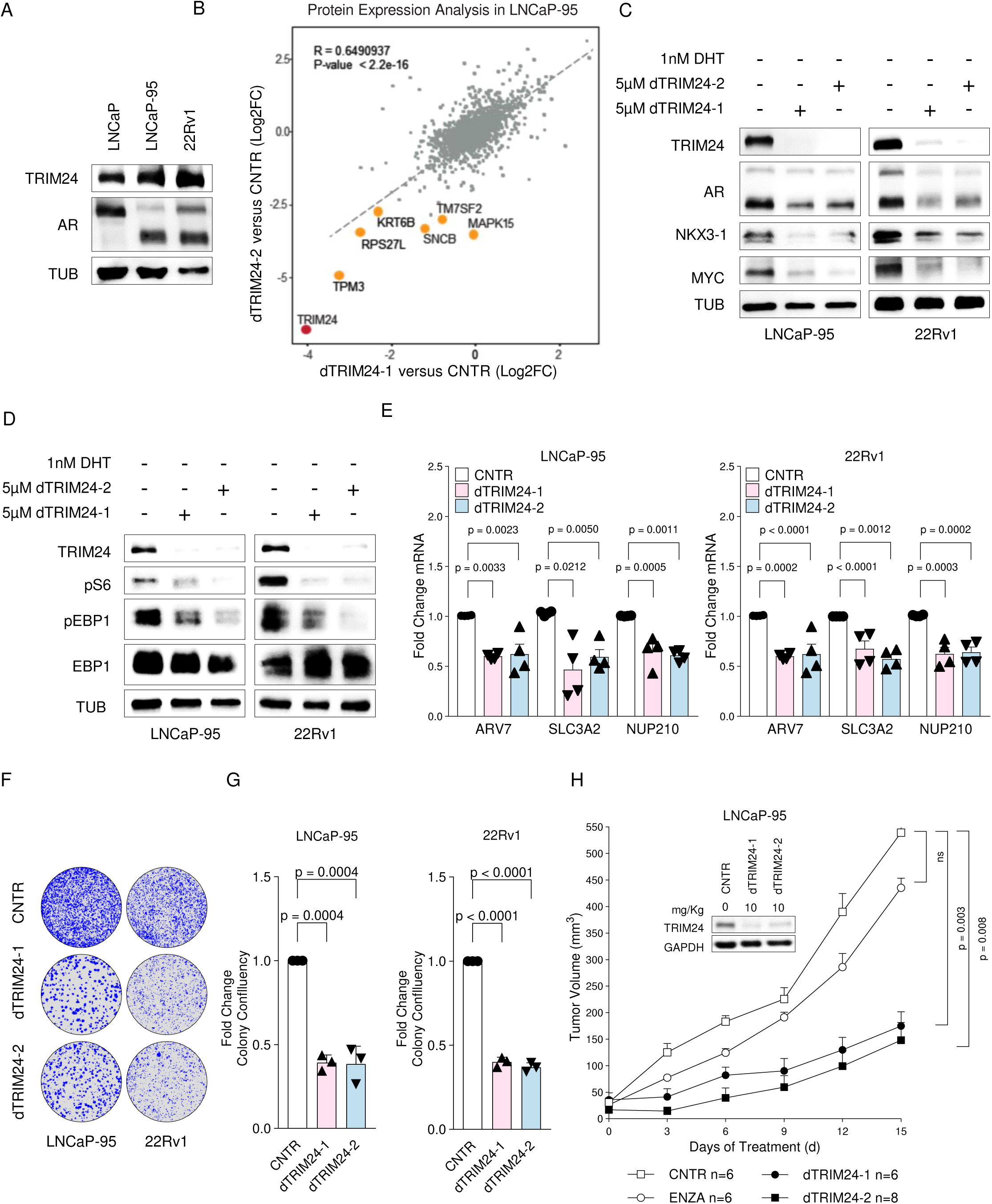
Degradation of TRIM24 inhibits the growth of the AR-V7-driven LNCaP-95 and 22Rv1 cells. **A** Immunoblot analysis of the indicated proteins in parental LNCaP (lane 1), LNCaP-95 (lane 2), and 22Rv1 (lane 3) cells. **B** A scattered plot showing log2 fold changes (FC) in protein levels after treating LNCaP-95 cells in CSS medium alone (CNTR) or with 5 μM dTRIM24-1 and dTRIM24-2 for 48 hours. The correlation of the fold changes was calculated using the Pearson correlation coefficient. **C-D** Immunoblot analysis of indicated proteins in LNCaP-95 (left panel) and 22Rv1 (right panel) maintained for 3 days in CSS medium alone (CNTR, lane 1) or supplemented with dTRIM24-1 (lane 2) or dTRIM24-2 (lane 3) at the indicated concentration. **E** Bar plot showing mRNA expression analysis of AR-V7 target genes in LNCaP-95 (left) and 22RV1 (left) cells cultured for 3 days in CSS medium alone (CNTR, white bar) or with 5 µM of dTRIM24-1 (light pink bar) and dTRIM24-2 (light blue bar) by qPCR. The fold change is calculated based on the average expression in the control condition. Data reported from at least 3 independent experiments, bars represent mean with Standard Error of Mean (SEM). Statistical significance is determined using an unpaired Student’s t-test. **F-G** Colony formation assay for LNCaP-95 (left), 22Rv1 (right) maintained for 21 days in CSS medium alone (CNTR) or with 5 µM of dTRIM24-1 and dTRIM24-2. In **F**, a representative image of colony confluency; in **G**, a bar plot showing fold change in colony confluency for both cell lines at the indicated condition. The fold change is calculated based on the average colony confluency observed in the control condition. This data is derived from four independent experiments, with the bars representing the mean and the Standard Error of the Mean (SEM). Statistical significance was evaluated using a one-way ANOVA test. **H** Upper panel, immunoblot analysis of the indicated proteins showing TRIM24 reduction in LNCaP-95 xenografts after daily intraperitoneal administration of dTRIM24-1 and dTRIM24-2 at 10mg/kg for five days. Lower panel, growth kinetics of LNCaP-95 xenografts in castrated NRG mice treated daily intraperitoneally with the vehicle (white square), 10 mg/kg dTRIM24-1 (black circle), and −2 (black square), and 30 mg/kg enzalutamide (white circle) for two weeks. The statistical significance of the results was tested using a two-way ANOVA test.

### TRIM24 degradation inhibits EMT markers in the context of TP53 and RB1 deficiency

Genetic loss-of-function mutations in p53 and RB1 are intimately linked to aggressive prostate cancer, EMT markers, and lineage plasticity^35,43,44^. Considering this, we set out to investigate whether TRIM24 degradation, in combination with castration and ENZA, can significantly affect the inhibition of tumors with alterations in p53 and RB1. We utilized LNCaP cells with CRISPR-mediated deletion of both tumor suppressor genes to test this (Supp. Fig. 15A)^43^. Upon castration, the tumors progressed rapidly to CRPC, and ENZA showed little additional benefit in this context (Supp. Fig. 15B). RNA-seq of LNCaP^TP53-/-; RB1-/-^ before and after castration revealed an increase in EMT-associated markers in the tumor relapse, especially when co-treated with ENZA (Supp. Fig. 15C). Additionally, gene expression signatures related to the NEPC and WNT pathways were enriched, along with decreased AR-related signatures (Supp. Fig. 15D, E). Treatment with dTRIM24-2, combined with ENZA and castration, led to increased tumor growth inhibition and significantly delayed tumor relapse. (Fig. 15F). As expected, AR expression was lower in both the control and treated tumors upon relapse (Fig. 15G). In contrast, the relapsed control tumors exhibited elevated protein levels of the increased mesenchymal marker N-cadherin (Fig. 15G). Remarkably, adding dTRIM24-2 counteracted the mesenchymal marker induction caused by androgen deprivation and ENZA in vivo, however, in this case unrelated to STAT3 activation/phosphorylation.

### TRIM24 degradation suppresses cancer progression associated with lineage plasticity

Subsequently, we explored whether disease progression through lineage plasticity is impacted by TRIM24 degradation. Following castration, the androgen-dependent patient-derived xenograft (PDX) model LuCaP147 transitions to AR-negative, stem-cell-like (SCL)-CRPC prostate cancer in immune-compromised NRG mice (Fig. 6A, B & Supp. Fig. 16A-B)^14^. We performed DNA exome sequencing before and after castration to determine if the switch to SCL-CRPC involved clonal selection or an adaptive process. The analysis revealed neither a significant change in the allelic fraction of prostate cancer driver mutations (e.g., AR, SPOP, BRCA2, PTEN, RNF43, STK11) nor the emergence of additional putative driver mutations in the relapse (Supp. Fig. 16C & Table S6)^45^, suggesting an adaptive process underlying lineage plasticity in this model. In support, we found an increase of mesenchymal markers and an enrichment of EMT-related gene set in the relapse (Fig. 6A, C). Because TRIM24 protein abundance did not increase during the regrowth after castration (Fig. 6A), we wondered if the relapse is linked to the change of TRIM24 chromatin occupancy, as previously observed in LNCaP cells compared to LNCaP-abl cells^7^. Indeed, the global chromatin occupancy increased in the relapse tumor and the TRIM24 peaks showed little overlap with the sites observed before castration (Fig. 6D, E, Supp. 16D & Table S6), indicating also an extensive rewiring of the TRIM24 genome binding related to lineage plasticity. Next, we investigated the transcription factor DNA binding motifs presents in the TRIM24 chromatin immunoprecipitation followed by sequencing (ChIP-seq) data. The motifs that were enriched before castration were related to transcription factors associated with AR signaling (e.g., ARE, FOXA1, HOXB13), as expected (Fig. 6F). In contrast, the top enriched motifs observed during the relapse was linked to transcription factors intimately linked to EMT, such as STAT4-6 and SMAD2/3 (Fig. 6F and Table S7). Thus, the data may point towards a critical involvement of TRIM24 in the onset of EMT-associated markers and the adaptation process to SCL-prostate cancer in this model as well. To test this hypothesis, we administered dTRIM24-1 and −2 in LuCaP147 xenograft models which effectively degraded TRIM24 in vivo (Fig. 7A & Supp Fig. 16E). To test the antitumor activity of dTRIM24-1 and −2, we treated LuCaP147 for 3 weeks after castration and monitored tumor regrowth after regression. The treatment with dTRIM24-1 and −2 effectively prevented tumor relapse (Fig. 7A). To increase our comprehension of the gene expression changes underlying residual disease, we performed scRNA-seq (Supp. Figs. 17). As expected, the clusters identified in the residual disease significantly differed from those observed before castration and showed a loss of AR and mTOR signaling (Supp. Figs. 18A, B). Interestingly, in the residual disease following castration, we observed that cluster 3 showed enrichment for EMT and activation of signaling pathways associated with disease progression, including TNFA-NFKB and IL6-JAK-STAT3 (Fig. 7B, C & Supp. Fig. 18C). The treatment with dTRIM24-1 led to a significant reduction in EMT and IL6-JAK-STAT3 signaling as well as a harsh downmodulation of AR signaling specifically within cluster 3 (Fig. 7B, C). To determine if thTRIM24 degradation may lead to more stable epigenetic changes, we assessed H3K27 acetylation(H3K27ac), a canonical marker for open chromatin, before castration and three weeks after we had stopped the treatment with vehicle and dTRIM24-1. Tumors exposed to dTRIM24-1 displayed a significant reduction of H3K27ac peaks as compared to tumors before and after castration (Fig. 7D). Consistently, global H3K27ac and TRIM24-co-occupied H3K27ac levels dropped considerably post dTRIM24-1 as well (Fig. 7E & Supp. Fig. 18D). When we investigated the transcription factors binding sites of AR and EMT, we found a reduction of chromatin opening for both, suggesting that related changes observed in residual disease remain post treatment (Fig. 7B, C, F & Supp. Fig. 18A, B). In line with this continued suppression of tumor growth post dTRIM24-1, gene set enrichment analysis of genes with differential H3K27ac marks revealed a suppression of EMT related pathways (e.g., TGFbeta, TNFalpha-NFkB), AR signaling (e.g., AR response, mTOR), and cell cycle (e.g., E2F targets, G2M checkpoint, mitotic spindle) (Fig. 7G & Table S7), while immunoblotting confirmed reduced levels of N-cadherin and phosphorylated STAT3 (Supp. Fig. 18E). Finally, both dTRIM24-1/2 further inhibited LuCaP147-CR tumor growth when retransplanted into new NRG mice (Supp. Fig. 18F), indicating persistent anti-tumor effect linked to chromatin rewiring during the development of lineage plasticity in this model.

**Figure 6.**
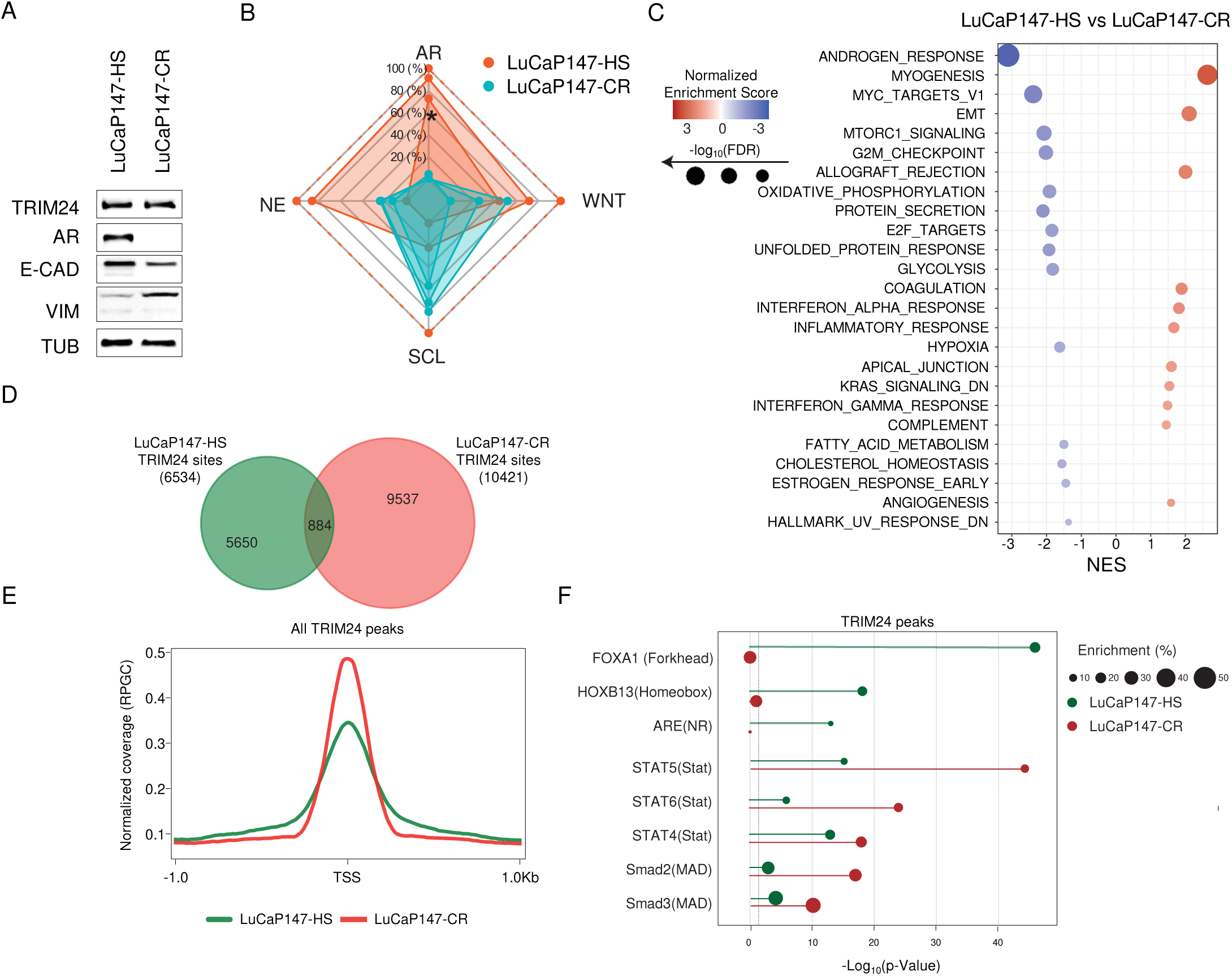
Characterization of the LuCaP147 Stem-Cell-Like lineage plasticity model. **A** Immunoblot analysis of the indicated proteins in LuCaP147 xenografts of the hormone-sensitive (-HS) and the castration-resistant relapse (-CR). **B** Radar plot displaying the gene set enrichment scores of AR-positive (CRPC-AR), neuroendocrine (CRPC-NE), WNT-driven (CRPC-WNT), and stem cell-like subtype (CRPC-SCL) on bulk RNAseq of LuCaP147-HS and LuCaP147-CR xenografts. **C** Gene set enrichment analysis (GSEA) of Hallmark signatures on bulk RNA-seq data of LuCaP147-HS and LuCaP147-CR xenografts. The dot plot illustrates the gene-enriched scores’ false discovery rate (FDR) of the comparison. **D** A Venn diagram showing genomic binding sites and their overlap for TRIM24 chromatin occupancy as determined by ChIP-seq in LuCaP147-HS and LuCaP147-CR. The total number of genomic binding sites for each sample is indicated in parentheses**. E** TRIM24 ChIP-seq signals (derived from the Venn diagram in 6D), are mapped on genome TSS, comparing LuCaP147-HS (green) and LuCaP147−CR (red) samples. **F** A lollipop plot displays the enriched transcription factor DNA binding sites in the TRIM24 ChIP-seq signaling of LuCaP147-HS (green) and LuCaP147-CR (red) tumors. The length of each bar represents - Log10(p-value) for the indicated DNA binding motif, while the size of the circle at the tip indicates the fold enrichment.

**Figure 7.**
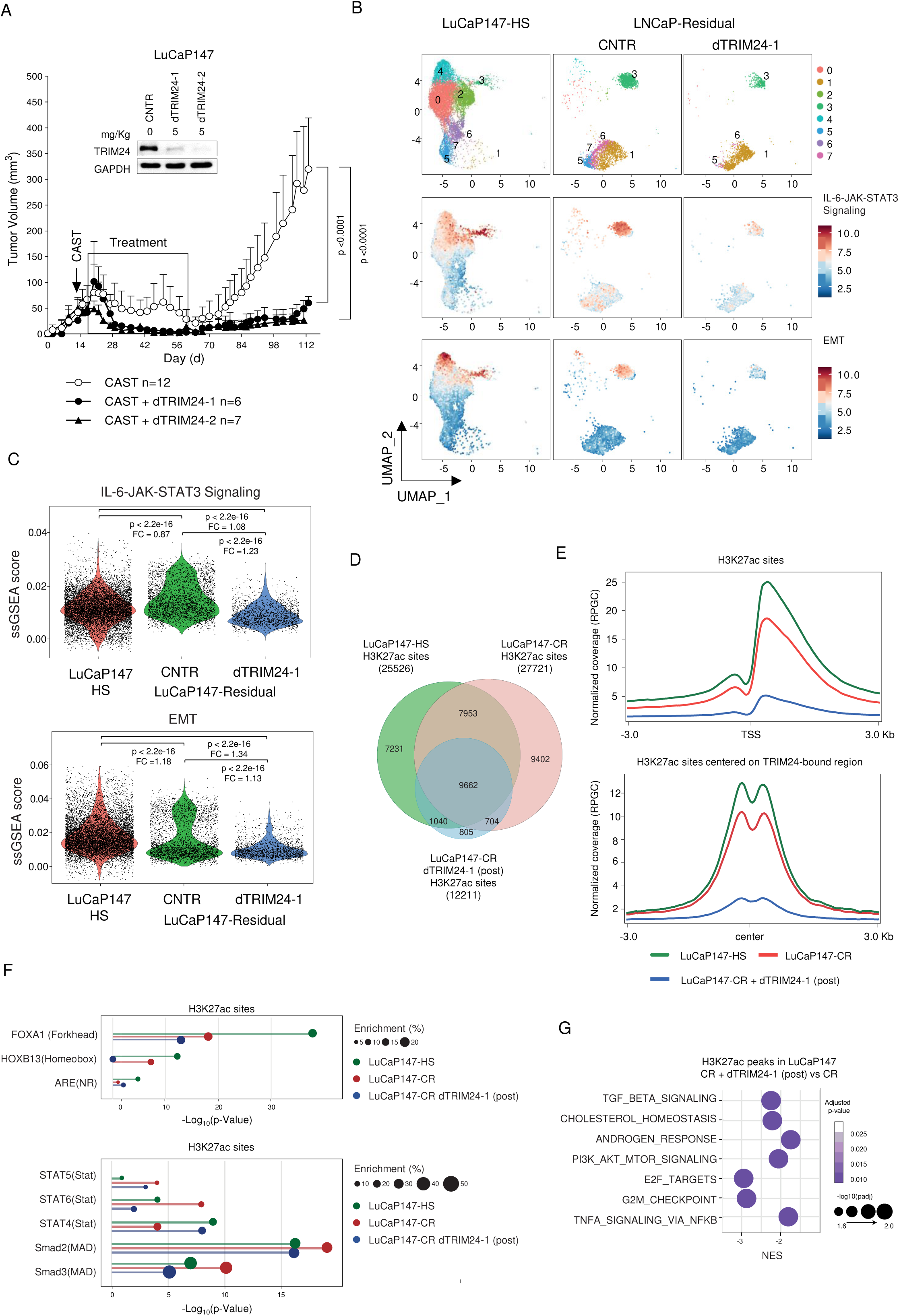
TRIM24 degradation prevents tumor relapse after castration in the LuCaP147 Stem-Cell-Like lineage plasticity model. A Upper panel, Immunoblot analysis of the indicated proteins showing TRIM24 reduction in LuCaP147 xenografts following daily intraperitoneal administration of dTRIM24-1 and dTRIM24-2 at 5mg/Kg for five days. Lower panel, growth kinetics of LuCaP147 xenografts in NRG mice after castration and treatment with either a vehicle (CNTR, white circle), dTRIM24-1 (black circle), or dTRIM24-2 (black triangle). Castration (indicated by arrow) is performed when the tumor reaches an average size of 50-75 mm3. The treatment was administered intraperitoneally for three weeks (indicated by a black rectangle), five days a week, at a 5 mg/kg dosage for both dTRIM24-1 and dTRIM24-2. Statistical significance of the results was assessed using a two-way ANOVA test. **B** UMAP plot of scRNA-Seq data showing the distribution of cell clusters (upper panel) alongside gene-set enrichment scores for the IL6-JAK-STAT3 signaling (middle panel) and EMT signature (low panel) in LuCaP147-HS tumors (left) compared with LuCaP147-Residual tumors treated with either vehicle (CNTR, middle) or dTRIM24-1 (right), sampled after castration at the end of the treatment. **C** Violin plots display ssGSEA scores for IL6-JAK--STAT3 signaling and EMT in the indicated samples. Differences between LUCaP147-HS, LUCaP147-Residual CNTR, and dTRIM24-1 were assessed using the Wilcoxon test. **D** The Venn diagram overlapping the genomic binding sites of H3K27ac across three samples: LuCaP147-HS (green), LuCaP147-CR (red), and LuCaP-CR three weeks post-dTRIM24-1 treatment (blue). The total number of genomic binding sites for each sample is indicated in parentheses. **E** Average H3K27ac ChIP-seq signals (derived from the Venn diagram in 7D) are mapped onto genome TSS (upper panel) and TRIM24-binding region (lower panel), comparing LuCaP147-HS (green), LuCaP147-CR (red), and LuCaP-CR post dTRIM24-1 treatment (blue) samples. **F** A lollipop plot displays the enriched transcription factor DNA binding sites in the H3K27ac ChIP-seq signaling of tumors from LuCaP147-HS (green), LuCaP147-CR (red), and LuCaP147-CR three weeks post dTRIM24-1 treatment (blue). The length of each bar represents - log10(p-value) for the indicated DNA binding motif, while the size of the circle at the tip indicates the fold enrichment. **G** Gene set enrichment analysis (GSEA) of Hallmark signatures using H3K27ac Chipseq signaling compared LuCaP147−CR and LUCaP147-CR dTRIM24-1 (post) samples. The dot plot illustrates the gene-enriched scores’ false discovery rate (FDR) for the comparisons.

## Discussion

Understanding residual disease upon therapeutic interventions is critical to developing more effective combination therapies that offer a prospect of cure or, at the least, more durable disease control. In prostate cancer, the dynamics of adaptation to inhibition of the AR signaling axis remain poorly understood. Here, we identify TRIM24 as a critical core component of this adaptation machinery. In line with this, TRIM24 loss-of-function is sufficient to abolish or at least substantially delay disease progression in multiple disease-relevant models linked to either the restoration of AR signaling or the loss of this lineage-specific oncogenic pathway through reprogramming.

In LNCaP xenograft models where tumor relapse was mainly driven by restoration of AR signaling, we observed in cancer cell subpopulations primed for tumor relapse an upregulation of AR and TRIM24 in agreement with our previously established role for TRIM24 in activating AR signaling in CRPC where androgens are limited^7^. In addition, we establish here that the engagement of AR signaling in CRPC is critically dependent on TRIM24 in different settings involving prevalent modifications of AR, such as point mutations that open the spectrum of possible ligands or the expression of AR-V7 that lacks the ligand binding domain^11,41,46^.

In all models tested, AR inhibition triggered changes in EMT-associated markers and related signaling pathways such as TNFA-NFKB, IL6-JAK-STAT3, and TGFβ, which subsequently increased activity with disease progression, in line with previous publications^17,25,47^. Mechanistically, we found that short-term androgen deprivation and ENZA directly enhanced the interaction of TRIM24 and STAT3, previously shown in glioblastoma^36^. While STAT3 activation was seen in multiple models (e.g., LNCaP, VCaP, LAPC4), it is conceivable that TRIM24’s pleotropic effects on other transcription factors related to EMT and dedifferentiation may also apply in our setting^48–53^. For example, TRIM24 occupancy increased in DNA binding motifs related to STAT4-6 and SMAD2/3 in the castration-resistant LuCaP-147 xenograft model. Nevertheless, in each case the pharmacological degradation of TRIM24 efficiently blocked the engagement of EMT-related pathways mentioned above under AR suppression. While pharmacological degradation of TRIM24 substantially delayed disease progression in LNCaP-derived models, it had a particularly profound effect on LuCaP147 xenografts^54^. Even upon treatment cessation, TRIM24 protein abundance remained low which precluded us from further characterizing TRIM24 chromatin occupancy post treatment. Notably, we observed a striking closure of chromatin as evidence by H3K27ac. Conceivably, the exceptional responses may partly be explained by a recurrent point mutation in SPOP, as we have previously shown that these early driver mutations consistently impair TRIM24 protein degradation^7,55,56^. Thus, the current data may be the first indication that TRIM24 stabilization represents a key Achilles heel in this subtype of prostate cancer. Interestingly, we noticed that resistance to TRIM24 degradation is paralleled by the upregulation of the target itself in LNCaP xenografts. Given that TRIM24 degraders used in this study is not as effective in eliminating tumor subpopulations primed for disease relapse as the knockdown by short hairpin RNAs, the improvement of the current degraders or the development of protein glues for targeted degradation may substantially improve therapeutic outcomes. Given the importance of TRIM24 for disease progression in a growing number of other cancer types, such next-generation TRIM24 degraders may not be limited to the treatment of prostate cancer^48, 50–53, 57–60^. Noteworthy in this regard, TRIM24 is an activator for estrogen receptor signaling and is involved in changes of EMT-associated markers in breast cancer as well^27,61^. In the current study in immune-compromised mice, we did not observe obvious toxic side effects (e.g., weight loss, changes in behavior, or loss of animals) of TRIM24 degraders even if dTRIM24-2 efficiently degraded Trim24 in the murine host in various organs. In agreement, Trim24-knockout mice are viable and fertile^62^. Nevertheless, given multiple reports indicating a function of TRIM24 in immune cells^63–65^, we cannot exclude possible side effects involving the innate or adaptive immune system, which will need to be assessed when further developing pharmacologic degradation of TRIM24 in a clinical setting.

In summary, we provide a blueprint for exploiting TRIM24 as an effective drug target in prostate cancer by its ability to inhibit AR signaling in various settings and directly counteract STAT3 activation and changes in EMT-associated markers involved in resistance development.

## Material and Methods

### Plasmids

The pLKO-SCR (SHCC002), the pLKO-shTRIM24-1 (TRCN0000195528), and the PLK shTRIM24-2 (TRCN0000021259) were purchased from Sigma-Aldrich. The pLV-Bsd-EF1A HA (control) and pLV-Bsd-EF1A-TRIM24-HA lentiviral vectors were obtained from VectorBuilder.

### Synthesis of Bifunctional TRIM24 Degraders

One gram of the bifunctional degrader TRIM24-1 and TRIM24-2 (see Supp. Fig. 3A) has been synthesized by ChemPartner based on the previously published structure for TRIM24-1^26^ The synthesis for dTRIM24-2 is detailed in Patent No. US 10,702,504 B2 (see Supplementary Methods). In the in vivo studies, the dTRIM24-1 and dTRIM24-2 have been resuspended in 5% DMSO and used at the indicated doses.

### Cell Lines

LNCaP, LAPC4, VCaP, 22RV1, and HEK 293T cell lines were purchased from ATCC (Manassas, USA). The LNCaP P53-/-; RB1-/- cell line was a gift from Peter Nelson (Fred Hutch Cancer Center, Seattle). In each case, only early passages (up to passage five) have been used for the experiments. The supernatant of all cell lines was routinely tested once per month) using the MycoAlertTM Mycoplasma Detection Kit (Catalog #: LT07-318 Lonza). All cell lines resulted negative for Mycoplasma infection.

### Cell Culture

LNCaP, the LNCaP P53-/-; RB1-/-, and the 22Rv1 cell lines were cultured RPMI 1640 medium (Cat 21875-24, Gibco) supplemented with 10% Fetal Bovine Serum (FBS-11A Capricorn Scientific) and 1% Penicillin/Streptomycin (15140-122 Life Technologies) with 5% CO2 at 37°C. The LAPC4 cell line was cultured in RPMI 1640 medium supplemented with 10% Fetal Bovine Serum, 1nM DHT (5α-Dihydrotestosterone, D-073, Sigma-Alderich), and 1% Penicillin/Streptomycin, with 5% CO2 at 37 °C. The LNCaP-95 cell line was cultured in Phenol-Red Free RPMI 1640 medium (11835-063 Gibo) containing 10% charcoal-stripped serum (CSS, Serum, Charcoal Stripped; Cat 12676-029 Gibco) and 1% Penicillin/Streptomycin with 5% CO2 at 37°C. The HEK 293T and the VCaP cell line were cultured in DMEM medium (Cat 2688-133, Gibco) supplemented with 10% FBS and 1% µg/ml Penicillin/Streptomycin with 5% CO2 at 37°C.

### PEI-mediated Transfection and Lentiviral Infection

The HEK 293T cells were transfected with a vector for each shRNA, packaging, and envelope vector using the PolyEthylenImine (PEI) method. The HEK 293T cells were seeded in a 100 mm culture dish (4×10^6 cells/plate) and incubated overnight at 37°C in a 5% CO2 humidified atmosphere. After 24 hours, the vector plasmid (3 μg), packaging plasmid (pCMV-dR8.2; 2.7 μg), and envelope plasmid (pVSV-G; 0.7 μg) were mixed in Opti-MEM culture medium (300μL/10 cm culture dish; Opti-MEM™ I Reduced Serum MediumCat 31985-070, Gibco) and with 1.25 mM PEI (Cat 919012, Sigma-Aldrich) solution (ratio μL PEI: μg DNA 4:1). The DNA/PEI mixture were incubated for 15 minutes at room temperature and added to the HEK 293T cell supernatant. After 48 hours of transfection, the viral supernatants were collected and filtered through a 0.45μm filter. The LNCaP, LNCaP-95, and 22RV1 cell lines were incubated with viral supernatant and 8 µg/ml of Polybrene (H9268, Sigma-Aldrich) for 72 hours. After incubation, the selection was performed using a specific antibiotic for a minimum of 10 days: 2 µg/ml of puromycin (P8833, Sigma-Aldrich) for cells infected with the PLKO vectors or 10 µg/ml of blasticidin (15205, Sigma-Aldrich) cells infected with the PLV vectors.

### Tumor Tissue Enzymatic Digestion

We followed a specific procedure to obtain a single-cell suspension of tumor cells from patient-derived xenograft (PDX) or xenografted tumor tissue. First, we cut the tumor tissue into small pieces (1-0.5 mm) using a scalpel blade. Then, we digested the tissue for 45-60 minutes in a DMEM/F-12, GlutaMAX™ medium (Cat 31331-028, Gibco) with Collagenase Type I (200U/ml; Cat 17018-029, Gibco) and DNAseI (1mg/ml; Cat 11284932001, Roche) at 37°C. After enzymatic dissociation, we filtered the cell suspension through a 100 µM cell strainer (Roche) to eliminate macroscopic tissue pieces and then centrifuged it. The resulting cell pellet was resuspended in a 2-volume RBC lysis buffer (Roche) and incubated for 3 minutes at room temperature. After centrifugation, the cells were resuspended in a PBS solution. We used this single-cell suspension to inject animals for pharmacological studies mentioned in the animal section.

### Colony Formation Assay

The LNCaP, the LAPC4, 22RV1, and LNCaP-95 parental cell lines were seeded in duplicate into 12-well plates at a density of 12500 cells per well. Before the seeding the 12-well plates were pre-coated with a solution of poly-D lysine (120 µg/mL in PBS) for 24 hours at 37°C. All cell lines were cultured in a testosterone-depleted medium (charcoal-stripped serum, CSS) supplemented with the following conditions: (i) 1 nM DHT alone or in combination with 5 µM of dTRIM24-1 or 5 µM of dTRIM24-2; (ii) without DHT, either alone or in combination with 5 µM of dTRIM24-1 or 5 µM of dTRIM24-2; (iii) without DHT, supplemented with 10 µM of enzalutamide alone or in combination with 5 µM of dTRIM24-1 or 5 µM of dTRIM24-2. The medium was refreshed 3 times a week. After 18-20 days, colony confluence was assessed using the IncuCyte live-cell analysis system. The LNCaP, 22RV1, and LNCaP-95 cells infected with control, shTRIM24-1, and shTRIM24-2 vectors were examined under the same conditions. The experiments with VCaP cells were performed using 1 µM of dTRIM24-1 or 1 µM of dTRIM24-2 instead of 5 µM and maintained a minimum concentration of 0.01 nM DHT in a testosterone-depleted condition. Each cell line and condition were tested in biological triplicate.

### Transwell Assay

LNCaP cells were cultured in a testosterone-depleted medium (charcoal-stripped serum – CSS) with either 10uM of enzalutamide (915087-33-1 Angene) alone or 5uM of dTRIM24 for 3 days. The LNCaP cells cultured in CSS supplemented with 1nM DHT (D-073 Sigma-Alderich) were used as a control group. After 3 days, 5 x 10^4 serum-starved cells were added to the upper compartment of the Boyden chamber, and the lower compartment was filled with RPMI medium containing 10% FBS. After 24 hours of incubation, the percentage of invaded cells was measured. The migrated cells were stained with a 2% crystal violet staining solution and visualized under a microscope.

### Incucyte® Scratch Wound Assay

A wound healing assay was conducted on LNCaP cells line and LNCaP cells infected with control, shTRIM4-1, and shTRIM24-2 pLKO vectors using live cell imaging on the IncuCyte® system. The cells were cultured in a 96-well plate with different medium conditions. Specifically, LNCaP cells at a density of 50000 cells/well were maintained in a testosterone-depleted medium (charcoal-stripped serum - CSS) supplemented with either 10uM of enzalutamide alone or 5uM of dTRIM24. The control group consisted of LNCaP cells cultured in CSS supplemented with 1nM DHT. After three days of culture, the feeder layer of cells was scratched, and then the cells were tested for 120 hours. The IncuCyte® software was used to measure the spatial cell density in the wound area relative to the spatial cell density outside the wound.

### Immunoblotting

The primary antibodies used are anti-GAPDH (sc-47724, Santa Cruz Biotechnology), anti-Tubulin (3873, Cell Signaling Technology), anti-TRIM28 (4123, Cell Signaling Technology), anti-AR (133273, Abcam), anti-NKX3.1 (87300, Cell Signaling Technology), anti-TRIM24 (14208-1-AP Proteintech), anti-E-Caderin (3195, Cell Signaling Technology), anti-Vimentin (5741, Cell Signaling Technology), anti-N-cadherin (13116, Cell Signaling Technology), anti-STAT3 (8019, Cell Signaling Technology), and anti-Phospho-STAT3(Ser727) (9134, Cell Signaling Technology). Snap-frozen tumor tissue (Fragments of 25-30 mg) or cellular pellet was lysed using RIPA Buffer supplemented with a cocktail of phosphatase inhibitors (4906845001 Roche) and protease inhibitors (5892953001 Roche). The protein concentration was determined using a BCA reagent (Cat A52255, Thermo Fisher Scientific). 30-50 μg of whole protein lysate was separated on 8-12% SDS–polyacrylamide gels and transferred onto PVDF membrane (88518 Thermo Fisher Scientific). The membranes were first blocked with 5% milk in Tris Buffered Saline with Tween 20 (TBST) for 30 minutes at room temperature. Only for the anti-Phospho-STAT3 antibody the blocking was made in 5% BSA. After that, they were incubated with primary antibodies overnight at 4°C. Then, they were incubated with secondary antibodies (anti-rabbit IgG HRP W401B and anti-mouse IgG HRP W402B Promega) for 1 hour at room temperature. The protein bands were visualized using the western bright quantum reagent (K-12042-D20 Advansta) and quantified using the Fusion Solo IV LBR system.

### Immunoprecipitation

Immunoprecipitation (IP) analysis was performed on LNcaP cells that were cultured in CSS medium for three days. The cells were treated with either DHT 1nM or Enzalutamide 10microM. To lysate the cellular pellet, the IP Buffer was used, which contained 20mM Tris-HCl (pH 7.5), 150mM NaCl, 1mM EDTA, 2mM Na3VO4, 5mM NaF, and 1% Triton X-100, along with protease and phosphatase inhibitor cocktails. Overnight incubation of anti-TRIM24 (3 micrograms) antibodies with 1mg of cell lysates was done. Anti-IgG was utilized as a control. Dynabeads Protein A (Cat 10001D, Invitrogen) was used to immunoprecipitated the samples. The immunoprecipitants were washed thrice with the IP buffer. The immunoprecipitated were resolved in a 2x SDS lysis buffer and analyzed in an SDS-PAGE gel.

### RT-qPCR analysis

RNA was extracted from a cellular pellet of LNCaP cells using an RNeasy kit from Qiagen (74104). The manufacturer’s guidelines were followed for the extraction, which was performed at the specified culture condition. According to the manufacturer’s protocol, RT-qPCR analysis was carried out using KAPA SYBR® FAST One-Step (Cat KK4600, Sigma-Aldrich). The primer sequences used in the analysis were obtained from PrimerBank, and a list of the primers can be found in the Supplementary Table 5. The housekeeping gene used in the qPCR analysis was Actin. The 2-ΔΔCt method was employed in the qPCR analysis.

### RNA Extraction for RNA-seq analysis

RNA was extracted from snap-frozen tumor fragments (25-30 mg) of cellular pellets using the RNeasy kit from Qiagen (74104) as per the manufacturer’s instructions. The extracted RNAs were processed using the NEB Next Ultra II Directional Library Prep Kit for Illumina and then sequenced on the Illumina NextSeq500 with single-end, 75-base pair-long reads for RNA-seq analysis.

### ChIP-seq

ChIP-seq analysis for TRIM24 and H3K27ac was conducted on LUCAP147 tumors under different growth conditions: LuCaP147-HS, LuCaP147-CR, and LUCAP147-CR post-treated with dTRIM24-1. The ChIP-seq was performed in duplicate. For each replicate, two tumors from LuCaP147-HS and LuCaP147-CR (both averaging 75 mm³), along with five tumors from LuCaP147-CR post-treated with dTRIM24 (averaging 30 mm²), were pooled. A corresponding of 20 mg of pooled tumor tissue was used for each immunoprecipitation. Briefly, the freshly excised LuCaP147 tumors were crushed and resuspended in GentleMACS C tubes (130-093-237, Miltenyi Biotec) using a PBS solution supplemented with protease and phosphatase inhibitors. The tumor fragments were then dissociated into small cell clusters using a gentleMACS™ tissue dissociator (130-134-029, GentleMACS™ OCTO dissociator, Miltenyi Biotec). The resulting cell suspension was filtered through a 100 µm cell strainer (431752, Corning) to remove macroscopic tissue debris. After centrifugation, the cells were fixed in 1% formaldehyde in PBS at room temperature for 10 minutes and subsequently quenched. ChIP-seq was performed using the iDeal ChIP-seq kit for Histones (C01010051, Diagenode SA) following the manufacturer’s instructions. The chromatin was sonicated at high power, applying 30 cycles of 30 seconds on and 30 seconds off using a Bioruptor Plus (Biosense). For each immunoprecipitation, either 1 μg of anti-H3K27ac antibody (document number C15410196, Diagenode SA) or 3 μg of anti-TRIM24 antibody (reference 14208-1-AP, Proteintech) was used. The genomic DNA was processed using the MicroPlex Library Preparation Kit v3 (document number C05010001, Diagenode SA) and sequenced on an Illumina NextSeq 2000 platform, producing single end reads that were 120 base pairs in length.

### Single-cell isolation for scRNA-seq

For scRNA-seq analysis, we isolated individual cells from freshly obtained tumor fragments that were already dissociated (see Ex vivo culture of PDX section for more information on the dissociation process). The isolated single cells were suspended in PBS and loaded (10,000-5,000 cells) into the 10x Chromium Controller at 10x Genomics in Pleasanton, CA, USA. We used the Chromium Next GEM Single Cell 3’ v3.1 reagent kit from 10x Genomics (PN-1000121) and followed the manufacturer’s instructions. The scRNA-Seq analysis was conducted as a single replicate using pooled biological replicate tumor samples. For the LNCaP-Residual tumors, the TRIM24-KD1/2 and control groups consisted of tumors pooled three weeks post-castration, when their average size was 15-20 mm³. In the case of the LNCaP-HS and LNCaP-CR tumors, which were post-treated with either vehicle or dTRIM24, three tumors were pooled when their average size was 75-100 mm³. For the LNCaP-Residual tumors that were treated with vehicle and dTRIM24-1, five tumors were pooled three weeks post-castration at the end of the treatment, with their average size being 15-20 mm³. Similarly, for the LuCaP147-Residual tumors treated with vehicle and dTRIM24-1, five tumors were pooled three weeks post-castration at the end of the treatment, with an average size of 25-30 mm³.

### Animal Experiments

All experiments involving animals were conducted following a protocol approved by the Swiss Veterinary Authority/Board (TI-40-2018, TI 41-18, TI-44-2019 TI30-21 TI-49-2023) and received approval from the ethical committee of the Institute of Oncology Research (IOR). All in vivo studies used male NRG (NOD-Rag1null IL2rgnull, NOD rag gamma) mice between 6-8 weeks old.

### Management and generation of Patient-Derived Xenograft (PDX) line

We use the LuCaP-147 PDX model. This line was maintained in vivo by transplanting Matrigel-embedded tissue tumor fragments into male NRG mice. To create LuCaP147-CR sublines, we harvested tissue fragments from a LuCaP-147 (that regrew after castration. This CR LuCaP147 model was then passed on to other castrated male mice and designated as CR after being passed on at least three times.

### In Vivo Pharmacology Studies

For our in vivo pharmacology studies, we utilized LNCaP and LNCaP P53-/-; RB1-/- cell lines and freshly isolated single-cell suspensions from the LuCaP147 tumor fragments. The tumor single cells were mixed with PBS and 50% Matrigel and injected subcutaneously into the dorsal flanks of the mice. For LNCaP and LNCaP P53-/-; RB1-/- cells, 2.5 x 10^6 cells per mouse were injected for LuCaP-147, 1 x 10^6 cells per mouse were injected. Tumor growth measured using a digital caliper and calculated the tumor volumes using the formula (L x W^2) /2, where L is the length, and W is the width of the tumor. Tumor volume was measured twice a week. After the tumor size reached 50-75 mm^3, we surgically castrated the mice. Then, we administered the treatment intraperitoneally for three weeks, five days per week. For LNCaP and LNCaP P53-/-; RB1-/- cell lines, we used 10 mg/kg of dTRIM24-1-2 and 30 mg/kg of Enzalutamide. For the LuCaP-147 PDX line, we used 5 mg/kg of dTRIM24-1-2. After the experiment, we euthanized the mice and removed the tumors for molecular assessment. For the studies in castrated NRG mice, we used LNCaP95 cells or freshly isolated single-cell suspensions from the LuCaP 147CR tumors. To initiate the study, we injected 1 x 10^6 cells/mouse for LNCaP95 or 1 x 10^6 cells/mouse for LuCaP 147 in PBS and 50% Matrigel subcutaneously into the dorsal flanks of the mice. Once the tumor size reached 25-50 mm^3, we began the drug treatment. We administrated dTRIM24 at a dose of 10 mg/kg and Enzalutamide at a dose of 30 mg/kg intraperitoneally once daily for two weeks. For the LuCaP-147 PDX, we used 5 mg/kg of dTRIM24 intraperitoneally once daily for three weeks, five day a week. After the completion of the treatment, we euthanized the mice and removed the tumors for molecular assessment.

## Statistical analysis

The statistical analysis was performed using GraphPad Prism. The results present the means ± standard errors. Student t-tests were used to identify differences between the two groups. For multiple groups, one-way analysis of variance (ANOVA) tests was used. The statistical analyses were based on data collected from at least three independent experiments. Statistically significant differences were indicated by P<0.05.

## Data availability

The bulk RNA-seq and single-cell RNA-seq data generated in this study have been deposited in the Gene Expression Omnibus (GEO) under accession code GSE267347. The ChIP-seq data for TRIM24 and H3K27ac are also included within this GEO submission. Whole-exome sequencing data have been submitted to the Sequence Read Archive (SRA) under BioProject PRJNA1213694 with BioSample accessions SAMN46343533, SAMN46343534, SAMN46343535, and SAMN46343536.

