## Supplemental Figures for "TRIM24 Degradation Counteracts Adaptation to Androgen Receptor Inhibition in Prostate Cancer"

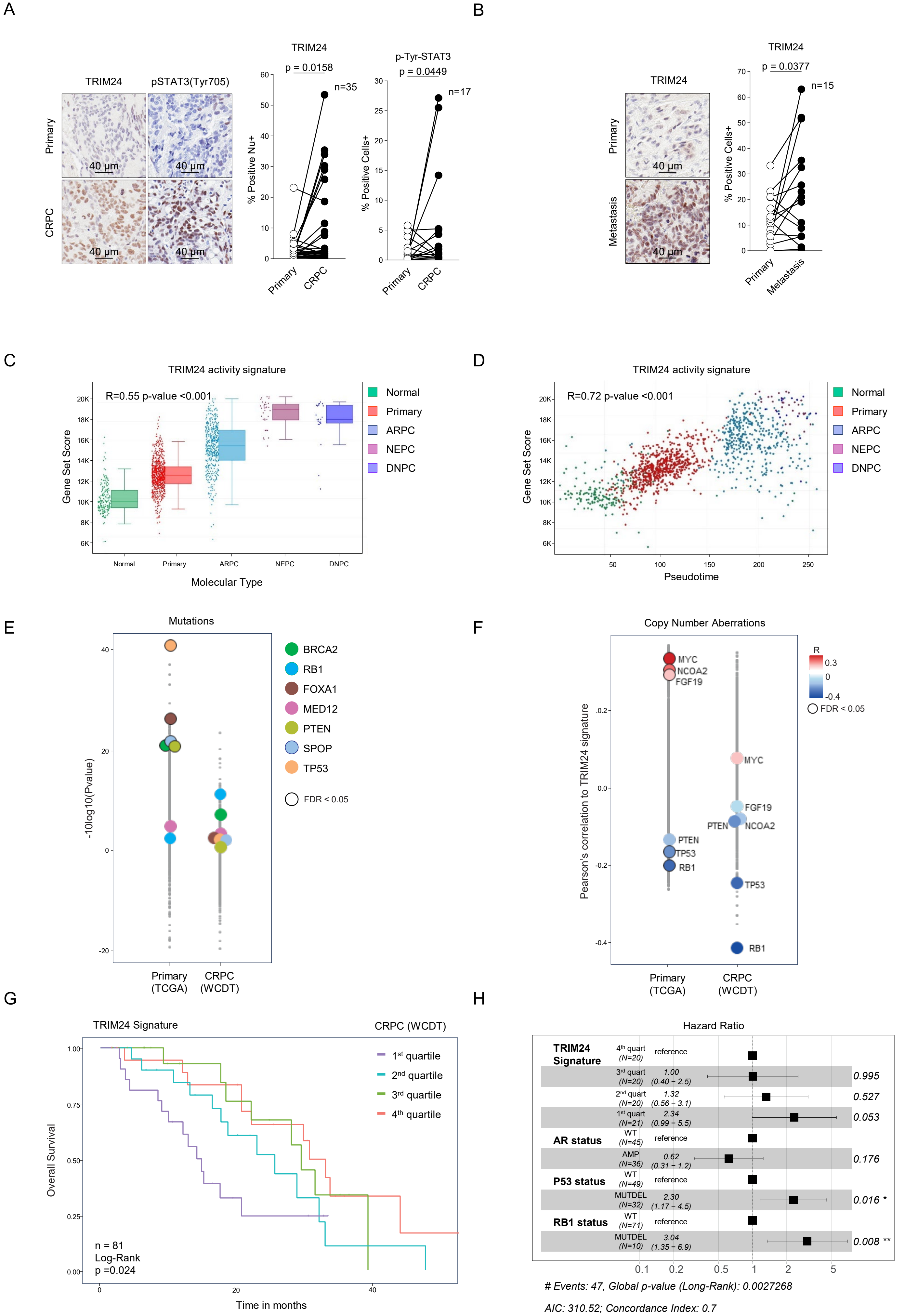
A
TRIM24
pSTAT3(Tyr705)
Primary
CRPC
40 µm
TRIM24
p = 0.0158
n=35
% Positive Nu+
p-Tyr-STAT3
p = 0.0449
n=17
% Positive Cells+
B
TRIM24
p = 0.0377
n=15
Metastasis
C
TRIM24 activity signature
R=0.55 p-value <0.001
Gene Set Score
Molecular Type
Normal
Primary
ARPC
NEPC
DNPC
D
R=0.72 p-value <0.001
Pseudotime
E
Mutations
-10log10(Pvalue)
BRCA2
RB1
FOXA1
MED12
PTEN
SPOP
TP53
FDR < 0.05
Primary (TCGA)
CRPC (WCDT)
F
Copy Number Aberrations
Pearson's correlation to TRIM24 signature
MYC
NCOA2
FGF19
PTEN
TP53
RB1
G
TRIM24 Signature
CRPC (WCDT)
Overall Survival
1st quartile
2nd quartile
3rd quartile
4th quartile
n = 81
Log-Rank
p =0.024
Time in months
H
Hazard Ratio
TRIM24 Signature
4th quart (N=20) reference
3rd quart (N=20) 1.00 (0.40 – 2.5) 0.995
2nd quart (N=20) 1.32 (0.56 – 3.1) 0.527
1st quart (N=21) 2.34 (0.99 – 5.5) 0.053
AR status
WT (N=45) reference
AMP (N=36) 0.62 (0.31 – 1.2) 0.176
P53 status
WT (N=49) reference
MUTDEL (N=32) 2.30 (1.17 – 4.5) 0.016 *
RB1 status
WT (N=71) reference
MUTDEL (N=10) 3.04 (1.35 – 6.9) 0.008 **
### Events: 47, Global p-value (Long-Rank): 0.0027268
AIC: 310.52; Concordance Index: 0.7

Figure S1

**Supplementary Figure 1. TRIM24 is associated with disease progression in human prostate cancers. A** TRIM24 and phosphor-STAT3(Tyr705) immunohistochemistry (IHC) analysis on human tumor tissue microarrays (TMAs) of primary and matched castration-resistant prostate cancer (CRPC). A representative image from one patient is shown on the left (scale bar 40 μm). In the middle, a bar plot quantifies the number of tumor cells exhibiting positive nuclear staining for TRIM24. Each dot represents the quantification of an individual patient's tumor tissue specimen. On the right, a bar plot displays the number of cells exhibiting positive nuclear staining for phospho-STAT3 (Tyr705) in a subgroup of 17 out of 35 human TMA, showing a rise in TRIM24-positive cells from primary to matched CRPC samples. **B** TRIM24 IHC analysis on human tumor TMA of primary and matched prostate cancer metastasis. On the left side, a representative image of a patient (scale bar 40 μm) is showing, and on the right is the quantification of cells with TRIM24-positive nuclear staining. Statistical differences in **A** and **B** were tested using an unpaired Student's t-test. **C** Box plot showing the gene set enrichment score of TRIM24 activity gene (Lv, D., Li, Y., Zhang, W. *et al.* TRIM24 is an oncogenic transcriptional co-activator of STAT3 in glioblastoma. *Nat Commun* 8, 1454, 2017) set across normal tissue (Normal), primary prostate cancer (Primary), AR-positive CRPC (ARPC), neuroendocrine CRPC (NEPC), and double-negative CRPC (DNPC). This analysis is based on a harmonized RNA sequencing atlas (https://prostatecanceratlas.org/app/home; Bolis, M., Bossi, D., Vallerga, A. *et al.* Dynamic prostate cancer transcriptome analysis delineates the trajectory to disease progression. *Nat Commun* 12, 7033, 2021). The centerline of each box represents the median score, while the bottom and top edges denote the first and third quartiles, respectively. Pearson correlation was employed to assess the relationship between TRIM24 activity and disease progression. **D** Corresponding scattered plot depicting the correlation of the enrichment score of the TRIM24 activity gene set and the pseudotime progression score. The correlation was assessed by a one-sample Wilcoxon test. **E** Monodimensional plot of P values associated with Pearson's correlation coefficients expressed in -10* log 10(P value) (FDR-adjusted). Coefficients were determined for the correlation between somatic mutations (0: wild type; 1: non-synonymous mutation) and the single-sample GSEA score computed on TRIM24 signature in primary (TCGA) and CRPC (WCDT; Abida W, Cyrta J, Heller G, et al. Genomic correlates of clinical outcome in advanced prostate cancer. *Proc Natl Acad Sci U S A*. 2019;116(23):11428-11436).) prostate cancer cohorts (see supplementary method section). Only recurrently mutated genes (at least in five individuals) were considered. BRCA2 (green), RB1 (blue), FOXA1 (brown), MED12 (pink), PTEN (lime), SPOP (light blue), and TP53 (orange). **F** Monodimensional plots showing Pearson's correlation coefficients between the single-sample GSEA score for the TRIM24 signature and numeric copy number status (–2 = homozygous deletion, –1 = heterozygous deletion, 0 = wild type, 1 = gain, 2 = amplification).

Data are stratified into primary (TCGA) and metastatic (CRPC/NEPC) prostate tumors. Each point corresponds to a single gene, with the color scale reflecting the sign and magnitude of the correlation (red = positive, blue = negative). **G** Kaplan-Meier curves showing the overall survival (OS) for 81 patients with castration-resistant prostate cancer (CRPC) from the West Coast Dream Team dataset. Patients were divided into quartiles based on the single-sample GSEA score computed on TRIM24 signature (see supplementary methods for details). Statistical differences were assessed using a log-rank test. **H** The forest plot showing the results of a multivariate analysis examining the relationship between TRIM24 signatures, AR status, p53 status, and RB1 status, and their association with overall survival in patients with the WCDT CRPC dataset. Significant factors ($p < 0.05$) are indicated as follows: * p-value < 0.05, ** p-value < 0.01

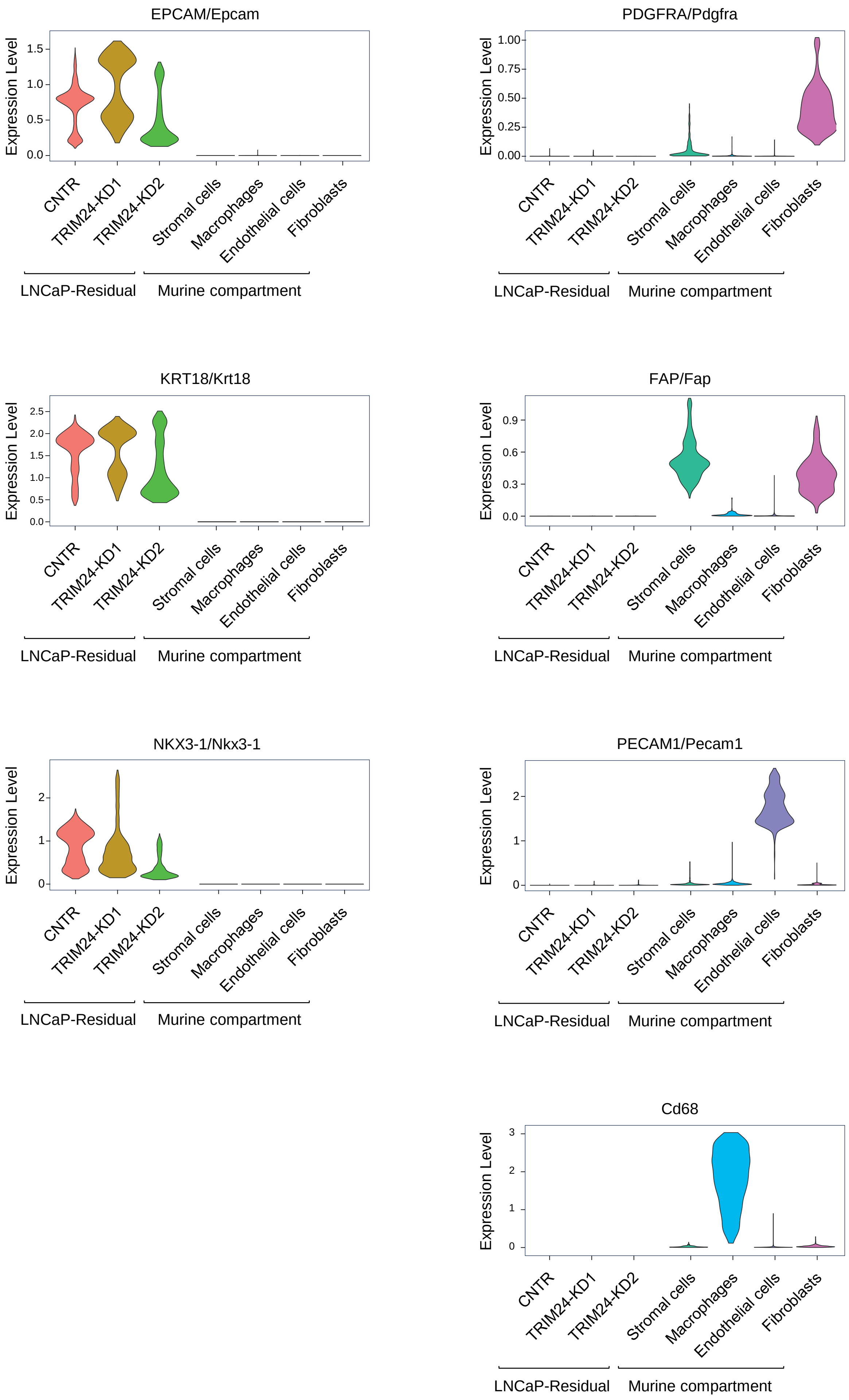
EPCAM/Epcam
PDGFRA/Pdgfra
KRT18/Krt18
FAP/Fap
NKX3-1/Nkx3-1
PECAM1/Pecam1
Cd68
Expression Level
CNTR
TRIM24-KD1
TRIM24-KD2
Stromal cells
Macrophages
Endothelial cells
Fibroblasts
LNCaP-Residual
Murine compartment

Figure S2

**Supplementary Figure S2. Analysis of epithelial and stromal markers of the human and murine compartments of scRNAseq data culled from LNCaP-Residual shSCR (CNTR1), TRIM24KD-1, and TRIM24KD-2 tumors.** Violin plots that illustrate the expression levels of key human prostate tumor markers, such as EPCAM, NKX3-1, and KRT18, alongside murine stromal and immune markers, including FAP, PDGFRA, PECAM1, and CD68.

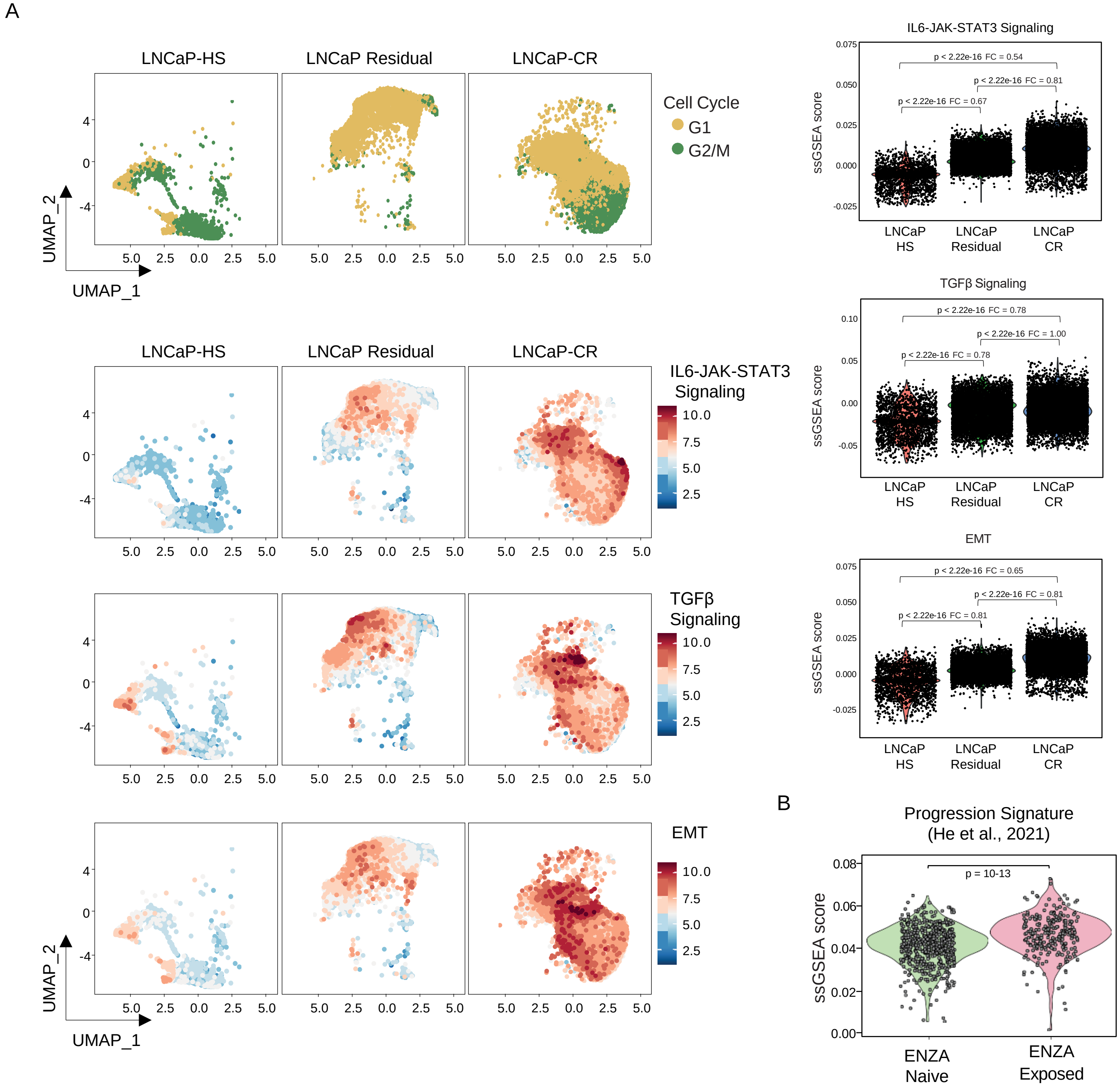
A
LNCaP-HS
LNCaP Residual
LNCaP-CR
Cell Cycle
G1
G2/M
UMAP_2
UMAP_1
IL6-JAK-STAT3 Signaling
TGFβ Signaling
EMT
ssGSEA score
LNCaP HS
LNCaP Residual
LNCaP CR
p < 2.22e-16 FC = 0.54
p < 2.22e-16 FC = 0.81
p < 2.22e-16 FC = 0.67
p < 2.22e-16 FC = 0.78
p < 2.22e-16 FC = 1.00
p < 2.22e-16 FC = 0.78
p < 2.22e-16 FC = 0.65
p < 2.22e-16 FC = 0.81
p < 2.22e-16 FC = 0.81
B
Progression Signature (He et al., 2021)
p = 10-13
ENZA Naive
ENZA Exposed

Figure S3

**Supplementary Figure 3. LNCaP xenograft derived progression signature is associated with ENZA resistance in human single-cell data. A** In the left panels, Uniform Manifold Approximation and Projection (UMAP) analysis of single-cell RNA sequencing (scRNA-Seq) data from LNCaP xenografted tumors, including hormone-sensitive LNCaP(-HS) tumors, LNCaP-Residual tumors sampled three weeks post-castration, and relapsed LNCaP(-CR) castration-resistant tumors. The upper panel presents a UMAP visualization showing the distribution of tumor cells in the G1 (yellow) and G2/M (green) phases. The lower panels display gene set enrichment scores for the indicated hallmark signatures across the samples. In the right panel, the corresponding violin plot displaying the single-sample gene set expression scores of the indicated hallmark signatures across the samples. Median pathway scores for each group are indicated, and fold-change values (the ratio of medians) are annotated between groups. Statistical significance was assessed using the Wilcoxon rank-sum test. **B** Violin plot illustrating the single-sample gene set enrichment score (ssGSEA) for the progression signature generated from LNCaP xenograft models derived from pseudo-bulk scRNA-seq data comparing LNCaP-CR with LNCaP-HS. The top 200 differentially expressed genes (DEGs), selected based on their log2 fold change (log2FC), were utilized to create this signature (see methods). The progression signature was used to interrogate a single-cell RNA sequencing (scRNA-seq) dataset of human tissue specimens obtained from prostate cancer biopsies of patients before and after ENZA treatment**.** The statistical significance of biological triplicates is calculated using the Wilcon-Cox test.

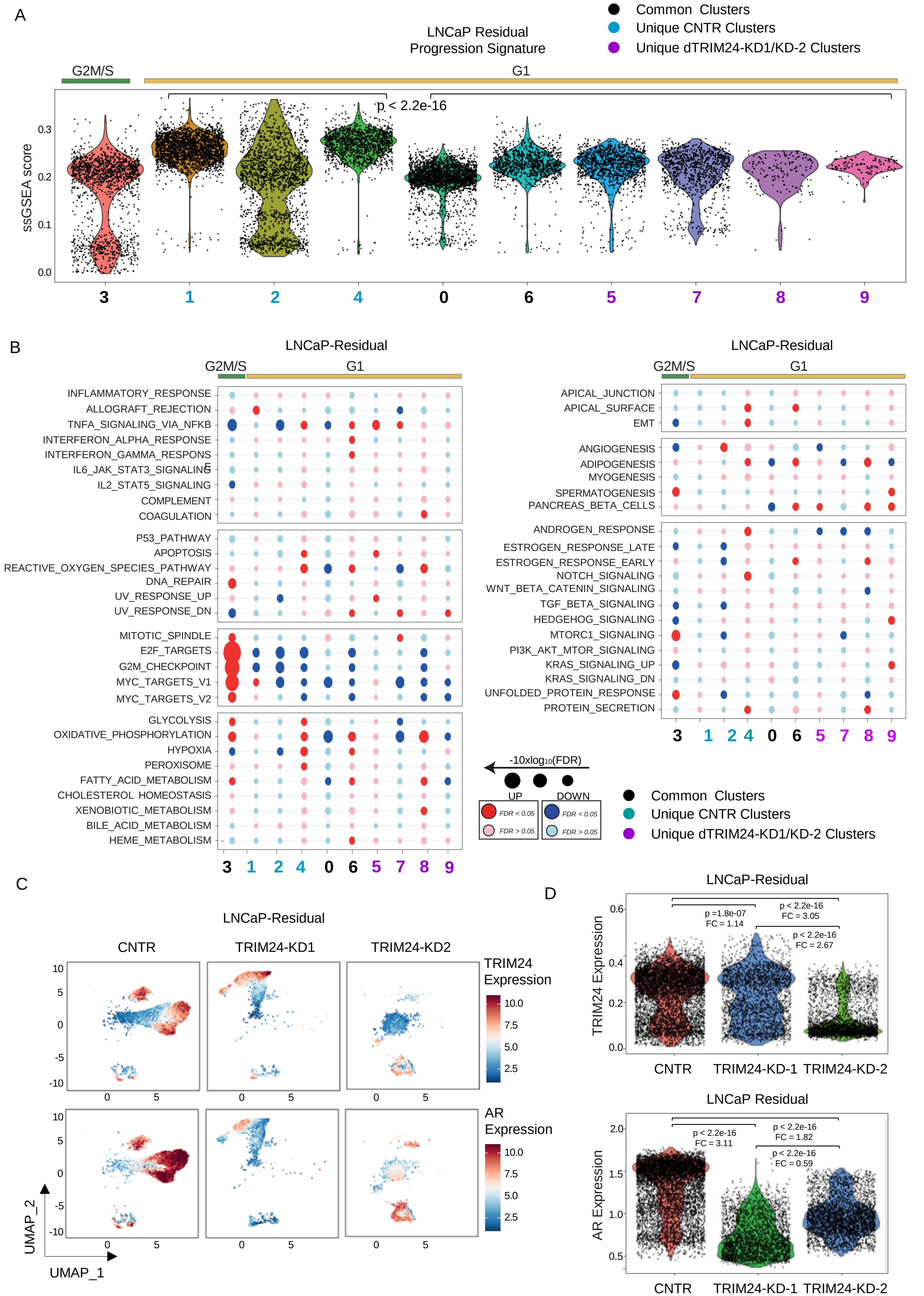
A
LNCaP Residual
Progression Signature
Common Clusters
Unique CNTR Clusters
Unique dTRIM24-KD1/KD-2 Clusters
G2M/S
G1
p < 2.2e-16
ssGSEA score
0.3
0.2
0.1
0.0
3
1
2
4
0
6
5
7
8
9
B
LNCaP-Residual
G2M/S
G1
INFLAMMATORY_RESPONSE
ALLOGRAFT_REJECTION
TNFA_SIGNALING_VIA_NFKB
INTERFERON_ALPHA_RESPONSE
INTERFERON_GAMMA_RESPONSE
IL6_JAK_STAT3_SIGNALING
IL2_STAT5_SIGNALING
COMPLEMENT
COAGULATION
P53_PATHWAY
APOPTOSIS
REACTIVE_OXYGEN_SPECIES_PATHWAY
DNA_REPAIR
UV_RESPONSE_UP
UV_RESPONSE_DN
MITOTIC_SPINDLE
E2F_TARGETS
G2M_CHECKPOINT
MYC_TARGETS_V1
MYC_TARGETS_V2
GLYCOLYSIS
OXIDATIVE_PHOSPHORYLATION
HYPOXIA
PEROXISOME
FATTY_ACID_METABOLISM
CHOLESTEROL_HOMEOSTASIS
XENOBIOTIC_METABOLISM
BILE_ACID_METABOLISM
HEME_METABOLISM
3 1 2 4 0 6 5 7 8 9
LNCaP-Residual
G2M/S
G1
APICAL_JUNCTION
APICAL_SURFACE
EMT
ANGIOGENESIS
ADIPOGENESIS
MYOGENESIS
SPERMATOGENESIS
PANCREAS_BETA_CELLS
ANDROGEN_RESPONSE
ESTROGEN_RESPONSE_LATE
ESTROGEN_RESPONSE_EARLY
NOTCH_SIGNALING
WNT_BETA_CATENIN_SIGNALING
TGF_BETA_SIGNALING
HEDGEHOG_SIGNALING
MTORC1_SIGNALING
PI3K_AKT_MTOR_SIGNALING
KRAS_SIGNALING_UP
KRAS_SIGNALING_DN
UNFOLDED_PROTEIN_RESPONSE
PROTEIN_SECRETION
3 1 2 4 0 6 5 7 8 9
-10xlog10(FDR)
UP
DOWN
FDR < 0.05
FDR > 0.05
FDR < 0.05
FDR > 0.05
Common Clusters
Unique CNTR Clusters
Unique dTRIM24-KD1/KD-2 Clusters
C
LNCaP-Residual
CNTR
TRIM24-KD1
TRIM24-KD2
TRIM24
Expression
10.0
7.5
5.0
2.5
AR
Expression
10.0
7.5
5.0
2.5
10
5
0
-5
-10
0
5
UMAP_2
UMAP_1
D
LNCaP-Residual
TRIM24 Expression
0.6
0.4
0.2
0.0
p =1.8e-07
FC = 1.14
p < 2.2e-16
FC = 3.05
p < 2.2e-16
FC = 2.67
CNTR
TRIM24-KD-1
TRIM24-KD-2
LNCaP Residual
AR Expression
2.0
1.5
1.0
0.5
p < 2.2e-16
FC = 3.11
p < 2.2e-16
FC = 1.82
p < 2.2e-16
FC = 0.59
CNTR
TRIM24-KD-1
TRIM24-KD-2

Figure S4

**Supplementary Figure 4. Pathway analysis related to TRIM24 knockdown of single-cell data related to residual disease. A** Violin plot showing the single-sample gene set enrichment analysis (ssGSEA) scores for the progression signature (refer to Supplementary Figure 3B and the methods section) across various cell clusters. These clusters were determined through a joint analysis of single-cell RNA sequencing (scRNA-seq) data from LNCaP-Residual samples, which include shSCR (CNTR1), TRIM24KD-1, and TRIM24KD-2 (as reported in Figure 1C). Different colors represent the cell clusters in the G1 phase, indicated by a yellow line, while Cluster 3 in the G2/S phase is marked by a green line. Statistical differences were calculated using the Wilcoxon test. **B** Gene set enrichment analysis of hallmark signatures across the identified cell clusters in the integrated scRNA-seq analysis of LNCaP-Residual tumors described in panel A. The dot plot shows the gene-enriched scores' false discovery rate (FDR) for each dataset among the cell clusters. **C-D** Panel **C** presents a UMAP plot, while panel **D** features the corresponding violin plot displaying the single-sample gene set expression scores for TRIM24 and AR in each LNCaP-Residual sample. Median pathway scores for each group are indicated, and fold-change values (the ratio of medians) are annotated between groups. Statistical significance was assessed using the Wilcoxon rank-sum test.

A

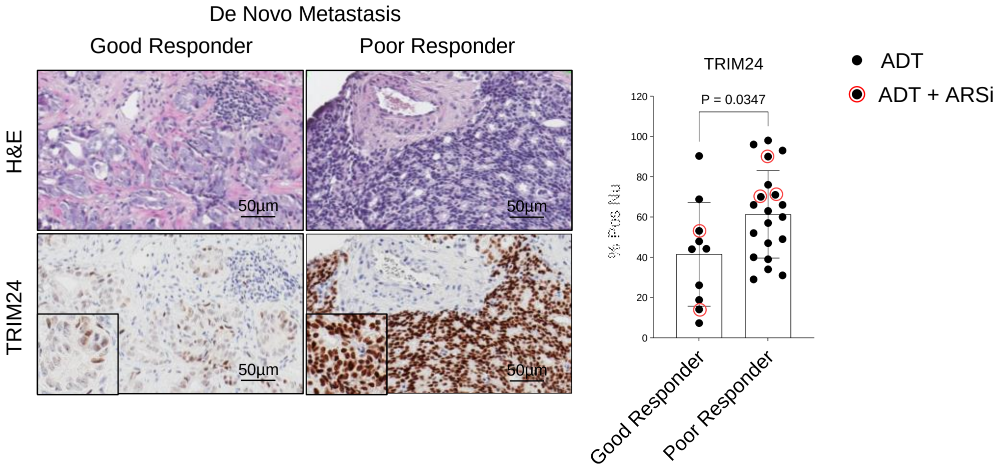


B

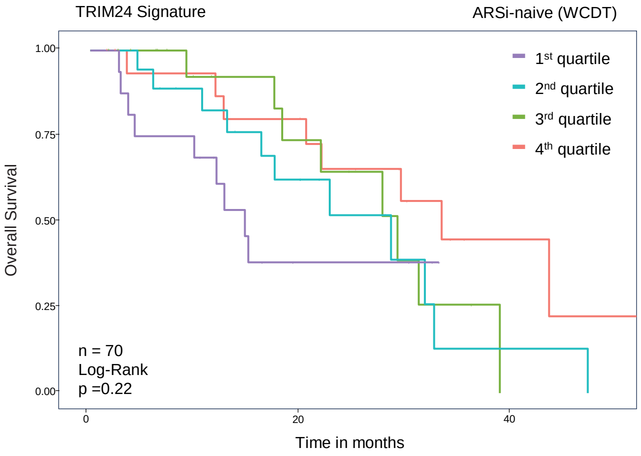


C

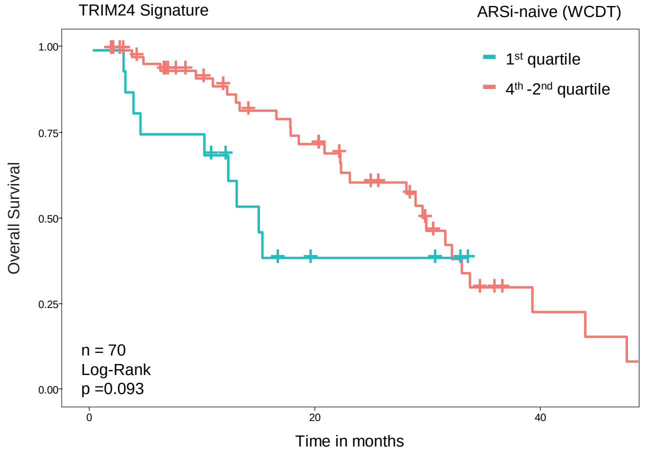


Figure S5

**Supplementary figure 5. TRIM24 is associated with poor response to ADT/ARSi.** A TRIM24 IHC analysis on human tumor biopsy of patients with de novo, high-volume and high grade, metastastic disease categorized into poor and good responders (see detail in Table SD1). On the left a representative image of tumor biopsy (scale bar 40 μm), and on the right a bar plot quantifies the number of tumor cells exhibiting positive nuclear staining for TRIM24. Each dot represents the quantification of an individual patient's biopsy. Tumor biopsies from patients treated in the first line with either androgen deprivation therapy (ADT) alone (black circle) or in combination with androgen receptor inhibitors (ARSi) (black circle with red around) are reported. **B-C** Kaplan-Meier curves illustrating the Overall Survival (OS) for 71 patients of enzalutamide/abiraterone-naïve CRPC patients (WCDT). The patients were divided into four quartiles based on the single-sample GSEA score computed on TRIM24 activity gene set, as shown in graph **B. C** Corresponding survival data comparing the first quartile with the binned, second to fourth quartiles (for more details, refer to the supplementary methods). Statistical differences were assessed using a log-rank test.

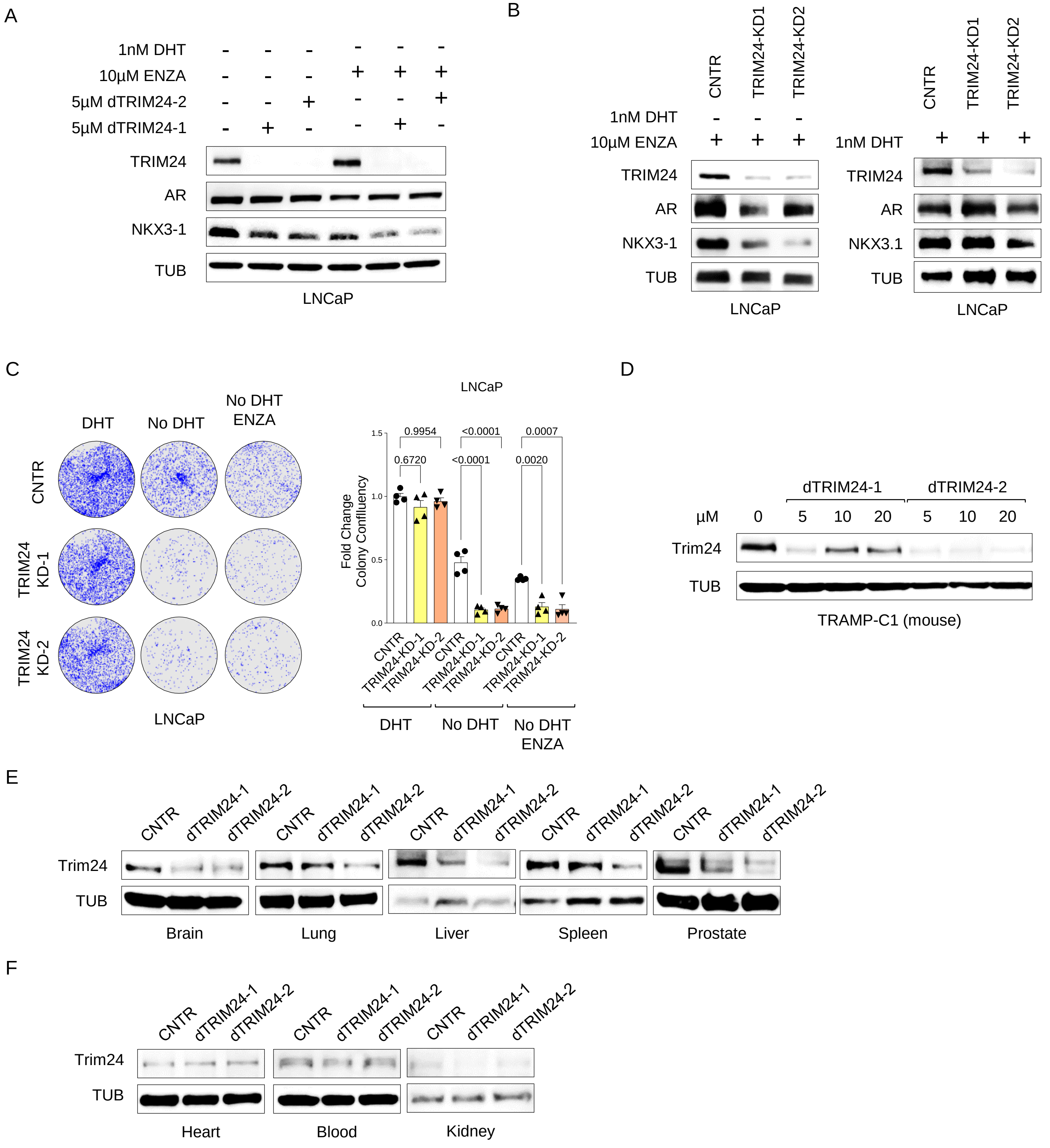
A
1nM DHT
10µM ENZA
5µM dTRIM24-2
5µM dTRIM24-1
TRIM24
AR
NKX3-1
TUB
LNCaP
B
CNTR
TRIM24-KD1
TRIM24-KD2
1nM DHT
10µM ENZA
TRIM24
AR
NKX3-1
TUB
LNCaP
NKX3.1
C
DHT
No DHT
No DHT ENZA
CNTR
TRIM24 KD-1
TRIM24 KD-2
LNCaP
Fold Change Colony Confluency
0.9954
<0.0001
0.0007
0.6720
<0.0001
0.0020
TRIM24-KD-1
TRIM24-KD-2
D
dTRIM24-1
dTRIM24-2
µM
0
5
10
20
Trim24
TUB
TRAMP-C1 (mouse)
E
dTRIM24-1
dTRIM24-2
Brain
Lung
Liver
Spleen
Prostate
F
Heart
Blood
Kidney

Figure S6

**Supplementary Figure 6. Effect of TRIM24 loss-of-function on tumor cell growth and AR signaling and effects of dTRIM24-1/2 on murine cells. A** Immunoblot analysis of indicated proteins in LNCaP cell cultures for 3 days in CSS medium alone (lane 1) or in combination with enzalutamide (lane 4) along with dTRIM24-1 (lane 2, 5) or dTRIM24-2 (lane 3, 6) at the indicated concentration. **B** Immunoblot analysis of indicated proteins in LNCaP knockdown cells (TRIM24KD-1 and TRIM24KD-2) compared to control cells (shSCR and CNTR). The cells were maintained for 3 days in CSS medium with 1 nM DHT (right) or CSS medium with 10 µM ENZA (left), along with dTRIM24-1 and dTRIM24-2 treatments at the indicated concentration. **C** Colonies formation assay LNCaP knockdown cells (TRIM24KD-1 and TRIM24KD-2) and control cells (shSCR and CNTR). The cells were maintained for 21 days in CSS with 1 nM DHT (DHT), CSS medium alone (no DHT), and CSS medium with 10 µM ENZA (no DHT + ENZA), along with dTRIM24-1 and dTRIM24-2 treatments. The concentrations used for the dTRIM24-1 and-2 were 5 µM. In the left panel, a representative image of colony confluency; in the right panel, a bar plot showing fold change in colony confluency for both cell lines at the indicated condition. The fold change is calculated based on the average colony confluency observed in the DHT condition. This data is derived from four independent experiments, with the bars representing the mean and the Standard Error of the Mean (SEM). Statistical significance was evaluated using a one-way ANOVA test. **D** Immunoblot analysis of the indicated proteins of murine TRAMPC1 cells treated with the indicated concentrations of dTRIM24-1 and dTRIM24-2 for 48h. **E-F** Immunoblot analysis of the indicated proteins of multiple murine tissues of NRG mice following daily intraperitoneal administration of 10mg/kg dTRIM24-1 and dTRIM24-2 for five days.

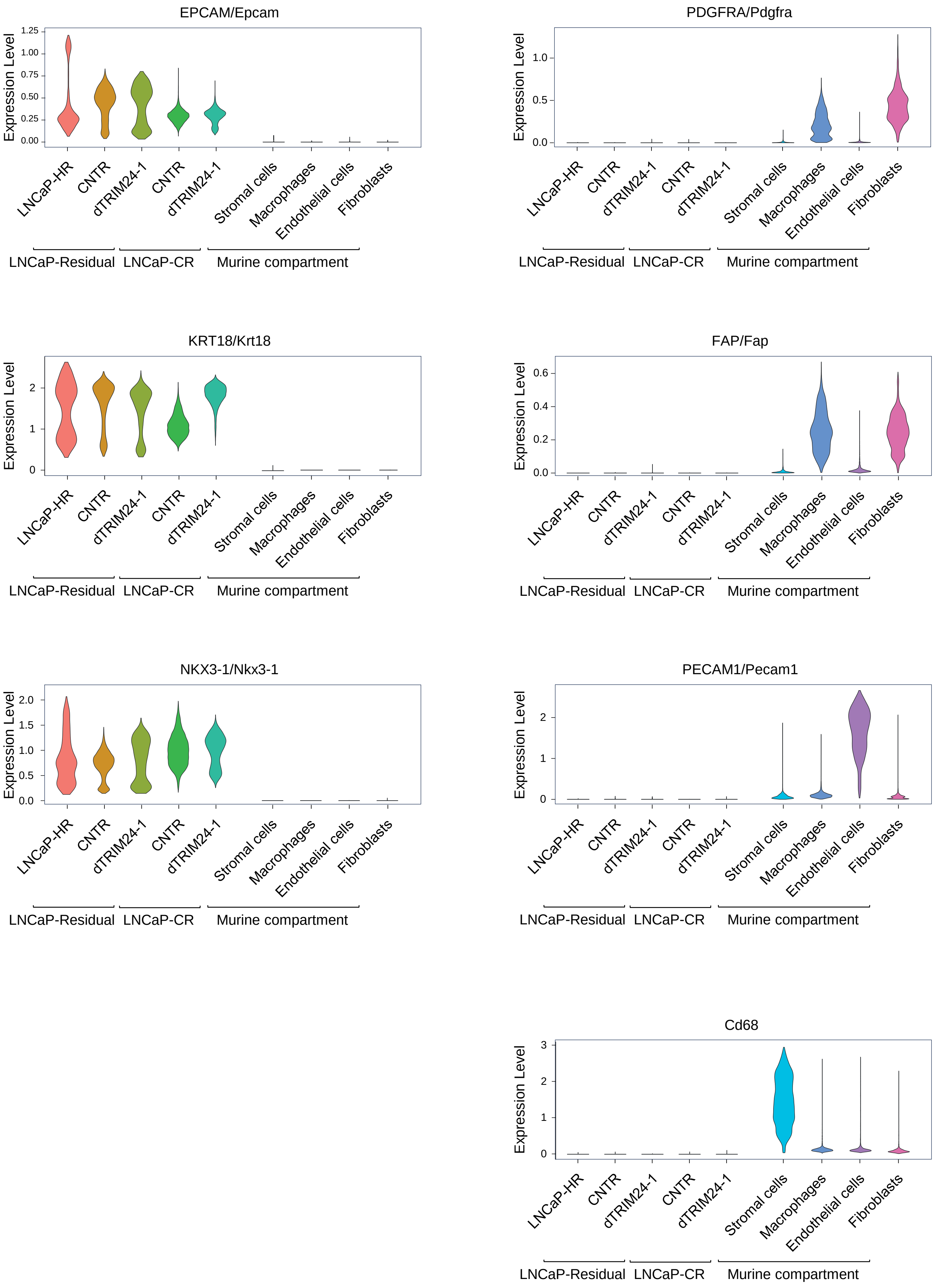

EPCAM/Epcam
Expression Level
1.25
1.00
0.75
0.50
0.25
0.00
PDGFRA/Pdgfra
1.0
0.5
0.0
KRT18/Krt18
2
1
0
FAP/Fap
0.6
0.4
0.2
0.0
NKX3-1/Nkx3-1
2.0
1.5
1.0
0.5
0.0
PECAM1/Pecam1
Cd68
3
LNCaP-HR
CNTR
dTRIM24-1
CNTR
dTRIM24-1
Stromal cells
Macrophages
Endothelial cells
Fibroblasts
LNCaP-Residual
LNCaP-CR
Murine compartment


Figure S7

**Supplementary Figure 7. Analysis of epithelial and stromal markers of human and murine compartment of scRNAseq data culled from LNCaP xenografts treated with ENZA and dTRIM24-1/2.** Violin plots illustrating the expression of canonical human prostate tumor markers (e.g., *EPCAM*, *NKX3-1*, *KRT18*) and murine stromal/immune markers (e.g., *FAP*, *PDGFRA*, *PECAM1*, *CD68*). The analysis includes LNCaP-HS (Hormone Sensitive, pre-castration), LNCaP-Residual (3 weeks post-castration), and LNCaP-CR (Castration-Resistant) under both vehicle (CNTR) and dTRIM24-1 treatments.

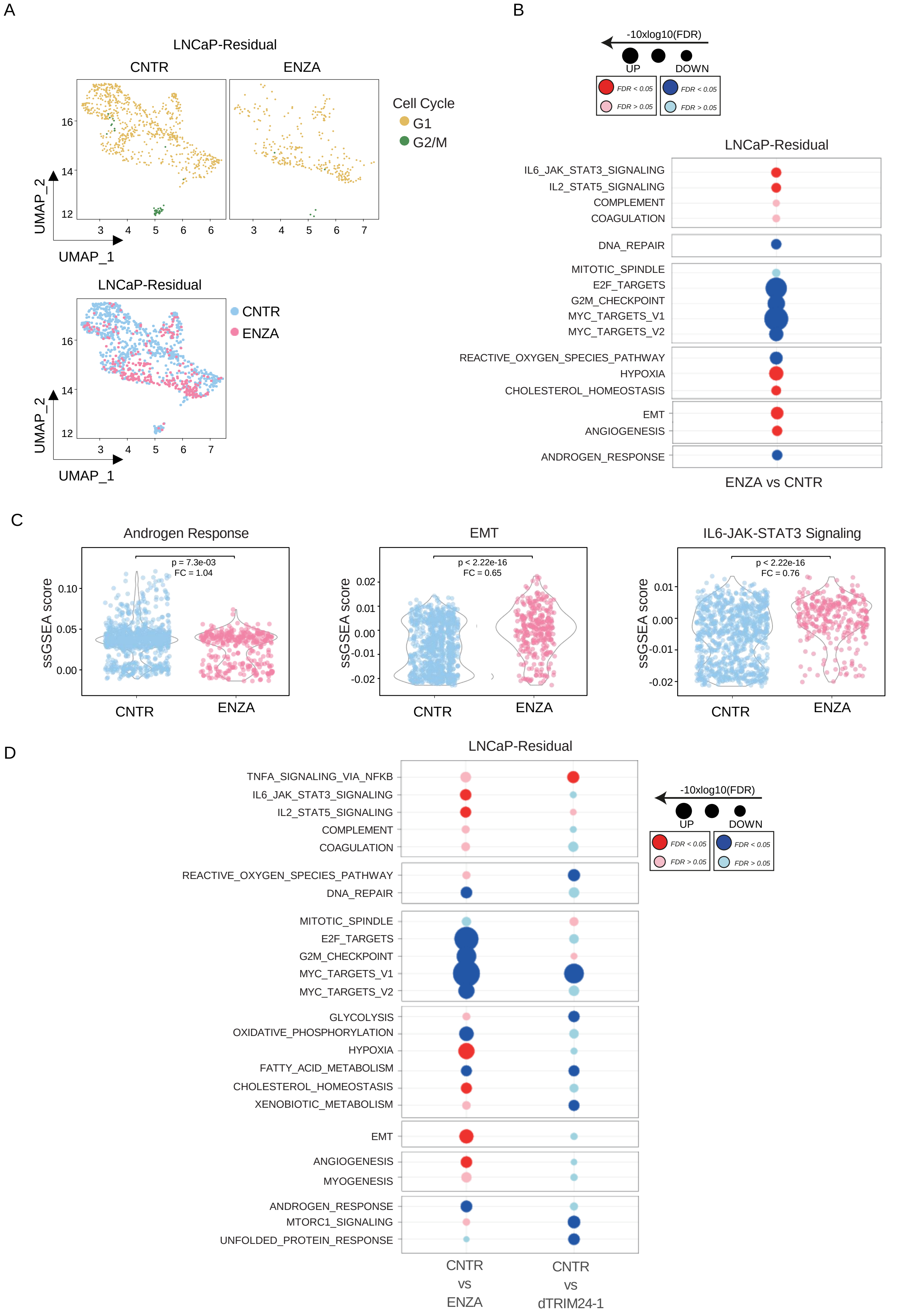

A
LNCaP-Residual
CNTR
ENZA
Cell Cycle
G1
G2/M
UMAP_2
UMAP_1
LNCaP-Residual
CNTR
ENZA
B
-10xlog10(FDR)
UP
DOWN
FDR < 0.05
FDR > 0.05
LNCaP-Residual
IL6_JAK_STAT3_SIGNALING
IL2_STAT5_SIGNALING
COMPLEMENT
COAGULATION
DNA_REPAIR
MITOTIC_SPINDLE
E2F_TARGETS
G2M_CHECKPOINT
MYC_TARGETS_V1
MYC_TARGETS_V2
REACTIVE_OXYGEN_SPECIES_PATHWAY
HYPOXIA
CHOLESTEROL_HOMEOSTASIS
EMT
ANGIOGENESIS
ANDROGEN_RESPONSE
ENZA vs CNTR
C
Androgen Response
p = 7.3e-03
FC = 1.04
ssGSEA score
EMT
p < 2.22e-16
FC = 0.65
IL6-JAK-STAT3 Signaling
p < 2.22e-16
FC = 0.76
CNTR
ENZA
D
LNCaP-Residual
TNFA_SIGNALING_VIA_NFKB
IL6_JAK_STAT3_SIGNALING
IL2_STAT5_SIGNALING
COMPLEMENT
COAGULATION
REACTIVE_OXYGEN_SPECIES_PATHWAY
DNA_REPAIR
MITOTIC_SPINDLE
E2F_TARGETS
G2M_CHECKPOINT
MYC_TARGETS_V1
MYC_TARGETS_V2
GLYCOLYSIS
OXIDATIVE_PHOSPHORYLATION
HYPOXIA
FATTY_ACID_METABOLISM
CHOLESTEROL_HOMEOSTASIS
XENOBIOTIC_METABOLISM
EMT
ANGIOGENESIS
MYOGENESIS
ANDROGEN_RESPONSE
MTORC1_SIGNALING
UNFOLDED_PROTEIN_RESPONSE
CNTR vs ENZA
CNTR vs dTRIM24-1


Figure S8

**Supplementary Figure 8. Enzalutamide promotes STAT3-signaling and EMT in LNCaP-residual tumors. A** The upper panel displays a UMAP plot of single-cell RNA sequencing (scRNA-seq) analysis data from LNCaP residual tumors treated after castration with either vehicle (CNTR) or enzalutamide (ENZA). The yellow highlights the G1 phase, while the green indicates cycling cells in the G2/M phases. The lower panel shows the cell cluster distribution in the vehicle-treated tumors (light blue dots; CNTR) compared to the enzalutamide-treated tumors (pink dots; ENZA). **B** Corresponding gene set enrichment analysis on pseudo bulk RNAseq of hallmark signatures. The dot plot shows the false discovery rate (FDR) of the indicated gene set enrichment scores comparing LNCaP-residual CNTR versus ENZA. **C** Violin plot displays the single-sample gene set enrichment scores for the indicated signatures in the scRNA-seq data of CNTR (light blue) and ENZA (pink) LNCaP-Residual sample. **D** Dot plot showing the false discovery rate (FDR) of the gene set enrichment analysis of hallmark signatures using pseudo bulk RNA sequencing, comparing the LNCaP-Residual tumors treated with vehicle to those treated with ENZA and those treated with dTRIM24-1

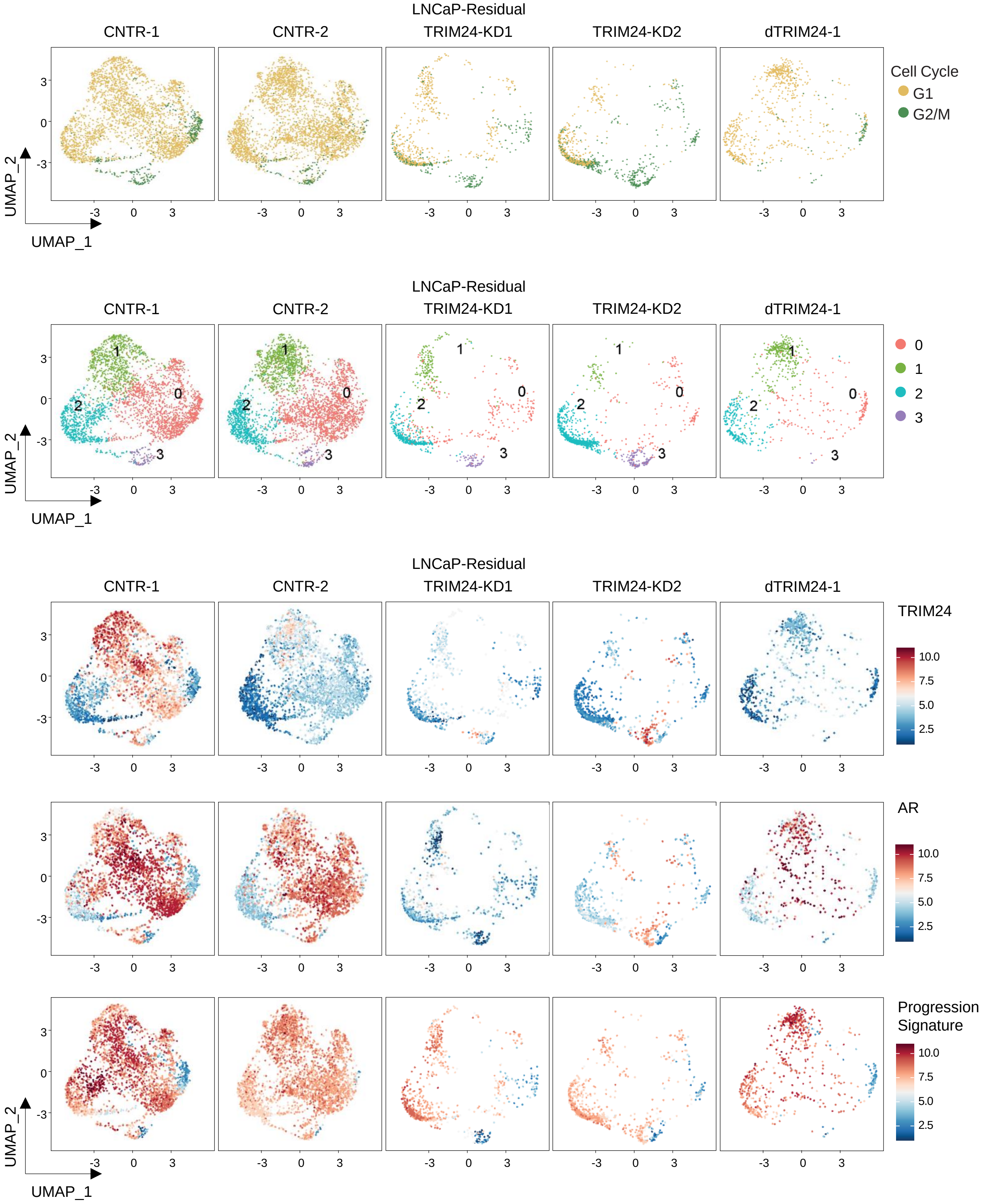
LNCaP-Residual
CNTR-1
CNTR-2
TRIM24-KD1
TRIM24-KD2
dTRIM24-1
Cell Cycle
G1
G2/M
UMAP_2
UMAP_1
0
1
2
3
TRIM24
AR
Progression Signature
10.0
7.5
5.0
2.5

Figure S9

**Supplementary Figure 9. Integrated scRNA-Seq analysis of LNCaP-Residual xenograft tumors.** This analysis compares the effects of TRIM24 knockdown with treatment using dTRIM24-1. The UMAP plot incorporates scRNA-Seq datasets from LNCaP-Residual samples, including shSCR (CNTR1), TRIM24KD-1, TRIM24KD-2, and samples treated after castration with dTRIM24-1, alongside a vehicle control (CNTR2). In the first panel, the different phases of the cell cycle are indicated, with G1 phase cells shown in yellow and G2/M phase cells in green. The second panel illustrates the various cell clusters identified in the analysis. The third and fourth panels present the gene expression levels for TRIM24 and AR, respectively. The fifth panel displays the gene set enrichment score for progression signatures (referred to Supplementary Figure 3B and the methods section).

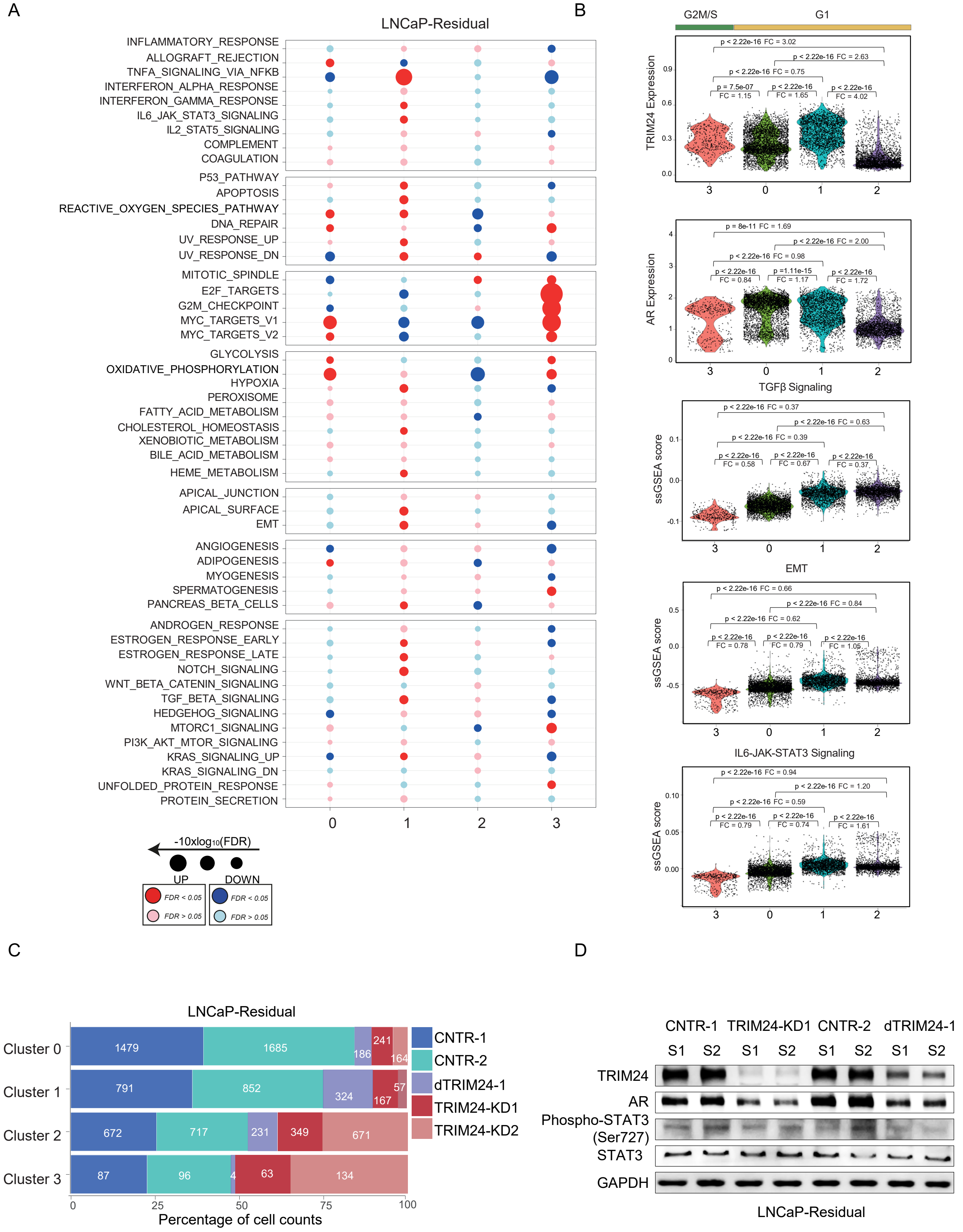
A
LNCaP-Residual
INFLAMMATORY_RESPONSE
ALLOGRAFT_REJECTION
TNFA_SIGNALING_VIA_NFKB
INTERFERON_ALPHA_RESPONSE
INTERFERON_GAMMA_RESPONSE
IL6_JAK_STAT3_SIGNALING
IL2_STAT5_SIGNALING
COMPLEMENT
COAGULATION
P53_PATHWAY
APOPTOSIS
REACTIVE_OXYGEN_SPECIES_PATHWAY
DNA_REPAIR
UV_RESPONSE_UP
UV_RESPONSE_DN
MITOTIC_SPINDLE
E2F_TARGETS
G2M_CHECKPOINT
MYC_TARGETS_V1
MYC_TARGETS_V2
GLYCOLYSIS
OXIDATIVE_PHOSPHORYLATION
HYPOXIA
PEROXISOME
FATTY_ACID_METABOLISM
CHOLESTEROL_HOMEOSTASIS
XENOBIOTIC_METABOLISM
BILE_ACID_METABOLISM
HEME_METABOLISM
APICAL_JUNCTION
APICAL_SURFACE
EMT
ANGIOGENESIS
ADIPOGENESIS
MYOGENESIS
SPERMATOGENESIS
PANCREAS_BETA_CELLS
ANDROGEN_RESPONSE
ESTROGEN_RESPONSE_EARLY
ESTROGEN_RESPONSE_LATE
NOTCH_SIGNALING
WNT_BETA_CATENIN_SIGNALING
TGF_BETA_SIGNALING
HEDGEHOG_SIGNALING
MTORC1_SIGNALING
PI3K_AKT_MTOR_SIGNALING
KRAS_SIGNALING_UP
KRAS_SIGNALING_DN
UNFOLDED_PROTEIN_RESPONSE
PROTEIN_SECRETION
0 1 2 3
-10xlog10(FDR)
UP DOWN
FDR < 0.05
FDR > 0.05
FDR < 0.05
FDR > 0.05
B
G2M/S G1
TRIM24 Expression
p < 2.22e-16 FC = 3.02
p < 2.22e-16 FC = 2.63
p < 2.22e-16 FC = 0.75
p = 7.5e-07 FC = 1.15
p < 2.22e-16 FC = 1.65
p < 2.22e-16 FC = 4.02
3 0 1 2
AR Expression
p = 8e-11 FC = 1.69
p < 2.22e-16 FC = 2.00
p < 2.22e-16 FC = 0.98
p < 2.22e-16 FC = 0.84
p =1.11e-15 FC = 1.17
p < 2.22e-16 FC = 1.72
3 0 1 2
TGFβ Signaling
ssGSEA score
p < 2.22e-16 FC = 0.37
p < 2.22e-16 FC = 0.63
p < 2.22e-16 FC = 0.39
p < 2.22e-16 FC = 0.58
p < 2.22e-16 FC = 0.67
p < 2.22e-16 FC = 0.37
3 0 1 2
EMT
ssGSEA score
p < 2.22e-16 FC = 0.66
p < 2.22e-16 FC = 0.84
p < 2.22e-16 FC = 0.62
p < 2.22e-16 FC = 0.78
p < 2.22e-16 FC = 0.79
p < 2.22e-16 FC = 1.05
3 0 1 2
IL6-JAK-STAT3 Signaling
ssGSEA score
p < 2.22e-16 FC = 0.94
p < 2.22e-16 FC = 1.20
p < 2.22e-16 FC = 0.59
p < 2.22e-16 FC = 0.79
p < 2.22e-16 FC = 0.74
p < 2.22e-16 FC = 1.61
3 0 1 2
C
LNCaP-Residual
Cluster 0
Cluster 1
Cluster 2
Cluster 3
1479 1685 186 241 164
791 852 324 167 57
672 717 231 349 671
87 96 4 63 134
0 25 50 75 100
Percentage of cell counts
CNTR-1
CNTR-2
dTRIM24-1
TRIM24-KD1
TRIM24-KD2
D
CNTR-1 TRIM24-KD1 CNTR-2 dTRIM24-1
S1 S2 S1 S2 S1 S2 S1 S2
TRIM24
AR
Phospho-STAT3 (Ser727)
STAT3
GAPDH
LNCaP-Residual

Figure S10

**Supplementary figure 10. Pathway analysis of cell clusters associated with the integrated single-cell RNA sequencing (scRNA-seq) data from Supplementary Figure 9.** **A** Gene Set Enrichment Analysis of the identified cell clusters from the integrated scRNA-seq analysis is presented. The dot plot shows the false discovery rate (FDR) of each cell cluster's gene set enrichment score. **B** Corresponding violin plot displaying the gene expression scores for TRIM24 and AR and the gene set enrichment scores for the specified signatures across different cell clusters in tumor samples (refer to the UMAP plot in supplementary Figure 9). The cell cycle phases of each cluster, G1, and G2/M, are indicated at the top of the plot with yellow and green lines, respectively. **C** The stacked bar plots illustrate the percentage of cells (on the x-axis) per cluster (clusters 0–3; on the y-axis) across different samples. Each bar is color-coded to represent the five LNCaP-Residual tumors within each cluster. The experimental groups include controls (shSCR, CNTR), TRIM24-knockdown, and TRIM24-degradation conditions in the LNCaP-Residual tumor. The total cell count for each cluster across the sample is shown within the bars. The height of each bar represents the percentage of cell count for that cluster, enabling a comparison of overall cellular abundance and the specific proportions of each sample across the different conditions. **D** Immunoblot analysis on the indicated proteins in LNCAP-Residual tumors sampled three weeks after castration. The CNTR-1 (lanes 1,2) and TRIM24-KD1 (lanes 3,4) tumors were compared with those treated with vehicle (CNTR-2; lanes 5,6) or dTRIM24-1 (lanes 7,8). Duplicates from 4 pooled tumors for each sample were analyzed.

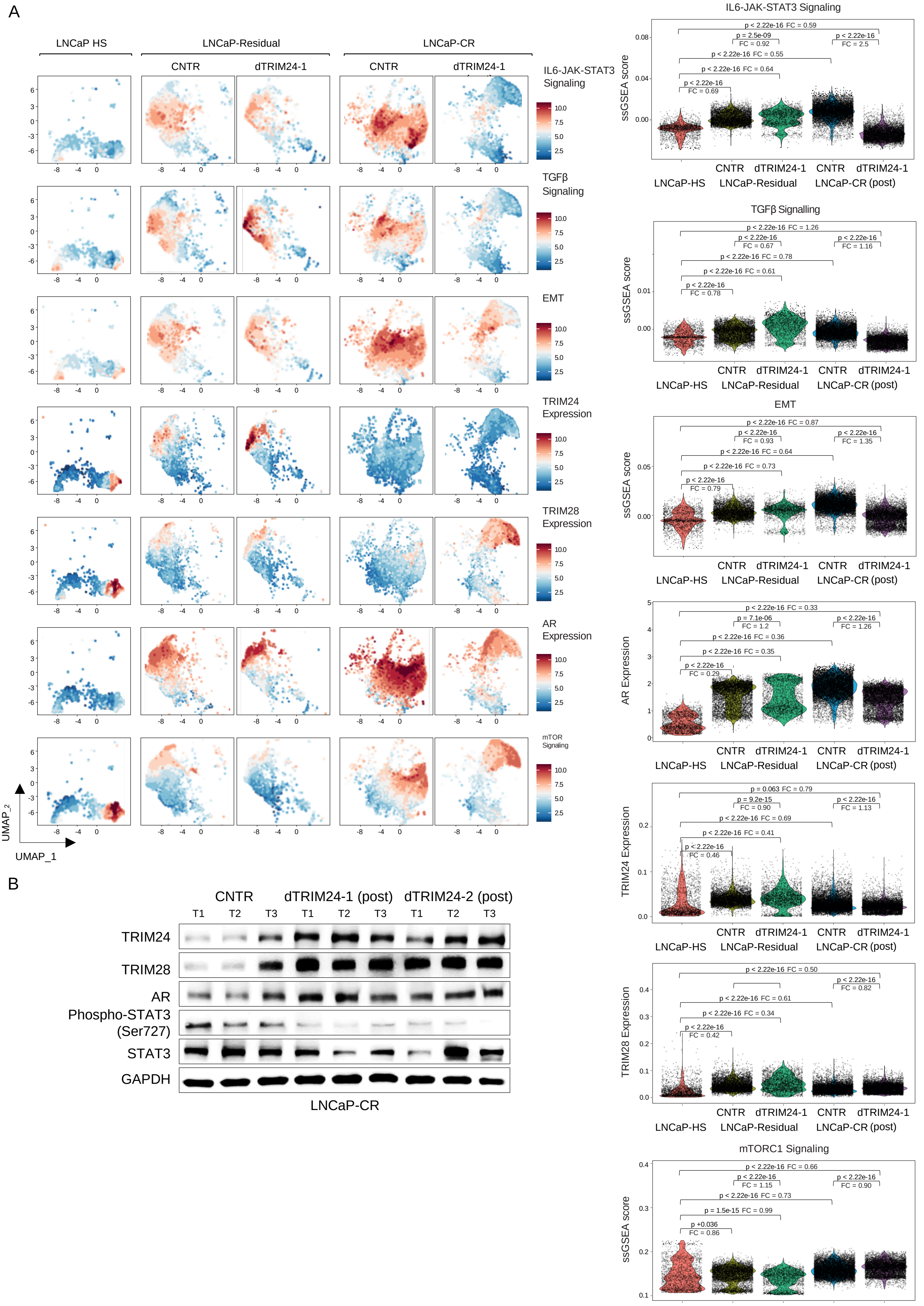

A
LNCaP HS
LNCaP-Residual
LNCaP-CR
CNTR
dTRIM24-1
IL6-JAK-STAT3 Signaling
TGFβ Signaling
EMT
TRIM24 Expression
TRIM28 Expression
AR Expression
mTOR Signaling
UMAP_2
UMAP_1
IL6-JAK-STAT3 Signaling
TGFβ Signalling
EMT
mTORC1 Signaling
ssGSEA score
AR Expression
TRIM24 Expression
TRIM28 Expression
LNCaP-HS
LNCaP-Residual
LNCaP-CR (post)
B
CNTR
dTRIM24-1 (post)
dTRIM24-2 (post)
T1
T2
T3
TRIM24
TRIM28
AR
Phospho-STAT3 (Ser727)
STAT3
GAPDH
LNCaP-CR


Figure S11

**Supplementary Figure 11. scRNA-Seq analysis of longitudinal LNCaP xenograft model**. Panels **A** and **B** show a UMAP plot (A) and the corresponding violin plot (B) that compare the gene set enrichment scores for the indicated signaling pathways and the expression levels of specified genes. The analysis includes hormone-sensitive LNCAP(-HS) tumors before castration, LNCAP-residual tumors after castration treated with either vehicle (CNTR) or dTRIM24-1, and LNCAP-CR tumors that relapsed after castration, also collected post-treatment with either vehicle (CNTR) or dTRIM24-1. **C** Immunoblot analysis of the indicated proteins in LNCAP-CR samples collected post-castration. The samples were treated with vehicle (CNTR; lanes 1-3), dTRIM24-1 (lanes 4-6), or dTRIM24-2 (lanes 7-9). Biological triplicates for each sample were analyzed.

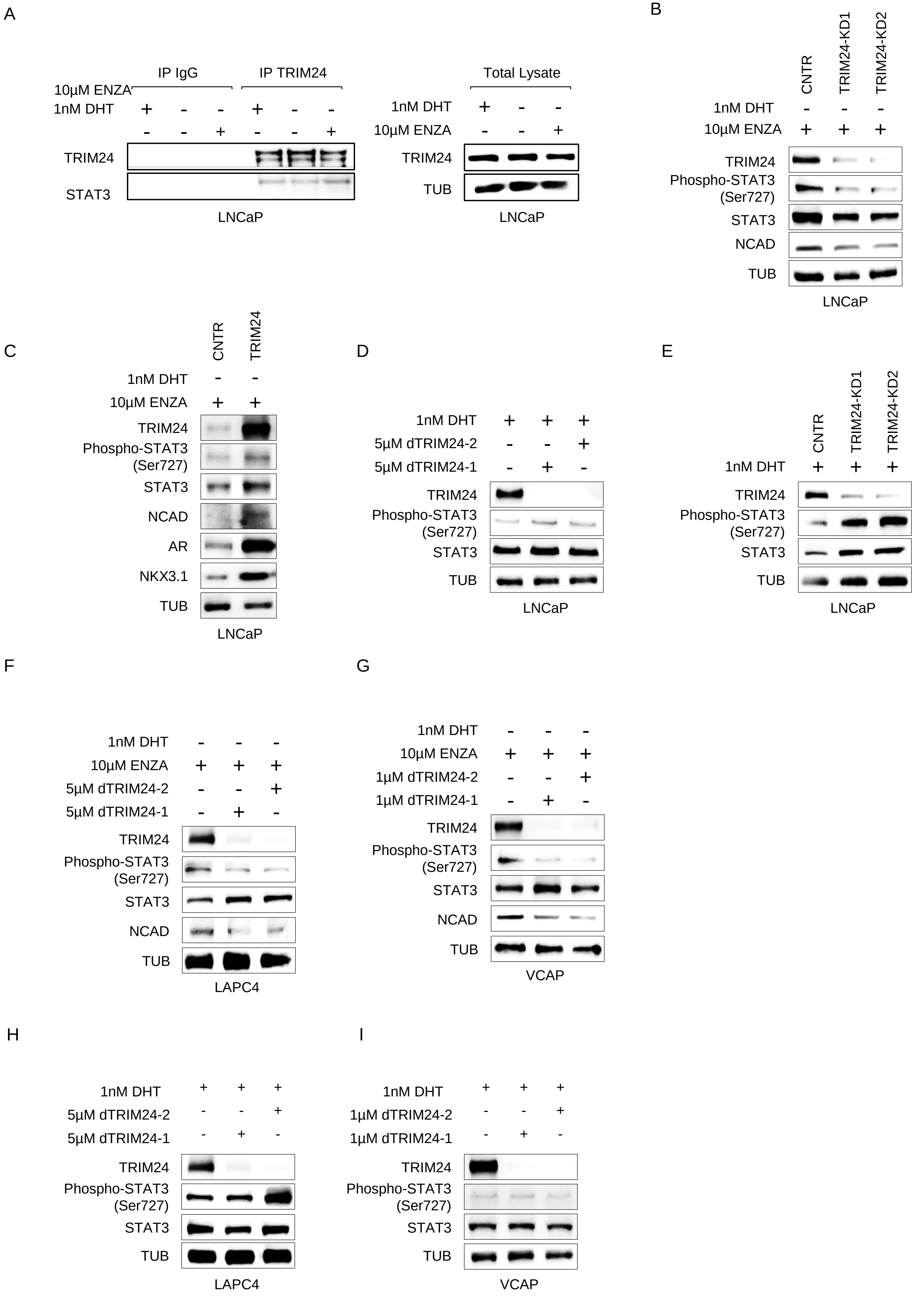

A
IP IgG
IP TRIM24
10µM ENZA
1nM DHT
TRIM24
STAT3
LNCaP
Total Lysate
1nM DHT
10µM ENZA
TRIM24
TUB
LNCaP
B
CNTR
TRIM24-KD1
TRIM24-KD2
1nM DHT
10µM ENZA
TRIM24
Phospho-STAT3 (Ser727)
STAT3
NCAD
TUB
LNCaP
C
CNTR
TRIM24
1nM DHT
10µM ENZA
TRIM24
Phospho-STAT3 (Ser727)
STAT3
NCAD
AR
NKX3.1
TUB
LNCaP
D
1nM DHT
5µM dTRIM24-2
5µM dTRIM24-1
TRIM24
Phospho-STAT3 (Ser727)
STAT3
TUB
LNCaP
E
CNTR
TRIM24-KD1
TRIM24-KD2
1nM DHT
TRIM24
Phospho-STAT3 (Ser727)
STAT3
TUB
LNCaP
F
1nM DHT
10µM ENZA
5µM dTRIM24-2
5µM dTRIM24-1
TRIM24
Phospho-STAT3 (Ser727)
STAT3
NCAD
TUB
LAPC4
G
1nM DHT
10µM ENZA
1µM dTRIM24-2
1µM dTRIM24-1
TRIM24
Phospho-STAT3 (Ser727)
STAT3
NCAD
TUB
VCAP
H
1nM DHT
5µM dTRIM24-2
5µM dTRIM24-1
TRIM24
Phospho-STAT3 (Ser727)
STAT3
TUB
LAPC4
I
1nM DHT
1µM dTRIM24-2
1µM dTRIM24-1
TRIM24
Phospho-STAT3 (Ser727)
STAT3
TUB
VCAP


Figure S12

**Supplementary figure 12. TRIM24 is required for STAT3 activity induction upon ADT and ENZA. A** Immunoprecipitation (IP) of TRIM24 and IgG control in LNCaP cells followed by immunoblotting for TRIM24 and STAT3. On the right, immunoblot analysis of input lysates for the indicated proteins. LNCaP cells maintained for 3 days in CSS medium alone (lane 2), in combination with DHT (CSS+DHT, lane 1), or with enzalutamide (CSS+ENZA, lane 3) at the indicated concentration. **B** Immunoblot analysis of indicated proteins in LNCaP cells with TRIM24 silencing (TRIM24-KD1 in lane 2, TRIM24-KD2 in lane 3) and in control cells (CNTR in lane 1) maintained for 3 days in CSS medium containing enzalutamide at the specified concentration. **C** Immunoblot analysis of indicated proteins in LNCaP cells overexpressing TRIM24 (ENZA, TRIM24 lane 2) and in control cells (ENZA, CNTR lane 1) cultured for 3 days in CSS medium containing enzalutamide at the specified concentration. **D** Immunoblot analysis of indicated proteins in LNCaP maintained for 3 days in CSS medium with 1nM DHT (DHT, lane 1) alone or in combination with dTRIM24-1 (DHT+dTRIM24-1, lane 2) or dTRIM24-2 (DHT+dTRIM24-1, lane 3) at the indicated concentration. **E** Immunoblot analysis of indicated proteins in LNCaP cells with TRIM24 silencing (TRIM24-KD1 in lane 2, TRIM24-KD2 in lane 3) and in control cells (CNTR in lane 1) maintained for 3 days in CSS medium with 1nM DHT a the indicate concentration. **F-G** Immunoblot analysis of indicated proteins in LAPC4 **(F)** and VCAP **(G)** cultures for 3 days in CSS medium with enzalutamide (ENZA, lane 1) alone or in combination with dTRIM24-1 (ENZA+dTRIM24-1, lane 2) or dTRIM24-2 (ENZA+dTRIM24-1, lane 3) at the indicated concentration. **H-I** Immunoblot analysis of indicated proteins in LAPC4 **(H)** and VCAP **(I)** cultures for 3 days in CSS medium with 1nM DHT (DHT, lane 1) alone or in combination with dTRIM24-1 (DHT+dTRIM24-1, lane 2) or dTRIM24-2 (DHT+dTRIM24-1, lane 3) at the indicated concentration.

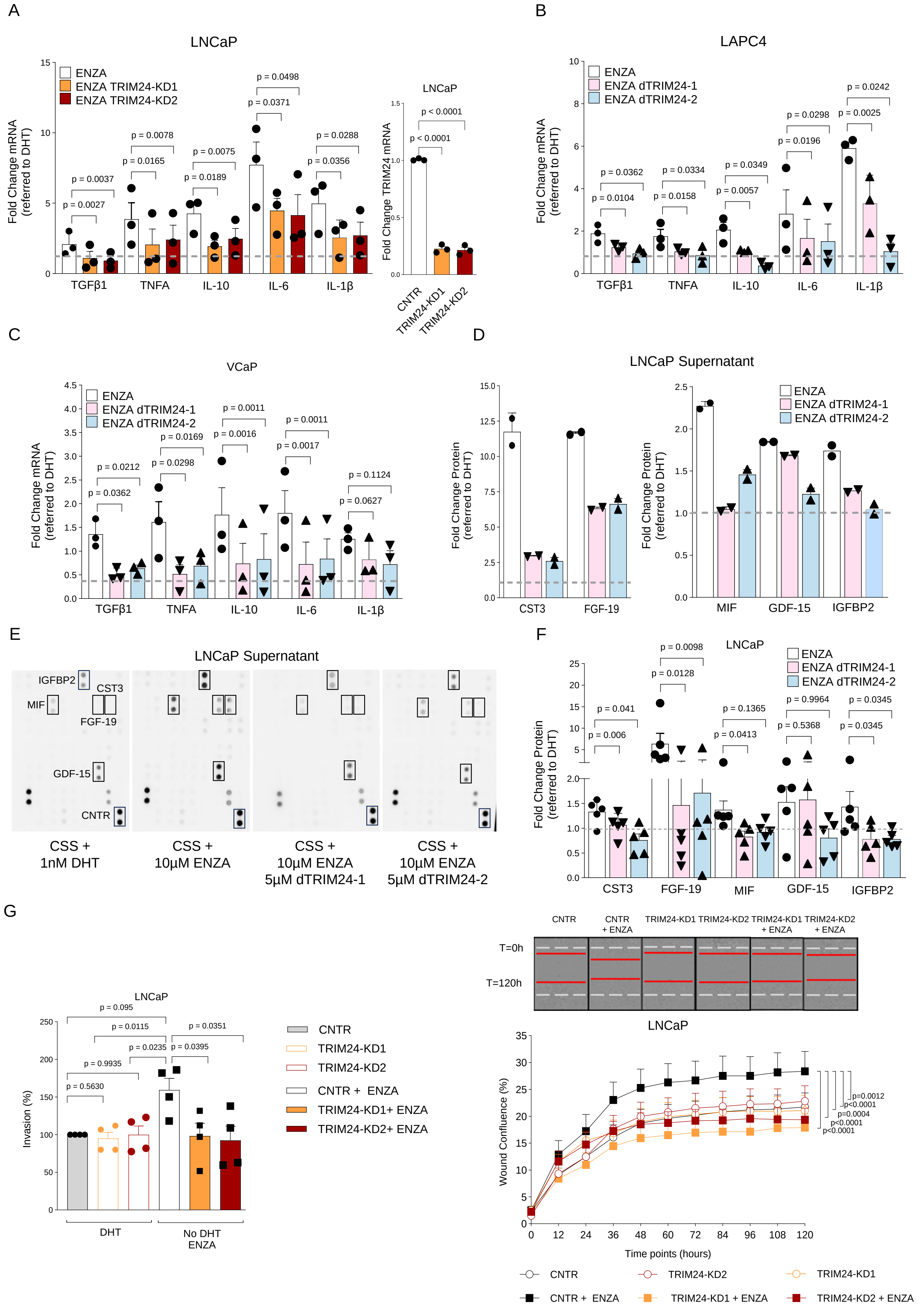
A
LNCaP
ENZA
ENZA TRIM24-KD1
ENZA TRIM24-KD2
Fold Change mRNA (referred to DHT)
TGFβ1
TNFA
IL-10
IL-6
IL-1β
Fold Change TRIM24 mRNA
p < 0.0001
CNTR
TRIM24-KD1
TRIM24-KD2
B
LAPC4
ENZA dTRIM24-1
ENZA dTRIM24-2
C
VCaP
D
LNCaP Supernatant
Fold Change Protein (referred to DHT)
CST3
FGF-19
MIF
GDF-15
IGFBP2
E
CNTR
CSS + 1nM DHT
CSS + 10µM ENZA
CSS + 10µM ENZA 5µM dTRIM24-1
CSS + 10µM ENZA 5µM dTRIM24-2
F
G
Invasion (%)
DHT
No DHT ENZA
CNTR + ENZA
TRIM24-KD1+ ENZA
TRIM24-KD2+ ENZA
T=0h
T=120h
Wound Confluence (%)
Time points (hours)

Figure S13

**Supplementary Figure 13. TRIM24 loss-of-function reduces cytokines induced by ADT and ENZA. A** Bar plot showing mRNA expression analysis of the indicated genes in LNCaP cells silenced for TRIM24 (TRIM24-KD1 and TRIM24-KD2) and control (CNTR) cultured for 3 days in CSS+DHT, CSS+ENZA (white bar), CSS+ENZA+dTRIM24-1 (dark yellow bar), or CSS+ENZA+dTRIM24-2 (dark red bar) by qPCR. **B-C** Bar plot showing mRNA expression analysis of the indicated genes in LAPC4 **(B)** and VCAP **(C)** cells, maintained under the same conditions described in Supplementary Figure 13A and Figure 4D. In **A-C**, the fold change is calculated based on the average expression of the DHT condition (dashed gray line). Data reported from at least 3 independent experiments, bars represent mean with Standard Error of Mean (SEM). Statistical significance is determined using an unpaired Student's t-test. **D** Cytokine array incubated with the supernatants of the LNCaP cells maintained for 3 days in CSS+DHT, CSS+ENZA, CSS+ENZA+dTRIM24-1, and CSS+ENZA+dTRIM24-2. The bar plot displays the top five robustly secreted factors that show an increase in the CSS+ENZA medium compared to CSS+DHT. All factors are down-modulated after the addition of 5 μM dTRIM24-1 or TRIM24-2. The Fold Changes (FC) of the signaling (density pixel) of each spot were calculated for the average signaling of the corresponding DHT condition (dashed gray line). The dots represent technical replicates from one cytokine array. **E** Image of the arrays with the spots for CST3, FGF-19, MIF, GDF-15, and IGFBP2 are indicated by squares. **F** Corresponding validation of mRNA expression changes of the indicated cytokines by qPCR. **G** Trans-well invasion assay (left) and Wound healing assay (right) on LNCaP cells silenced for TRIM24 (TRIM24-KD1 and TRIM24-KD2) and control (CNTR) cells cultured for 3 days in CSS+DHT (CNTR), CSS+ENZA, CSS+ENZA+dTRIM24-1, and CSS+ENZA+dTRIM24-2. In the left panel, LNCaP cells were incubated in a Boyden chamber for 24 hours. After staining with crystal violet, the migrated cells were quantified, showing significant differences, with bars representing the mean (± SEM) from four independent experiments and statistical significance analyzed using an unpaired Student's t-test. The right panel features the wound healing assay, where LNCaP cells were monitored over 120 hours post-scratch using the IncuCyte system. The graph displays the relative wound density, with bars indicating the mean (± SEM) from a representative experiment conducted in triplicate. Microscope images (10X magnification) show the wound at time point 0 (white line) and 120 hours (red line). Statistical significance was confirmed using an unpaired Student's t-test, reinforcing the findings.

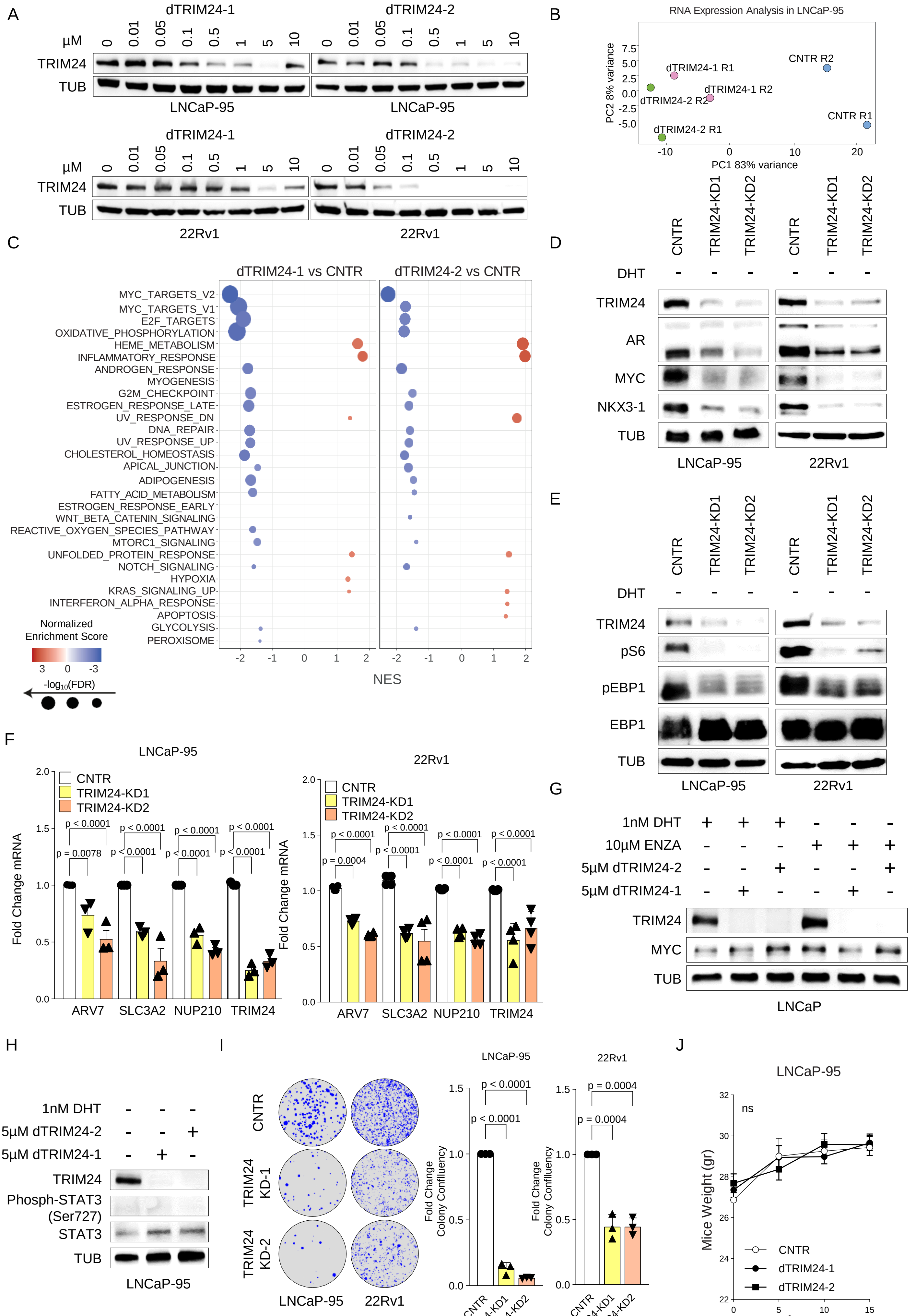

A
dTRIM24-1
dTRIM24-2
μM
0 0.01 0.05 0.1 0.5 1 5 10
TRIM24
TUB
LNCaP-95
22Rv1
B
RNA Expression Analysis in LNCaP-95
PC2 8% variance
PC1 83% variance
CNTR R1
CNTR R2
dTRIM24-1 R1
dTRIM24-1 R2
dTRIM24-2 R1
dTRIM24-2 R2
C
dTRIM24-1 vs CNTR
dTRIM24-2 vs CNTR
MYC_TARGETS_V2
MYC_TARGETS_V1
E2F_TARGETS
OXIDATIVE_PHOSPHORYLATION
HEME_METABOLISM
INFLAMMATORY_RESPONSE
ANDROGEN_RESPONSE
MYOGENESIS
G2M_CHECKPOINT
ESTROGEN_RESPONSE_LATE
UV_RESPONSE_DN
DNA_REPAIR
UV_RESPONSE_UP
CHOLESTEROL_HOMEOSTASIS
APICAL_JUNCTION
ADIPOGENESIS
FATTY_ACID_METABOLISM
ESTROGEN_RESPONSE_EARLY
WNT_BETA_CATENIN_SIGNALING
REACTIVE_OXYGEN_SPECIES_PATHWAY
MTORC1_SIGNALING
UNFOLDED_PROTEIN_RESPONSE
NOTCH_SIGNALING
HYPOXIA
KRAS_SIGNALING_UP
INTERFERON_ALPHA_RESPONSE
APOPTOSIS
GLYCOLYSIS
PEROXISOME
NES
Normalized Enrichment Score
-log10(FDR)
D
CNTR TRIM24-KD1 TRIM24-KD2 CNTR TRIM24-KD1 TRIM24-KD2
DHT
TRIM24
AR
MYC
NKX3-1
TUB
LNCaP-95
22Rv1
E
CNTR TRIM24-KD1 TRIM24-KD2 CNTR TRIM24-KD1 TRIM24-KD2
DHT
TRIM24
pS6
pEBP1
EBP1
TUB
LNCaP-95
22Rv1
F
LNCaP-95
22Rv1
CNTR
TRIM24-KD1
TRIM24-KD2
Fold Change mRNA
p < 0.0001
p = 0.0078
p = 0.0004
ARV7 SLC3A2 NUP210 TRIM24
G
1nM DHT
10μM ENZA
5μM dTRIM24-2
5μM dTRIM24-1
TRIM24
MYC
TUB
LNCaP
H
1nM DHT
5μM dTRIM24-2
5μM dTRIM24-1
TRIM24
Phosph-STAT3 (Ser727)
STAT3
TUB
LNCaP-95
I
CNTR
TRIM24 KD-1
TRIM24 KD-2
LNCaP-95
22Rv1
Fold Change Colony Confluency
p < 0.0001
p = 0.0004
CNTR TRIM24-KD1 TRIM24-KD2
J
LNCaP-95
ns
Mice Weight (gr)
Days of Treatment
CNTR
dTRIM24-1
dTRIM24-2


Figure S14

**Supplementary Figure 14**. **TRIM24 loss-of-function inhibits the growth of AR-V7-positive prostate cancer cells**. Immunoblot analysis of the indicated proteins of LNCaP-95 (upper panel), and 22Rv1 (lower panel) cells incubated with the indicated concentrations of dTRIM24-1 and dTRIM24-2. **B-C** In **B** principal component analysis (PCA) of bulk RNA-seq data of LNCaP-95 cells with DMSO (CNTR; blue dot) dTRIM24-1 (pink dot) and dTRIM24-2 (green dot) for 48 hours at 5μM. Each dot represents a biological duplicate. In **C** the corresponding normalized gene set enrichment scores (NES) of the indicated hallmark signatures after administration of TRIM24-1 and dTRIM24-2 as compared to DMSO. **D** Immunoblot analysis of indicated proteins in LNCaP-95 (left panel) and 22Rv1 (right panel) maintained for 3 days in CSS medium without DHT, comparing to TRIM24-KD1 (lane 2) or dTRIM24-KD2 (lane 3) cells alongside the control (CNTR) cells. **E** The bar plot showing mRNA expression analysis of AR-V7 target genes in LNCaP-95 (on the left) and 22RV1 (on the right) cells following the silencing of TRIM24 (TRIM24-KD1 and TRIM24-KD2) compared to the control (CNTR) in CSS medium. The fold change is calculated based on the average expression levels in the control condition. Data is reported from at least three independent experiments, with bars representing the mean and the Standard Error of the Mean (SEM). Statistical significance was assessed using an unpaired Student's t-test. **F** Immunoblot analysis of indicated proteins of LNCaP cells maintained for 3 days in CSS medium without DHT. The cells were treated with either DHT (CSS+DHT, left) or enzalutamide (CSS+ENZA, left), each used alone (lane 1) or in combination with dTRIM24-1 (CSS+ENZA+dTRIM24-1, lane 2) or dTRIM24-2 at the indicated. **G** Immunoblot analysis of indicated proteins of LNCaP-95 cultured for 3 days in CSS alone (lane 1) or in combination with dTRIM24-1 (lane 2) or dTRIM24-2 (lane 3) at the indicated concentration. **H** Colonies formation assay for LNCaP-95 (left), 22Rv1 (right) cells following the silencing of tTRIM24 (TRIM24-KD1 and TRIM24-KD2) compared to the control (CNTR) in CSS medium without DHT for 21 days. On the **left**, a representative image of colony confluency; on the **right**, a bar plot showing fold change in colony confluency for both cell lines at the indicated condition. The fold change is calculated based on the average colony confluency observed in the control condition. This data is derived from four independent experiments, with the bars representing the mean and the Standard Error of the Mean (SEM). Statistical significance was evaluated using a one-way ANOVA test. **J** Weight kinetics of NRG mice bearing LNCaP-95 xenograft during the treatment with vehicle (n=6 mice), dTRIM24-1 (n=6 mice, 10 mg/kg), dTRIM24-2 (n=8 mice, 10 mg/kg), and enzalutamide (n=6 mice) for two weeks

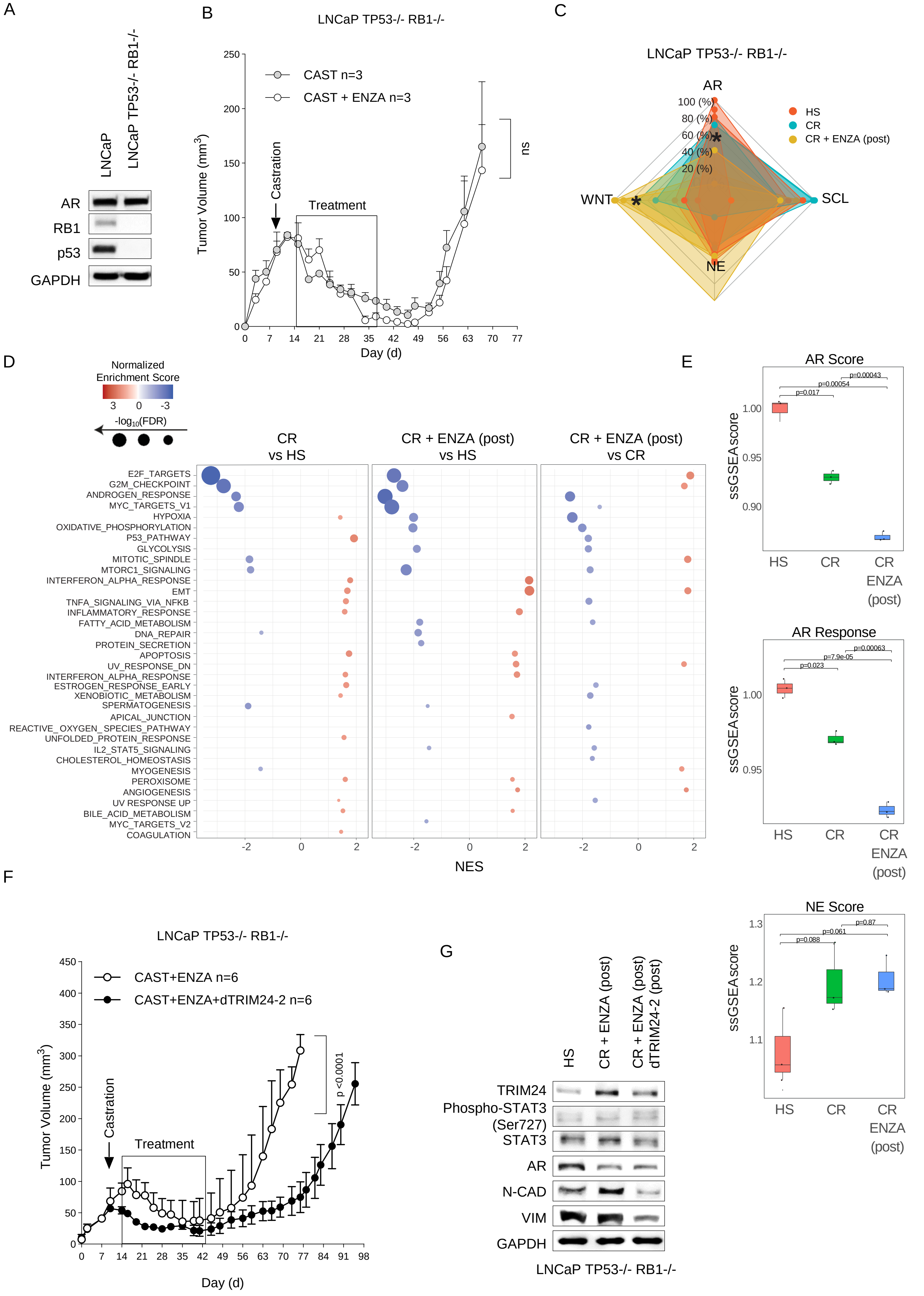
A
LNCaP
LNCaP TP53-/- RB1-/-
AR
RB1
p53
GAPDH
B
LNCaP TP53-/- RB1-/-
CAST n=3
CAST + ENZA n=3
Castration
Treatment
ns
Tumor Volume (mm3)
Day (d)
C
LNCaP TP53-/- RB1-/-
AR
WNT
SCL
NE
HS
CR
CR + ENZA (post)
D
Normalized Enrichment Score
-log10(FDR)
CR vs HS
CR + ENZA (post) vs HS
CR + ENZA (post) vs CR
E2F_TARGETS
G2M_CHECKPOINT
ANDROGEN_RESPONSE
MYC_TARGETS_V1
HYPOXIA
OXIDATIVE_PHOSPHORYLATION
P53_PATHWAY
GLYCOLYSIS
MITOTIC_SPINDLE
MTORC1_SIGNALING
INTERFERON_ALPHA_RESPONSE
EMT
TNFA_SIGNALING_VIA_NFKB
INFLAMMATORY_RESPONSE
FATTY_ACID_METABOLISM
DNA_REPAIR
PROTEIN_SECRETION
APOPTOSIS
UV_RESPONSE_DN
INTERFERON_ALPHA_RESPONSE
ESTROGEN_RESPONSE_EARLY
XENOBIOTIC_METABOLISM
SPERMATOGENESIS
APICAL_JUNCTION
REACTIVE_OXYGEN_SPECIES_PATHWAY
UNFOLDED_PROTEIN_RESPONSE
IL2_STAT5_SIGNALING
CHOLESTEROL_HOMEOSTASIS
MYOGENESIS
PEROXISOME
ANGIOGENESIS
UV RESPONSE UP
BILE_ACID_METABOLISM
MYC_TARGETS_V2
COAGULATION
NES
E
AR Score
AR Response
NE Score
ssGSEAscore
HS
CR
CR ENZA (post)
F
LNCaP TP53-/- RB1-/-
CAST+ENZA n=6
CAST+ENZA+dTRIM24-2 n=6
Castration
Treatment
p <0.0001
Tumor Volume (mm3)
Day (d)
G
HS
CR + ENZA (post)
CR + ENZA (post) dTRIM24-2 (post)
TRIM24
Phospho-STAT3 (Ser727)
STAT3
AR
N-CAD
VIM
GAPDH
LNCaP TP53-/- RB1-/-

Figure S15

**Supplementary Figure 15. Degradation of TRIM24 inhibits the growth of TP53/RB1-deficient LNCaP xenograft models. A** Immunoblot of the indicated proteins in parental LNCaP and LNCaPTP53-/-; RB1-/- cells. **B** Growth kinetics of LNCaPTP53-/-; RB1-/- xenografts in NRG mice. Castration was performed when the tumor size reached an average of 50-75 mm3, indicated by an arrow. Subsequently, mice were treated intraperitoneally with either the vehicle or Enzalutamide (ENZA) at 30 mg/kg for three weeks, five days a week. A black square visually highlights the treatment period. Statistical significance is assessed using a two-way ANOVA test. **C** Radar plot displaying the gene set enrichment scores of AR-positive (CRPC-AR), neuroendocrine (CRPC-NE), WNT-driven (CRPC-WNT), and stem cell-like subtype (CRPC-SCL) on bulk RNAseq of LNCaP$^{TP53-/-; RB1-/-}$ xenografts pre-castration (HS) and after relapse following castration (CR) and castration with daily intraperitoneal administration of 30mg/kg enzalutamide for three weeks (CR+ENZA). **D Normalized g**ene set enrichment scores (NES) of the indicated hallmark signatures on bulk RNA-seq data comparing LNCaPTP53-/-; RB1-/- -HS, -CR, -CR+ENZA xenografts. The significance (log10 FDR) is shown by dot size. **E** Corresponding box plot showing the single-sample enrichment (ssGSEA) for the AR score (left), AR response (middle), and NE score (right). The statistical significance of biological triplicates is calculated using the Wilcon-Cox test. **F** Growth kinetics of LNCaP$^{TP53-/-; RB1-/-}$ xenografts in NRG mice. Castration was performed when the tumor size reached an average of 50-75 mm3, indicated by an arrow. Upon castration, tumors were treated with either 30mg/kg enzalutamide alone or in combination with 10 mg/kg dTRIM24-2 for three weeks as described above. The black square marks the treatment period. Statistical significance is assessed using a two-way ANOVA test. **G** Immunoblot analysis of the indicated proteins in LNCaP$^{TP53-/-; RB1-/-}$ xenografts pre-castration and relapse after castration and post enzalutamide treatment alone or in combination with dTRIM24-2.

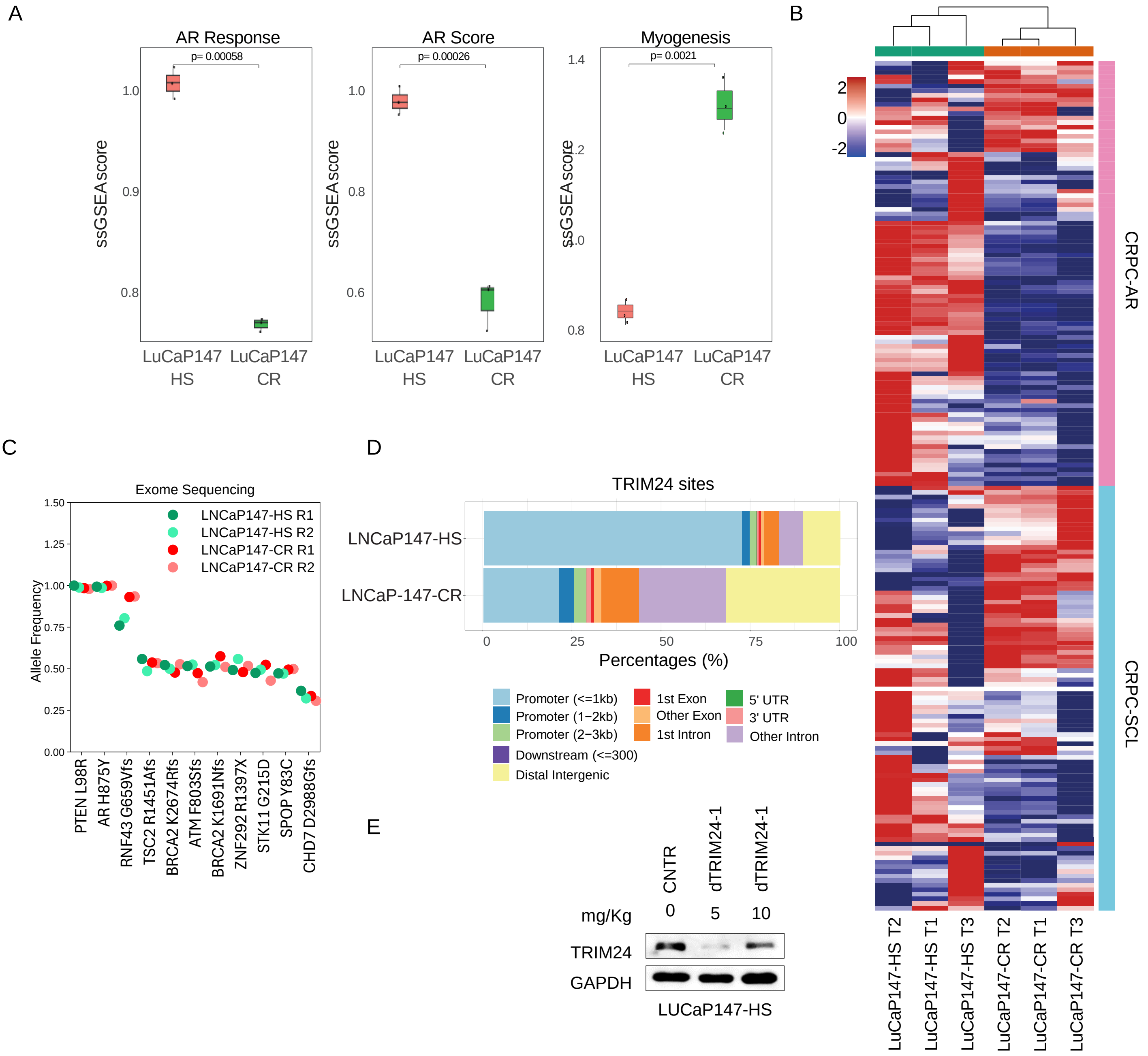
A
AR Response
p= 0.00058
AR Score
p= 0.00026
Myogenesis
p= 0.0021
ssGSEAscore
LuCaP147 HS
LuCaP147 CR
B
CRPC-AR
CRPC-SCL
LuCaP147-HS T2
LuCaP147-HS T1
LuCaP147-HS T3
LuCaP147-CR T2
LuCaP147-CR T1
LuCaP147-CR T3
C
Exome Sequencing
LNCaP147-HS R1
LNCaP147-HS R2
LNCaP147-CR R1
LNCaP147-CR R2
Allele Frequency
PTEN L98R
AR H875Y
RNF43 G659Vfs
TSC2 R1451Afs
BRCA2 K2674Rfs
ATM F803Sfs
BRCA2 K1691Nfs
ZNF292 R1397X
STK11 G215D
SPOP Y83C
CHD7 D2988Gfs
D
TRIM24 sites
LNCaP147-HS
LNCaP-147-CR
Percentages (%)
Promoter (<=1kb)
Promoter (1−2kb)
Promoter (2−3kb)
Downstream (<=300)
Distal Intergenic
1st Exon
Other Exon
1st Intron
5' UTR
3' UTR
Other Intron
E
CNTR
dTRIM24-1
dTRIM24-1
mg/Kg
0
5
10
TRIM24
GAPDH
LUCaP147-HS

Figure S16

**Supplementary Figure 16. Characterization of the LuCaP147 stem-cell-like lineage plasticity model. A** Box plot showing the single-sample enrichment (ssGSEA) for the AR score (left), AR response (middle), and Myogenesis (right) on bulk RNA-seq data comparing the hormone-sensitive LuCaP147(-HS) and castration-resistant LuCaP147(-CR) relapse after castration. The statistical significance of biological triplicates is calculated using the Wilcon-Cox test. **B** Heatmap showing normalized gene-set enrichment scores of bulk RNA-seq data of LuCaP147-HR and LuCaP147-CR xenografts using the gene signatures of the CRPC-AR and CRPC-SCL subtypes (see also Fig. 6B). **C** Plot showing the allele frequency for LuCaP147-HS and LuCaP147-CR. Each point represents the allele frequency for the indicated somatic point mutation from duplicate exome sequences. Each point was color-coded based on the samples: green for HS and red for CR. Biological replicates are marked with dark colors for replicate 1 (R1) and light colors for replicate 2 (R2)**. D** Genome-wide binding location of the TRIM24 peaks in LuCaP147-HS in comparison to LuCaP147-CR. The assessed binding distribution included regions up to 3 kb upstream of TSS (promoter), 3 kb downstream of TSS (downstream), 5′ and 3′ UTRs, coding exons, introns, and distal intergenic regions. **E** Immunoblot analysis of the indicated proteins showing TRIM24 reduction in LuCaP147-HS following daily intraperitoneal administration of either vehicle (CNTR) or dTRIM24-1 at doses of 5 mg/kg and 10 mg/kg over five days.

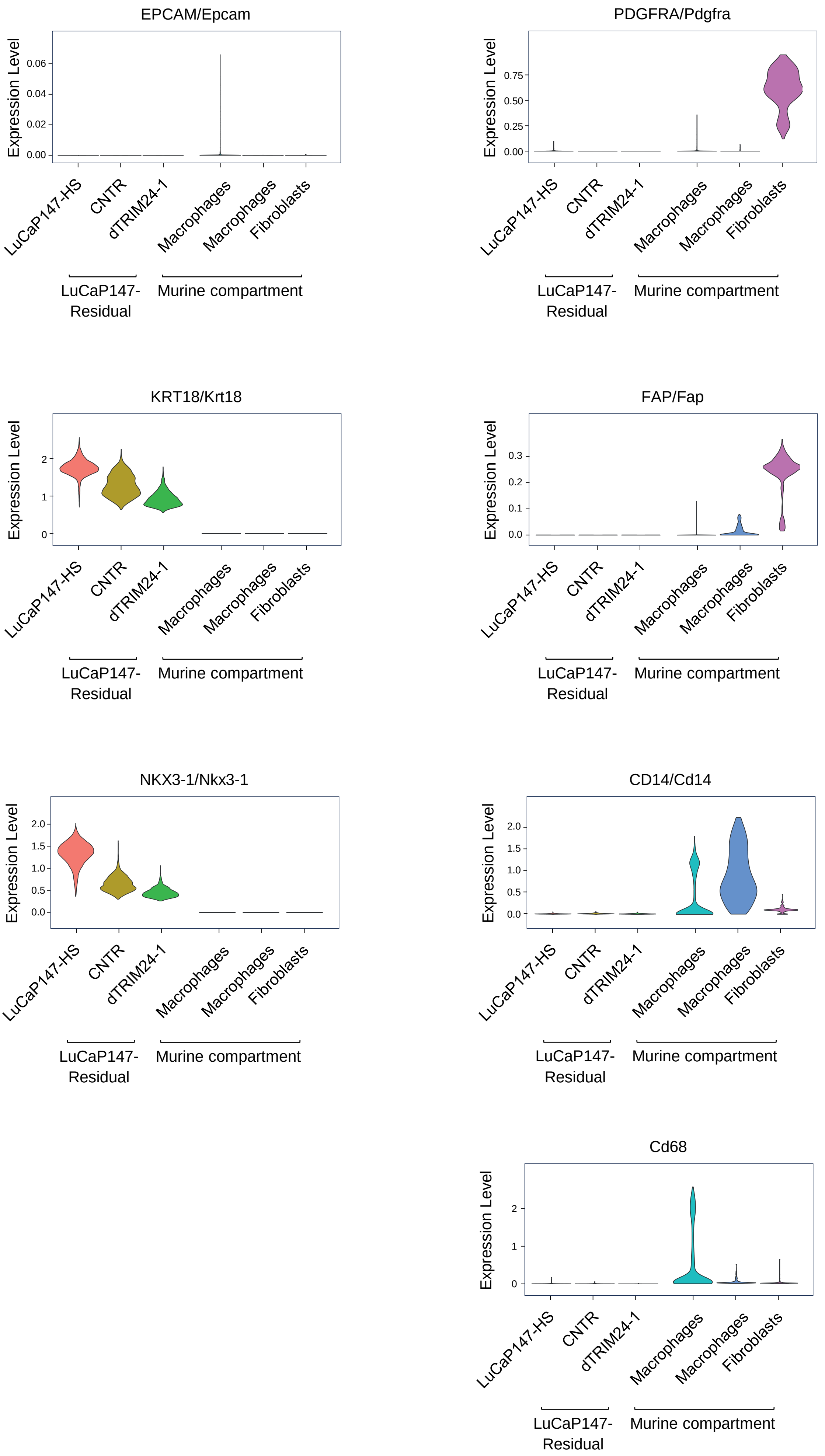
EPCAM/Epcam
Expression Level
0.00
0.02
0.04
0.06
LuCaP147-HS
CNTR
dTRIM24-1
Macrophages
Macrophages
Fibroblasts
LuCaP147-
Residual
Murine compartment
PDGFRA/Pdgfra
Expression Level
0.00
0.25
0.50
0.75
KRT18/Krt18
Expression Level
0
1
2
FAP/Fap
Expression Level
0.0
0.1
0.2
0.3
NKX3-1/Nkx3-1
Expression Level
0.0
0.5
1.0
1.5
2.0
CD14/Cd14
Expression Level
0.0
0.5
1.0
1.5
2.0
Cd68
Expression Level
0
1
2

Figure S17

**Supplementary figure 17. Analysis of epithelial and stromal markers of the human and murine compartments of scRNAseq data culled from LuCaP147 xenografts.** Violin plots illustrating the expression of canonical human prostate tumor markers (e.g., *EPCAM*, *NKX3-1*, *KRT18*) and murine stromal/immune markers (e.g., *FAP*, *PDGFRA*, CD14, *CD68*). The analysis included sample from LuCaP147-HS (Hormone Sensitive, pre-castration), and LuCaP147-Residual (3 weeks post-castration), which were treated with either vehicle (CNTR) or dTRIM24-1.

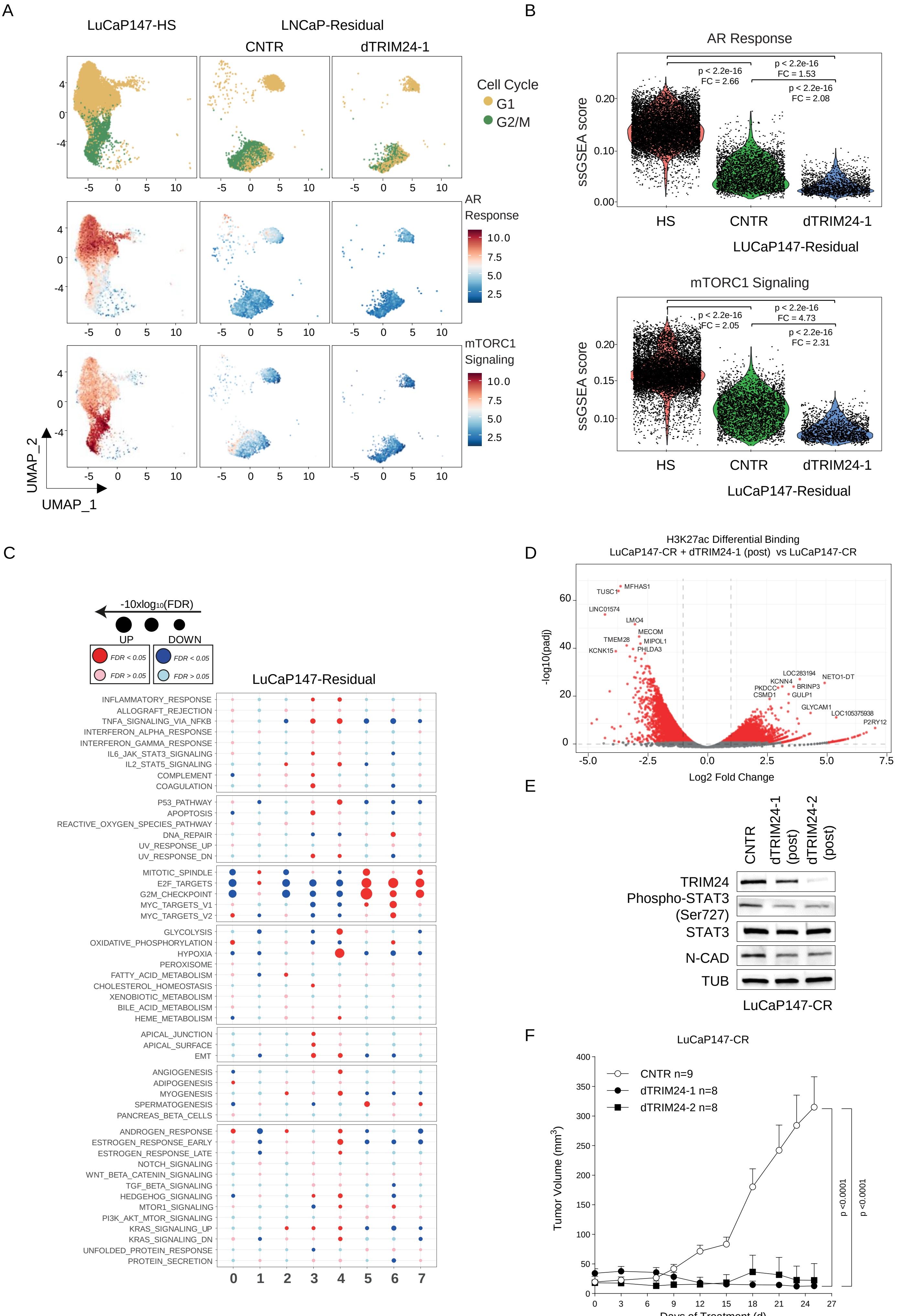
A
LuCaP147-HS
LNCaP-Residual
CNTR
dTRIM24-1
Cell Cycle
G1
G2/M
AR Response
mTORC1 Signaling
10.0
7.5
5.0
2.5
UMAP_2
UMAP_1
B
AR Response
ssGSEA score
p < 2.2e-16
FC = 2.66
p < 2.2e-16
FC = 1.53
p < 2.2e-16
FC = 2.08
HS
CNTR
dTRIM24-1
LUCaP147-Residual
mTORC1 Signaling
p < 2.2e-16
FC = 2.05
p < 2.2e-16
FC = 4.73
p < 2.2e-16
FC = 2.31
LuCaP147-Residual
C
-10xlog10(FDR)
UP
DOWN
FDR < 0.05
FDR > 0.05
LuCaP147-Residual
INFLAMMATORY_RESPONSE
ALLOGRAFT_REJECTION
TNFA_SIGNALING_VIA_NFKB
INTERFERON_ALPHA_RESPONSE
INTERFERON_GAMMA_RESPONSE
IL6_JAK_STAT3_SIGNALING
IL2_STAT5_SIGNALING
COMPLEMENT
COAGULATION
P53_PATHWAY
APOPTOSIS
REACTIVE_OXYGEN_SPECIES_PATHWAY
DNA_REPAIR
UV_RESPONSE_UP
UV_RESPONSE_DN
MITOTIC_SPINDLE
E2F_TARGETS
G2M_CHECKPOINT
MYC_TARGETS_V1
MYC_TARGETS_V2
GLYCOLYSIS
OXIDATIVE_PHOSPHORYLATION
HYPOXIA
PEROXISOME
FATTY_ACID_METABOLISM
CHOLESTEROL_HOMEOSTASIS
XENOBIOTIC_METABOLISM
BILE_ACID_METABOLISM
HEME_METABOLISM
APICAL_JUNCTION
APICAL_SURFACE
EMT
ANGIOGENESIS
ADIPOGENESIS
MYOGENESIS
SPERMATOGENESIS
PANCREAS_BETA_CELLS
ANDROGEN_RESPONSE
ESTROGEN_RESPONSE_EARLY
ESTROGEN_RESPONSE_LATE
NOTCH_SIGNALING
WNT_BETA_CATENIN_SIGNALING
TGF_BETA_SIGNALING
HEDGEHOG_SIGNALING
MTOR1_SIGNALING
PI3K_AKT_MTOR_SIGNALING
KRAS_SIGNALING_UP
KRAS_SIGNALING_DN
UNFOLDED_PROTEIN_RESPONSE
PROTEIN_SECRETION
0 1 2 3 4 5 6 7
D
H3K27ac Differential Binding
LuCaP147-CR + dTRIM24-1 (post) vs LuCaP147-CR
-log10(padj)
Log2 Fold Change
MFHAS1
TUSC1
LINC01574
LMO4
MECOM
TMEM28
MIPOL1
KCNK15
PHLDA3
LOC283194
NETO1-DT
KCNN4
PKDCC
BRINP3
CSMD1
GULP1
GLYCAM1
LOC105375938
P2RY12
E
CNTR
dTRIM24-1 (post)
dTRIM24-2 (post)
TRIM24
Phospho-STAT3 (Ser727)
STAT3
N-CAD
TUB
LuCaP147-CR
F
LuCaP147-CR
CNTR n=9
dTRIM24-1 n=8
dTRIM24-2 n=8
Tumor Volume (mm3)
Days of Treatment (d)
p <0.0001
p <0.0001

Figure S18

**Supplementary Figure 18. A** UMAP plot displaying scRNA-Seq data from LUCaP147(-HS) hormone-sensitive sample, both pre-castration and LuCaP147-Residual after castration, treated with either a vehicle (CNTR) or dTRIM24-1 at the end of the treatment (refer to Fig. 5C). Upper panel shows the distribution of G1 and G2/M cell cycle phases. The lower panel shows the gene set enrichment scores for AR Response (middle panel) and the mTORC1 signaling (lower panel). **B** Corresponding violin plots displaying the single-sample gene set enrichment (ssGSEA) score for the AR Response and mTOR signaling for the indicated samples. Statistical significance is calculated using the Wilcon test. **C** Gene set enrichment analysis of the cell clusters identified in LuCaP147-HS and LuCaP147-Residual samples treated with either vehicle (CNTR) or dTRIM24-1 (see Fig. 7B). The dot plot illustrates the false discovery rate (FDR) associated with each cell cluster's gene enrichment scores, with red dots representing significant upregulation and blue dots indicating downregulation. **D** Volcano plot highlightingc key gene expression changes linked to H3K27ac marks in LuCaP147-CR tumors treated with dTRIM24-1 compared to those treated with the vehicle. **E** Immunoblot analysis of the indicated proteins was carried out on LuCaP147-CR tumors following castration and subsequent treatment with dTRIM24-1 (post), dTRIM24-2 (post), or the vehicle (CNTR) as indicated. **F** Growth kinetics of LuCaP147-CR in castrated NRG mice. The mice were treated intraperitoneally with either the vehicle or dTRIM24-1 and dTRIM24-2 at 5 mg/kg for three weeks, five days a week. The statistical significance of the results was tested using a two-way ANOVA test.
