## Supplemental Methods for "TRIM24 Degradation Counteracts Adaptation to Androgen Receptor Inhibition in Prostate Cancer"

### Supplementary Methods

#### Cytokine Array Profile

The Human XL Cytokine Array Kit (ARY022B R&D system) was used to analyze the cytokines, chemokines, and other soluble factors present in the LNCaP supernatant. The manufacturer's instructions were followed for this analysis. The LNCaP cells were cultured in charcoal-stripped serum (CSS) for three days and then supplemented with 10uM Enzalutamide alone or combined with 5uM of dTRIM24-1 or 5uM of dTRIM24-1. The supernatant from LNCaP cells cultured in CSS supplemented with 1nM DHT was used as a reference for the assay. For each array, 1 milliliter of the cell culture supernatant was used and left for 24 hours. Pixel density plots were then detected. The protein bands were visualized using the chemiluminescent detection reagent with the kit. Results were quantified using the Fusion Solo IV LBR system. The images were analyzed using Image Lab Software. The pixel density of each duplicated spot was measured, and the negative spot control signal was used as a reference value for the background. The average background signal was then subtracted from each spot, and the resulting signal was compared to the corresponding DHT-treated condition to calculate the average Fold Changes (FC) (Table S3).

#### Liquid chromatography-tandem mass spectrometry (LC-MS/MS)

Protein extraction and enzymatic digestion

Four biological replicates were prepared for each experimental condition by pelleting 2 x 106 cells. The pellets were washed twice in phosphate-buffered saline (PBS), flash-frozen, and stored at -80°C. Cell lysis and protein extraction were performed in 8M urea in 50 mM ammonium bicarbonate (ABC), sonicating the samples in a Bioruptor (Diagenode, 15 cycles, 30s on, 30s off, high mode). Proteins were reduced with 10 mM dithiothreitol for 20 minutes at room temperature and alkylated with 50 mM iodoacetamide for 30 minutes at room temperature. Sequential digestion was carried out in 8M urea in 50 mM ABC for 2 hours at room temperature with LysC (Wako Fujifilm, 1:100 w/w), after which the digestion buffer was diluted to final 2M urea with 50 mM ABC and trypsin (Promega, 1:100 w/w) was added for overnight digestion at room temperature. Digestion was halted by adding acetonitrile (ACN) to 2% and trifluoroacetic acid (TFA) to 0.3%, and the samples were cleared by centrifugation at maximum speed for 5 minutes. The digested peptides in the supernatants were purified with C18 StageTips (Rappsilber et al., 2007) and eluted with 80% ACN and 0.5% acetic acid. Finally, the elution buffer was removed by vacuum centrifugation, and purified peptides were dissolved in 2% ACN, 0.5% acetic acid, and 0.1% TFA for single-shot LC-MS/MS measurements.

### LC-MS/MS analysis

Peptides were separated on an EASY-nLC 1200 HPLC system (Thermo Fisher Scientific) coupled online via a nanoelectrospray source (Thermo Fisher Scientific) to a Q Exactive HF mass spectrometer (Thermo Fisher Scientific). Peptides were loaded in buffer A (0.1% formic acid) in a 75 µm inner diameter, 50 cm length column, in-house packed with ReproSil-Pur C18-AQ 1.9 µm resin (Dr. Maisch HPLC GmbH), and eluted over a 150-min linear gradient of 5 to 30% buffer B (80% ACN, 0.1% formic acid) at a flow rate of 250 nl/min. The Q Exactive HF was operated by the Xcalibur software (Thermo Scientific) in a data-dependent mode, with a survey scan range of 300-1,650 m/z, resolution of 60,000 at 200 m/z, maximum injection time of 20 ms, and AGC target of 3e6.

The top 10 most abundant ions with charges 2 to 5 were isolated with a 1.8 m/z isolation window and fragmented by higher-energy collisional dissociation (HCD) with a normalized collision energy 27. MS/MS spectra were acquired with a resolution of 15,000 at 200 m/z, a maximum injection time of 55 ms, and an AGC target of 1e5. A 30 s dynamic exclusion was employed to avoid repeated sequencing.

### LC-MS/MS data analysis

MS raw files were processed using the MaxQuant software v.1.6.7.0 (Cox & Mann, 2008). The Andromeda search engine (Cox *et al.*, 2011) was employed to search spectra against the Human UniProt database (June 2019) and a common contaminants database (247 entries) to identify peptides and proteins with a false discovery rate of < 1%. Enzyme specificity was set as “Trypsin/P” with a maximum of 2 missed cleavages and a minimum length of 7 amino acids. N-terminal protein acetylation and methionine oxidation were set as variable modifications, and cysteine carbamidomethylation was set as a fixed modification. Match between runs was enabled to transfer identifications across samples based on mass and normalized retention time, with a matching time window of 0.7 min and an alignment time window of 20 min. Label-free protein quantification (LFQ) was calculated by the MaxLFQ algorithm (Cox *et al.*, 2014) with a minimum peptide ratio count of 1. Data analysis was performed using the Perseus software v.1.6.2.3 (Tyanova et al., 2016). The proteinGroups.txt table was pre-processed by removing proteins only identified by site, reverse hits, and potential contaminants. After log2 transformation of LFQ intensities, biological replicates of each experimental condition were grouped, and proteins were filtered for a minimum of 3 valid values in at least one group. Missing data points were replaced by imputation from a normal distribution with 0.3 widths and 1.8 downshift, and a two-sided two-sample t-test (0.05 FDR, 250 randomizations) was used to identify significant changes in protein intensity between groups. A list of TRIM24 targets was used to evaluate their distribution in the corresponding volcano plots.

#### Immunohistochemistry on a TMA of primary and matched castration-resistant human prostate cancer

The abundance of TRIM24 protein was studied in tissue samples of matched primary and castration-resistant prostate cancer (CRPC) patients from the University of Basel using immunohistochemical staining. The study was conducted retrospectively, with samples obtained under the approval of the Ethics Committee of Northwestern and Central Switzerland (EKNZ) with the approval numbers EK/1311 and 2015/228. The benign prostate samples were obtained from three patients using tumor-free prostate core needle biopsies. The prostate cancer biopsies were taken during routine clinical treatment. They were selected based on the following criteria: (a) diagnosed with histological PCa, (b) tumor-containing biopsies available at HN and CR state, and (c) sufficient quality and amount of material, as evaluated by experienced pathologists (LB and KM). The criteria for castration resistance were defined as either biochemical progression or clinical progression. A total of 112 matched HN/CR tissue specimens were included in the study, but only 35 high-quality matched samples remained after sectioning due to tissue loss. The Bond-III automated staining system was utilized to analyze slides with 8 µm sections for TRIM24 and phosphorylated STAT3 (Tyr705) immunohistochemistry (IHC) using commercially available reagents. For antigen retrieval, the slides were incubated for 60 minutes in citrate buffer at pH 9 at 98°C for TRIM24 and 60 minutes in EDTA buffer at pH 9 at 98°C for phosphorylated STAT3 (Tyr705). Then, slides were incubated with a rabbit polyclonal antibody against TRIM24 (Cat HPA043495 from Sigma-Aldrich) at a dilution of 1:400 and a rabbit polyclonal antibody against phospho-STAT3(Tyr705). Detection was performed using the Detection Refine DAB kit (Leica). Aperio ImageScope was used to evaluate immunohistochemical staining as a percentage of tumor cells with nuclear positivity for TRIM24.

#### Immunohistochemistry on a TMA human primary prostate cancer and matched metastatic samples

Immunohistochemistry was performed on a TMA of human prostate cancer, including primary tumors and matched nodal metastases. The TMA (catalog number TA3069) was obtained from Tristar Technology Group (Rockville, MD, USA), and contained 15 matched specimens of primary and metastatic tissues. Immunohistochemistry for TRIM24 was conducted using the protocols previously reported (see Immunohistochemistry on a TMA of human prostate cancer primary and matched castration-resistant samples).

#### Immunohistochemistry on human de novo prostate cancer metastasis

Prostate biopsy specimens were collected at the Oncology Institute of Southern Switzerland from patients who presented with de novo metastatic hormone-sensitive prostate cancer (mHSPC) and began first-line therapy between 2009 and 2021. Each patient either provided informed consent for data collection or was deceased by the time of data analysis. This study adhered to a retrospective biological sample collection protocol approved by the local ethics committee and complied with the 1964 Helsinki Declaration and its later amendments or comparable ethical standards. Clinical follow-up data were extracted from electronic medical records, and radiographic progression-free survival (rPFS) was calculated for all eligible patients. Patients included in the study met all of the following criteria: (a) Males aged 21 years or older; (b) Histologically confirmed prostate cancer from prostate biopsy; (c) International Society of Urological Pathology (ISUP) Grade Group 4 or 5; (d) Evidence of metastatic disease at diagnosis; (e) High-volume disease as defined by the CHAARTED criteria; (f) Received first-line treatment with either androgen deprivation therapy (ADT) monotherapy or ADT in combination with docetaxel or an androgen receptor pathway inhibitor (ARPI); (g) Classified as either a 'poor responder' (rPFS ≤ 9 months) or an 'exceptional responder' (rPFS > 38 months) following the initiation of ADT. Patients were excluded if they met any of the following criteria: (a) Primary pure small cell carcinoma of the prostate; (b) Concurrent malignancies requiring cytotoxic therapy; (c) Missing clinical follow-up data. Immunohistochemistry for TRIM24 was performed using the same conditions previously reported (see Immunohistochemistry on TMA of human matched Primary and Castration-Resistance Prostate Cancer (CRPC) samples. The IHC for TRIM24 was performed using the same conditions previously reported (see Immunohistochemistry on a TMA of human prostate cancer primary and matched castration-resistant samples). For the H&E staining, the FFEP human tissue was cut into 4 μm sections for Hematoxylin and eosin (H&E) staining (Diapath, C0303) and (Diapath, C0363), respectively. Once dried, the sections were treated with OTTIX plus solution (Diapath, X0076) and OTTIX shaper solution (Diapath, X0096) to dewax and rehydrate the sections. Tissues were incubated for 5 min in Hematoxylin followed by 3 min in Eosin solution. Once mounted, the slides were acquired with an Aperio AT2 Slide scanner using Imagescope software (Leica Biosystems).

**Survival analysis**

Gene expression (FPKM) and clinical outcome data (overall survival) for the SU2C prostate cancer cohort were obtained from publicly available resources. Samples with complete outcome information were retained. To compute the TRIM24-associated signature score, the gene set was evaluated using single-sample gene set enrichment analysis (ssGSEA) via the GSVA package (R, version 4.3). Each sample was assigned a ssGSEA enrichment score, which was subsequently used to stratify patients into groups (e.g., upper vs. lower quartile) for survival comparisons. Survival analyses were performed using the R packages *survival* and *survminer*. Kaplan–Meier survival curves were generated via the *survfit* function, and significance was assessed by the log-rank test. Multivariable Cox proportional hazards regression (*coxph*) was used to evaluate the prognostic impact of the signature while adjusting for relevant clinical covariates (including Gleason score, AR amplification, and TP53 or RB1 alteration status). Hazard ratios and 95% confidence intervals were summarized, and proportional hazard assumptions were checked. Subgroup analyses (e.g., patients "Naive" to abiraterone/enzalutamide vs. "Exposed") were conducted by stratifying the dataset prior to Kaplan–Meier and Cox modeling. All analyses were performed in R, and significance was defined at $p < 0.05$ unless otherwise specified.

**Correlation analysis between TRIM24 signature and mutation**

For downstream correlation analyses, somatic mutations were filtered to retain coding or splice-altering events (e.g., frameshift, in-frame, missense, nonsense). A binary gene-by-sample mutation matrix was constructed, where rows represented individual genes and columns represented samples, coded "1" for the presence of a mutation in that gene and "0" for wild-type status. According to clinical annotations, these samples were previously categorized into primary or advanced disease states (castration-resistant or neuroendocrine). A TRIM24 signature score was calculated for each sample using single-sample gene set enrichment analysis (ssGSEA), and these scores were correlated with mutation status.
Specifically, the distribution of TRIM24 scores in mutated versus wild-type samples for each gene was compared using a Wilcoxon rank-sum test, while point-biserial, Pearson and Spearman correlations were separately evaluated to capture both linear and nonparametric relationships. Genes with mutations in fewer than five samples per cohort were excluded to ensure statistical robustness. Multiple-testing considerations (e.g., false discovery rate) were applied to control for Type I error across the large panel of genes. Comparisons were performed independently within primary and CRPC/NEPC cohorts to assess stage-specific associations. Results were then ranked by effect size and significance, and selected gene-level comparisons were visualized using ggplot2. This approach provided an integrated perspective on the relationship between mutation burden and TRIM24 signature activity across different disease states.

**Correlation analysis between TRIM24 signature and copy number alterations**

Copy number data were obtained for both primary and advanced (CRPC/NEPC) prostate tumors, and genomic segments were categorized into discrete copy number states ranging from homozygous deletion (–2) to high-level amplification (+2). To correlate the copy number status at each gene with the TRIM24 ssGSEA signature, a repeated “leave-half-out” strategy was employed. In brief, each cohort (primary or CRPC/NEPC) was subsampled 10 times without replacement, and for each gene, Pearson’s correlation was computed between the copy number values and the corresponding TRIM24 ssGSEA scores. The resulting correlation coefficients and p-values from each subsample were averaged to obtain robust estimates of association. Multiple-testing correction (e.g., FDR) was performed to control for false positives and genes with insufficient CN events (<5 occurrences of either homozygous deletion or high-level amplification) were excluded from final analyses. Visualizations, including monodimensional scatter plots of correlation coefficients versus statistical significance, were generated with ggplot2, highlighting select genes of interest (e.g., MYC, PTEN, TP53, RB1) to illustrate potentially clinically relevant associations.

**Radar plots**

RNA-sequencing data from LNCaP-derived cell lines harboring double knockout of TP53 and RB1 and from LuCaP-147 xenografts (hormone-sensitive and castration-resistant) were analyzed to characterize their transcriptional profiles and classify them into clinically relevant prostate cancer subtypes. To achieve this, we applied Nearest Template Prediction (NTP) using the CMScaller R package. NTP employs predefined gene expression signatures to determine how closely a sample’s expression pattern aligns with established subtype templates. In this study, four predefined subtype-specific gene-expression signatures were used, corresponding to Androgen Receptor (AR), Stem Cell–Like (SCL), Neuroendocrine (NE), and WNT signaling, drawn from previously published literature. The *ntp* function was run with 10,000 permutations, providing both the subtype assignments and statistical confidence estimates (p-values and false discovery rates, FDRs).

Following NTP classification, the raw distance metrics were transformed into similarity scores (1 – distance) to better visualize the relative relationship of each sample to the four subtypes. A radar (spider) plot was generated, where each axis represented one of the four subtype dimensions, and the polygon defined by connecting these values depicted the sample’s overall signature profile. Samples were color-coded by their condition, allowing intuitive visual comparison among pre-castration, post-castration, and post-castration with enzalutamide groups.

For statistical assessment of subtype differences, we performed one-way ANOVA tests on the similarity scores derived from the NTP results for each subtype. The different conditions (e.g., pre-castration, post-castration, and post-castration with enzalutamide for the LNCaP model, and hormone-sensitive versus castration-resistant for LuCaP-147) were used as the independent variable, and each subtype’s NTP-derived similarity score served as the dependent variable. ANOVA p-values were derived from F-tests assessing whether the mean similarity scores differed among the three conditions. Dimensions with $p < 0.05$ were considered significantly different across conditions. These statistically significant dimensions were marked on the radar chart by adding an asterisk (*) to the corresponding axis label. This integrated approach provided a clear, quantitative visualization of how each sample's transcriptional profile aligned with distinct CRPC subtypes across varying experimental and therapeutic contexts.

**Exome-sequencing data processing and mutation calling**

Whole-exome sequencing (WES) was performed on frozen tumor fragments from the LuCaP147-HS and LuCaP147-CR patient-derived xenograft (PDX) models. WES was conducted on biological duplicates for each sample. Genomic DNA extraction was carried out using the DNAeasy Blood and Tissue Kit (69506 Qiagen) according to the manufacturer's instructions. Library preparation for next-generation sequencing was performed using Twist EF Library Prep 2.0 + UDIs and Standard Hyb (p/n 104384; Twist Bioscience) and Twist Comprehensive Exome (p/n 103697; Twist Bioscience) according to manufacturer's protocol. Quality controls were performed on Bioanalyzer 2100 (Agilent Technologies, Santa Clara, CA, USA) and Qubit V4 (Invitrogen Thermo Fisher Scientific, Waltham, MA, USA). Next-generation sequencing was performed on NextSeq2000 (Illumina, San Diego, CA, USA) with the P2 reagents kit V3 (200 cycles; Illumina). Samples were processed starting from stranded, paired-ended 108bp-long sequencing reads.

To address mouse contamination inherent to PDX samples, raw reads were first aligned to a hybrid genome composed of the human (GRCh38) and mouse (GRCm39) reference sequences using BWA-MEM with default parameters. The resulting BAM files were sorted, and all reads aligning to mouse chromosomes were removed with *samtools*. The filtered human-only reads were converted back to FASTQ and subjected to a second alignment step against the GRCh38 reference genome. This iterative approach ensured minimal contribution of mouse DNA to subsequent analyses. Following alignment, read group information (including sample name, library, platform, and flowcell ID) was assigned during BAM file generation to facilitate downstream processing.

After re-alignment against GRCh38, all human-specific reads were processed using Genome Analysis Toolkit (GATK, version 4.4.0) Best Practices to generate high-quality BAM files. The unmapped BAM files were merged with their corresponding mapped reads to retain complete read information, and the resulting merged BAM was coordinate sorted. NM and UQ tags were corrected to accurately reflect mismatch counts and unique mapping qualities. Duplicate reads were marked to produce duplication metrics and prevent inflation of variant allele frequencies. Base Quality Score Recalibration (BQSR) was then conducted in two steps using dbSNP and *Mills_and_1000G_gold_standard indels* as known variant resources. First, the recalibration model was generated with BaseRecalibrator, then recalibrated scores were applied with *ApplyBQSR*. We called variants using GATK Mutect2 restricted to exonic targets (supplemented by a panel of normals and a germline resource from *gnomAD*) to improve detection specificity. Finally, the resulting variant call files were indexed and annotated with ANNOVAR, referencing multiple databases such as RefGene, cytoBand, avsnp147, ClinVar, and COSMIC. Throughout, quality control metrics (including total read depth and percentage of duplicate reads) were monitored to ensure reliable coverage and data integrity for subsequent downstream analyses.

**ChIP-sequencing data processing and peak calling**

ChIP-sequencing reads from H3K27ac and TRIM24 immunoprecipitates and matched input controls were processed using the following computational pipeline. First, raw single-end FASTQ files were assessed for quality metrics (e.g., adapter presence, per-base quality) to ensure only high-quality reads were retained. Bowtie2 (version 2.5.4) was then employed to align reads to the GRCh38 reference genome using single-end mode and default sensitivity settings. SAM files were converted to coordinate-sorted BAM files with Samtools, and Picard MarkDuplicates (version 3.3.0) was used to remove redundant reads and generate duplication metrics. Subsequently, deduplicated BAM files were indexed for downstream processing. Peak calling was performed with MACS3 (version 3.0.2), with the appropriate input library for each immunoprecipitate as a control and specifying parameters (e.g., *"-g hs*" for human genome size) to identify significantly enriched regions. For TRIM24 ChIP-seq, significantly enriched regions were identified at a false discovery rate threshold of $q < 0.1$, whereas for H3K27ac ChIP-seq, $q < 0.01$ was used. Genome-wide coverage tracks were first generated from the deduplicated ChIP-seq alignments using bamCoverage (deepTools, version 3.5.5) with read-per-genomic-content (RPGC) normalization. A bin size of 10 bp and an effective genome size of 2,913,022,398 (GRCh38) were specified to ensure consistent resolution and comparability. These primary bigWig files were then merged across biological replicates using bigwigCompare (also from deepTools, version 3.5.5), applying the "mean" operation to create a single averaged coverage track per condition. By combining replicates in this manner, coverage noise and technical variation were reduced while preserving representative signal intensity. The resulting merged bigWig files were subsequently used for integrative analyses, such as genome browser visualization and quantitative comparisons of H3K27ac- or TRIM24-immunoprecipitated chromatin relative to matched input controls across LuCaP-147 and LuCaP-147 CR models. To compare the peak intensities between different models, we used the *computeMatrix* module (deepTools, version 3.5.5) in "reference-point" mode to measure read coverage from bigWig files in a 3-kb window (±3 kb) centered on each merged peak of interest. The resulting matrix was then visualized with *plotProfile* to generate average signal profiles, directly comparing enrichment patterns between conditions.

#### Motif analysis in ChIP–seq peaks

Motif analyses were performed using HOMER (*findMotifsGenome.pl*), applying the narrowPeak intervals output by MACS3 as input regions. Each peak set was examined against the GRCh38 reference genome using a 500 bp window centered on the peak summit. HOMER's default motif databases and statistical thresholds were used to identify both known and de novo transcription factor binding motifs enriched in each peak set. Chi-square statistical test was used to show the significant relative enrichment of transcription factor motifs. The resulting motif enrichments were compiled for downstream comparative analyses.

#### Differential binding analysis and gene-set enrichment in ChIP–seq

Reads mapping to the ChIP-sequencing peaks of genomic regions were quantified using featureCounts (*subread* package in R), specifying single-end mode, the hg38 annotation (annot.inbuilt="hg38"), and disallowing multi-mapping reads to ensure robust count estimation. To identify differentially enriched sites across conditions, the resulting count matrix was imported into the edgeR workflow. Library size normalization was performed with the TMM method (*calcNormFactors*), and dispersion parameters were estimated (*estimateDisp*) prior to fitting a generalized linear model (*glmQLFit*). The contrast of interest was tested with a quasi-likelihood F-test (glmQLFTest) to detect significantly differentially bound regions. Multiple-testing correction was applied using the false discovery rate (FDR) approach, and significant sites were annotated by mapping Entrez gene identifiers to gene symbols (org.Hs.eg.db).
Gene set enrichment analysis (GSEA) was conducted with *fgsea* and *clusterProfiler*. A rank-ordered gene list was derived by sorting sites according to log2FC, and these were mapped to hallmark gene sets from the Molecular Signatures Database (MSigDB). Statistically significant enrichment (FDR < 0.1) was visualized via dot plots generated in *ggplot2*. This integrated approach enabled a comprehensive assessment of differential occupancy and functional pathway associations.

#### Gene Expression Quantification of Bulk RNA-Seq Data

The overall quality of sequencing reads was evaluated using FastQC (v.0.11.9). Sequence alignments to the reference human genome (GRCh38.p13) were performed using STAR (v.2.7.1b) in two-pass mode, to significantly increase sensitivity to novel splice junctions compared to the regular single mapping. In the two-pass mapping procedure, reads are mapped twice: in the first pass, the novel junctions are detected and inserted into the genome indices; in the second pass, all reads are re-mapped using annotated (from the GTF file) and novel (detected in the first pass) junctions. Gene expression was quantified at the gene level in the second pass by using the comprehensive annotations made available by Gencode (v37 GTF File). Strand-specific information was not maintained to avoid technical differences between stranded and unstranded libraries. Samples were adjusted for library size and normalized with the variance stabilizing transformation (vst) in the R statistical environment using DESeq2 (v1.34.0) pipeline. When performing differential expression analysis between groups, we applied the embedded *IndependentFiltering* procedure to exclude genes that were not expressed at appreciable levels in most of the samples considered.

**Gene-Set Enrichment Analysis**

All GSEAs were performed using *Custerprofiler*(v4.2.2) package (GSEA, eps=1e-50). Gene-set collections were retrieved from the Molecular Signature Database (MSigDB) or previous publications (AR/NE-Score). P values were corrected for multiple testing using the false discovery rate (FDR) procedure, with the significance threshold set to 0.05. In addition, GSEA significance was logarithmically transformed in the form of −10log10(p-adjusted), with a bold intercept (x = 13.01) indicating the FDR threshold depicted in the corresponding plots. Over-representation analyses (ORA) were performed using *Custerprofilerc*(v4.2.2) package (*enrichR*).

**Gene Expression Quantification of scRNA-Seq Data**

*Fastq* files were generated by demultiplexing raw data using *Cellranger mkref* (v7.1.0). We generated a custom genome with *Cellranger*, using the same reference (GRCh38.p13) and annotations (Gencode v37) we used for STAR when performing bulk RNA-seq analysis. To discriminate between human and murine cells that may infiltrate the tumors in the in vivo setting, we created a Mouse-Human reference, by creating a hybrid genome (GRCh38.p13 + GRCm39) and hybrid gene-annotations (Gencode v37 and M26, for human and mouse genes, respectively). Such reference has been used for reads alignment, performed using *Cellranger*. To avoid conflicts, a suffix has been added to mouse genomic coordinates (i.e., chr1M, chr2M, etc.). Subsequently, *Cellranger* was used to quantify gene expression in the form of an h5-filtered matrix where Ensembl gene IDs were used as identifiers.

**Data filtering and clustering of scRNA-Seq Data**

Expression quantification files were imported into R statistical environment using Seurat (4.3.0) package. Individual cells from the data matrix were discarded by using a two-filtering procedure: first, we aimed at detecting transcriptional outliers, and second, we looked for putative doublets, which we also discarded.

We computed per-cell quality control metrics using *scater* (v1.22.0). The total amount of mitochondrial and ribosomal gene expression was quantified for both human and mouse cells. The number of genes being detected per cell, the total amount of reads per cell, and the mitochondrial and ribosomal fraction of the transcriptome were used to determine the skewness-adjusted multivariate outlyingness for each cell (robustbase v0.95-0). Outliers were detected by median absolute deviation and removed at both tails.

Counts were then normalized (Seurat::NormalizeData, method = LogNormalize, scale.factor = 1000) and the top 2000 most variable features were selected (Seurat::FindVariableFeatures, method = vst). Data were then scaled (Seurat::ScaleData) and PCA was performed up to the top 50 components (Seurat::RunPCA). Subsequently, we identified and eliminated putative doublets using DoubletFinder (v2.0.3). Having identified outliers and doublets, we removed them from the original count data and repeated the preprocessing step (i.e., normalization, scaling, and dimensionality reduction). We then proceeded to determine the *k*-nearest neighbors of each cell and the construction of a shared nearest-neighbor (SNN) graph (Seurat::FindNeighbors), then we identified clusters using the SNN modularity optimization-based clustering algorithm (Seurat::FindClusters, resolution = 0.5). Finally, we performed Umap dimensionality reduction on the first 30 PCs, annotated the previously identified clusters, and generated plots accordingly.

**Species Discrimination and Downstream Analysis**

Following dimensional reduction (PCA and UMAP) and clustering in Seurat (version 4.3.0), each cluster's distinguishing genes were identified using the *FindMarkers* function with a receiver operating characteristic (ROC) test, focusing on transcripts highly enriched in that cluster relative to all others. Clusters whose top markers corresponded predominantly to annotated human genes were attributed to human tumor cells, whereas clusters enriched in mouse genes were classified as murine. Any cluster showing a mixed profile (i.e., no clear majority of human or mouse markers) was excluded from subsequent analyses. This approach ensured robust discrimination of tumor-derived (human) and stromal or infiltrating (mouse) cell populations in the dataset.

**Cell type annotation (mouse subset)**

To further confirm the assignment of mouse-derived cells, the subset of murine gene-expression profiles was converted into a SingleCellExperiment object and annotated with SingleR (version v1.8.1). Two reference panels were used: ImmGenData and MouseRNAseqData (from the *celldex* package), each containing well-characterized transcriptomic profiles of various mouse cell types. SingleR was run in both “single-cell” mode and “cluster” mode, the latter assigning labels based on the averaged gene expression of each Seurat cluster. Clusters were then annotated according to the closest matching reference cell type. Throughout these steps, data integrity and cluster assignments were verified via known marker genes (e.g., those enriched in immune vs. stromal vs. parenchymal compartments). This systematic workflow of hybrid genome alignment, separation into human vs. mouse compartments, and reference-based cell identity assignment ensured an accurate distinction between tumor cells of human origin and infiltrating murine cells in the xenograft samples.

**Integration of Multiple Single-Cell Datasets**

Performing an integrated analysis of multiple single-cell datasets presents distinct challenges in the field. One significant challenge involves accurately identifying cell populations that are shared across these datasets within conventional workflows. To address this issue, Seurat v4 introduces a range of techniques designed to align shared cell populations across datasets. These techniques initially identify pairs of cells from different datasets that correspond to matched biological states, known as "anchors." The identified anchors serve a dual purpose: they facilitate the correction of technical variations between datasets, such as batch effects, and enable comparative analysis of scRNA-seq data across different experimental conditions. The first step was to perform a uniform dataset normalization using the *SCT* function implemented in Seurat. This step includes splitting the dataset into a list of separate Seurat objects, and the subsequent normalization and identification of variable features for each dataset independently. Last, we selected features that are repeatedly variable across datasets for the integration (Seurat::*SelectIntegrationFeatures*). Then, we then identify anchors using the *FindIntegrationAnchors*() function, which takes a list of Seurat objects as input, and use these anchors to integrate the two datasets with *IntegrateData*(). We perform subsequent downstream analysis on the corrected data: the original unmodified data still resides in the 'RNA' assay in the integrated Seurat object.

**Dealing with Drop-Out Events**

Drop-out events are very frequent in the single-cell experiment performed using 10x Chromium technology. When drop-out occurs, the absence of data for a particular gene in a specific cell can introduce biases and distort the overall picture of gene expression patterns. This can lead to incorrect interpretations of the data, especially when trying to identify rare cell types or subtle differences between cells. To address these issues, we applied Markov Affinity-based Graph Imputation of Cells (*MAGIC* algorithm, RMagic v2.0.3)

**Differential Expression Analysis and Gene-Set Enrichment**

Differential expression was performed between cell clusters subjected to different treatment conditions (Seurat::*Findmarkers*) using a hurdle model tailored to scRNA-seq data (MAST method). Genes were subsequently ranked for $\log_2$ FC, and the *Camera* algorithm (pre-ranked) was used to determine gene-set enrichments for each comparison. Cell-specific gene-set enrichments were determined using single-sample GSEA (Seurat::*AddModuleScore*), computed using gene-expression values of each cell following *RMagic* imputation.

**Identification of Cell Cycle Phase**

According to Seurat's annotations, we retrieved the list of cell cycle markers and subdivided it into G2/M and S phase markers. We then used this information to infer the cell cycle phase in our samples (Seurat::*CellCycleScoring*). This function calculates a cell cycle score for each cell by comparing the expression of cell cycle marker genes against a reference gene list These scores represent the relative position of each cell in the cycle, with higher scores indicating cells in S or G2/M phase and lower scores indicating cells in G1 phase.

**Gene Signatures**

The gene signatures utilized are derived primarily from the Molecular Signature Database (MSigDB) and the scientific literature. The Molecular Signature Database provides a comprehensive collection of gene sets that represent various biological processes, pathways, and functional annotations. Specifically, it includes hallmark gene sets (H), which capture fundamental biological processes and signaling pathways that are commonly dysregulated in different diseases or conditions. Additionally, the curated gene sets in MSigDB (C2) encompass specific biological knowledge curated from diverse sources. Also, the Gene Ontology database (C5) has been used to enrich for specific biological processes or molecular functions.

The *Progression Signature* was derived from single-cell RNA-sequencing data after differential expression analysis (Seurat::*FindMarkers*, method=MAST) between the regrowth condition and the pre-castration sample. Differentially expressed genes were ordered by log2FC and the first 200 hints were used to create the signature.

**Pathway Scoring and Fold-Change Computation**

Pathway enrichment scores were computed for each cell using Seurat's *AddModuleScore* function, which aggregates the expression values of a predefined gene set and applies background correction. Violin plots were then used to visualize these enrichment scores across different sample groups. To facilitate ratio-based comparisons, a small constant (equal to the absolute value of the minimum score plus a small offset, e.g., 1e–5) was added to all scores where necessary, ensuring they remained positive. The median score was then calculated for each condition, and fold changes were determined by taking the ratio of medians across pairwise group comparisons. For gene expression values, fold changes were computed by dividing the median expression of one group by that of another. In both cases, the resulting fold-change values were annotated directly onto violin plots, providing a straightforward comparison of relative pathway activity or gene expression levels. Statistical significance was assessed using the Wilcoxon rank-sum test (e.g., via *stat_compare_means*), with *p*-values displayed alongside the fold-change annotations.

**Chemistry analytic methods**

dTRIM24-1 and dTRIM24-2 were obtained by fee-for-service. The synthesis for dTRIM24-1 has been described in a previous publication[26], while the synthesis for dTRIM24-2 is detailed in Patent No. US 10,702,504 B2: Degradation of tripartite motif- containing protein 24 (TRIM24) by conjugation of TRIM24 inhibitors with E3 ligase ligand and methods of use, applied by the Dana-Farber Cancer Institute in Boston. Assayed compounds were tested as TFA salts, and purities of assayed compounds were in all cases greater than 95%, as determined by reverse-phase UPLC analysis (Waters Acquity UPLC/MS system (Waters PDA eλ Detector, QDa Detector, Sample manager - FL, Binary Solvent Manager) using Acquity UPLC® BEH C18 column (2.1 x 50 mm, 1.7 μm particle size): solvent gradient = 85% A at 0 min, 1% A at 1.7 min; solvent A = 0.1% formic acid in water; solvent B = 0.1% formic acid in Acetonitrile; flow rate: 0.6 mL/min). NMR spectra were acquired on a 500 MHz Bruker Avance III spectrometer, operating at the denoted spectrometer frequency in MHz for the specified nucleus. Unless otherwise noted, all experiments were acquired at 298.0 K with a calibrated Bruker Variable Temperature Controller. The chemical shifts are reported in parts per million (ppm), and coupling constants (J) are given in Hertz (Hz). 1H NMR spectra are reported with the solvent resonance as the reference unless noted otherwise (CDCl3 at 7.26 ppm, CD3OD at 3.31 ppm, DMSO-d6 at 2.50 ppm). Peaks are reported as (s = singlet, d = doublet, t = triplet, q = quartet, m = multiplet or unresolved, br = broad signal, coupling constant(s) in Hz, integration).

**dTRIM24-1 compound characterization**

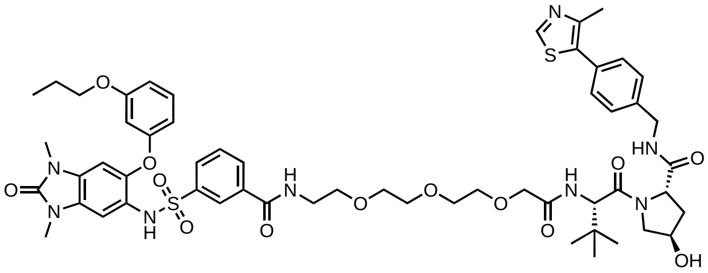

**$^{1}$H NMR** (500 MHz, DMSO-$d_6$) δ = 9.80 (s, 1H), 8.96 (s, 1H), 8.66 (t, *J* = 5.6 Hz, 1H), 8.58 (t, *J* = 6.1 Hz, 1H), 8.20 (t, *J* = 1.8 Hz, 1H), 7.99 (dt, *J* = 7.8, 1.4 Hz, 1H), 7.76 (ddd, *J* = 7.8, 2.0, 1.1 Hz, 1H), 7.47 (t, *J* = 7.8 Hz, 1H), 7.42 (d, *J* = 9.7 Hz, 1H), 7.39 (s, 4H), 7.10 – 7.00 (m, 2H), 6.71 (s, 1H), 6.54 (ddd, *J* = 8.3, 2.3, 0.8 Hz, 1H), 6.11 (ddd, *J* = 8.2, 2.3, 0.9 Hz, 1H), 6.04 (t, *J* = 2.3 Hz, 1H), 5.15 (d, *J* = 3.6 Hz, 1H), 4.56 (d, *J* = 9.6 Hz, 1H), 4.44 (t, *J* = 8.2 Hz, 1H), 4.41 – 4.33 (m, 2H), 4.25 (dd, *J* = 15.8, 5.7 Hz, 1H), 3.95 (s, 2H), 3.79 (t, *J* = 6.5 Hz, 2H), 3.67 (dd, *J* = 10.7, 4.0 Hz, 1H), 3.63 – 3.52 (m, 9H), 3.50 (t, *J* = 6.0 Hz, 2H), 3.39 (q, *J* = 6.0 Hz, 2H), 3.28 (s, 3H), 3.18 (s, 3H), 2.43 (s, 3H), 2.06 (dd, *J* = 12.9, 7.6 Hz, 1H), 1.90 (ddd, *J* = 13.0, 8.8, 4.5 Hz, 1H), 1.67 (h, *J* = 7.1 Hz, 2H), 0.96 – 0.91 (m, 12H). **$^{13}$C NMR** (126 MHz, DMSO-$d_6$) δ = 172.2, 169.6, 169.1, 165.2, 160.1, 159.0, 154.6, 151.9, 148.2, 145.3, 141.3, 139.9, 135.2, 131.6, 131.2, 130.3, 130.2, 129.7, 129.4, 129.2, 128.8, 127.9, 126.4, 126.3, 109.4, 109.0, 107.6, 103.9, 100.9, 70.9, 70.3, 70.1, 70.1, 69.4, 69.3, 67.4, 59.2, 57.0, 56.2, 36.2, 27.6, 27.5, 26.6, 22.4, 16.4, 10.9.

**LC-MS:** $C_{55}H_{68}N_8O_{13}S_2$ 1112.43; found $[M+H]^+$ 1113.31.

**dTRIM24-1 $^{1}$H NMR** (500 MHz, DMSO-$d_6$)

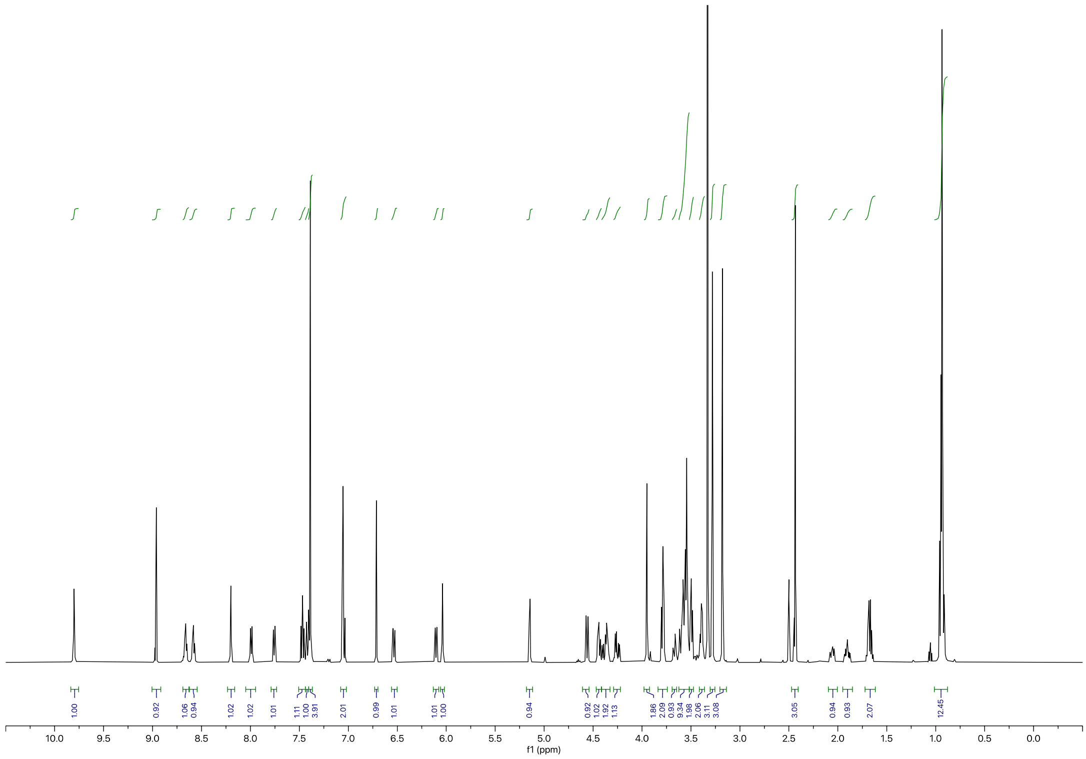


**dTRIM24-1 $^{13}$C NMR** (126 MHz, DMSO-d$_6$)

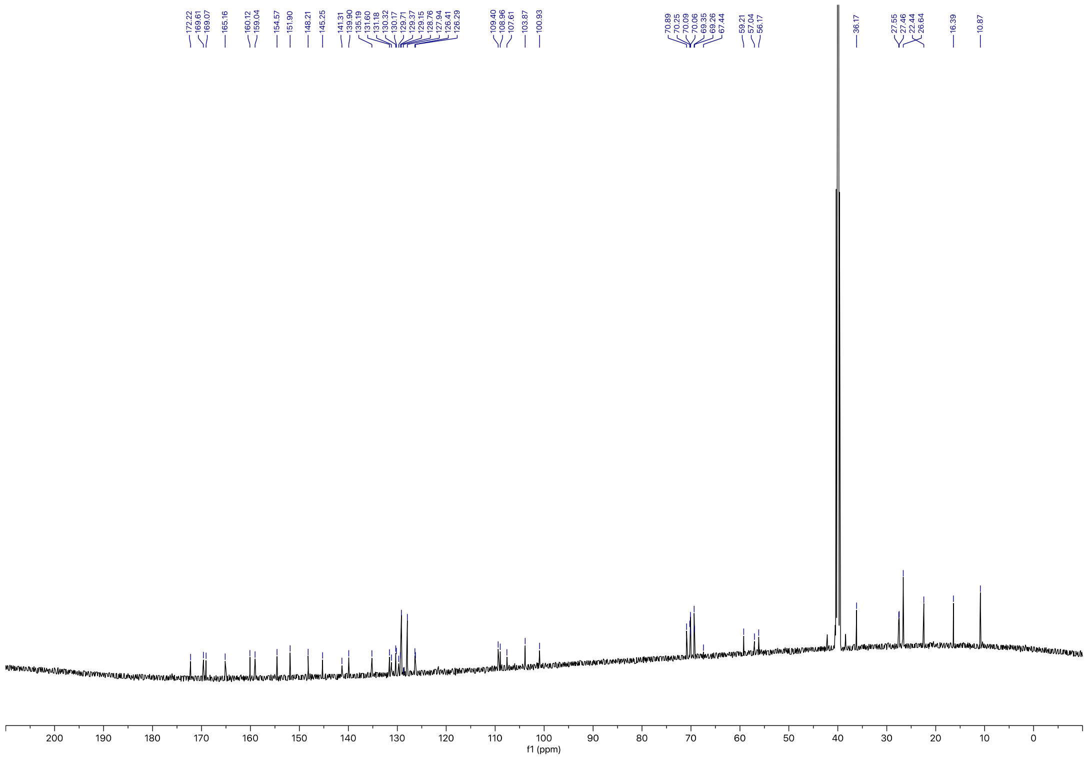

**dTRIM24-2 compound characterization**

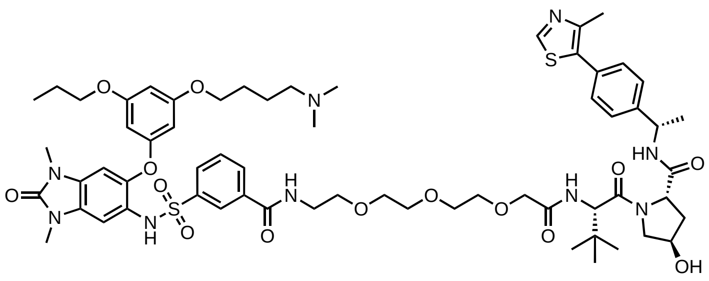

**$^{1}$H NMR** (500 MHz, DMSO-$d_6$) δ = 8.98 (s, 1H), 8.74 (t, *J* = 5.6 Hz, 1H), 8.46 (d, *J* = 7.7 Hz, 1H), 8.25 (t, *J* = 1.8 Hz, 1H), 8.05 (dt, *J* = 7.9, 1.3 Hz, 1H), 7.76 (ddd, *J* = 7.8, 2.0, 1.1 Hz, 1H), 7.49 (t, *J* = 7.9 Hz, 1H), 7.43 (dt, *J* = 7.4, 1.9 Hz, 2H), 7.40 – 7.34 (m, 3H), 7.02 (s, 1H), 6.75 (s, 1H), 6.11 (t, *J* = 2.1 Hz, 1H), 5.67 (d, *J* = 2.2 Hz, 1H), 5.16 (s, 1H), 4.90 (p, *J* = 7.0 Hz, 1H), 4.55 (d, *J* = 9.6 Hz, 1H), 4.45 (t, *J* = 8.2 Hz, 1H), 4.29 (s, 1H), 3.95 (s, 2H), 3.84 (d, *J* = 5.6 Hz, 2H), 3.78 (t, *J* = 6.5 Hz, 2H), 3.63 – 3.51 (m, 14H), 3.46 – 3.39 (m, 4H), 3.28 (s, 3H), 3.20 (s, 3H), 2.80 – 2.70 (m, 2H), 2.52 – 2.49 (m, 6H), 2.46 (s, 3H), 2.06 (dd, *J* = 12.9, 7.7 Hz, 1H), 1.78 (ddd, *J* = 13.0, 8.8, 4.6 Hz, 1H), 1.67 (q, *J* = 5.8 Hz, 4H), 1.37 (d, *J* = 7.0 Hz, 3H), 0.99 – 0.91 (m, 12H). **$^{13}$C NMR** (126 MHz, DMSO-$d_6$) δ = 170.9, 169.5, 169.0, 165.1, 160.8, 160.6, 159.8, 154.6, 152.0, 148.2, 145.2, 145.1, 141.3, 135.1, 131.6, 130.2, 129.4, 129.3, 129.1, 126.8, 126.5, 121.8, 107.6, 101.3, 96.4, 95.8, 70.9, 70.3, 70.1, 69.4, 69.3, 69.2, 67.5, 59.0, 57.6, 57.0, 56.2, 48.2, 43.6, 40.5, 40.4, 40.3, 40.3, 40.2, 40.1, 40.0, 39.8, 39.7, 39.5, 38.2, 36.2, 27.6, 27.5, 26.7, 26.5, 22.9, 16.5, 10.9.

**LC-MS** $C_{62}H_{83}N_9O_{14}S_2$ 1241.55; found $[M+H]^+$ 1246.62.

**dTRIM24-2 $^{1}H$ NMR** (500 MHz, DMSO-$d_6$)

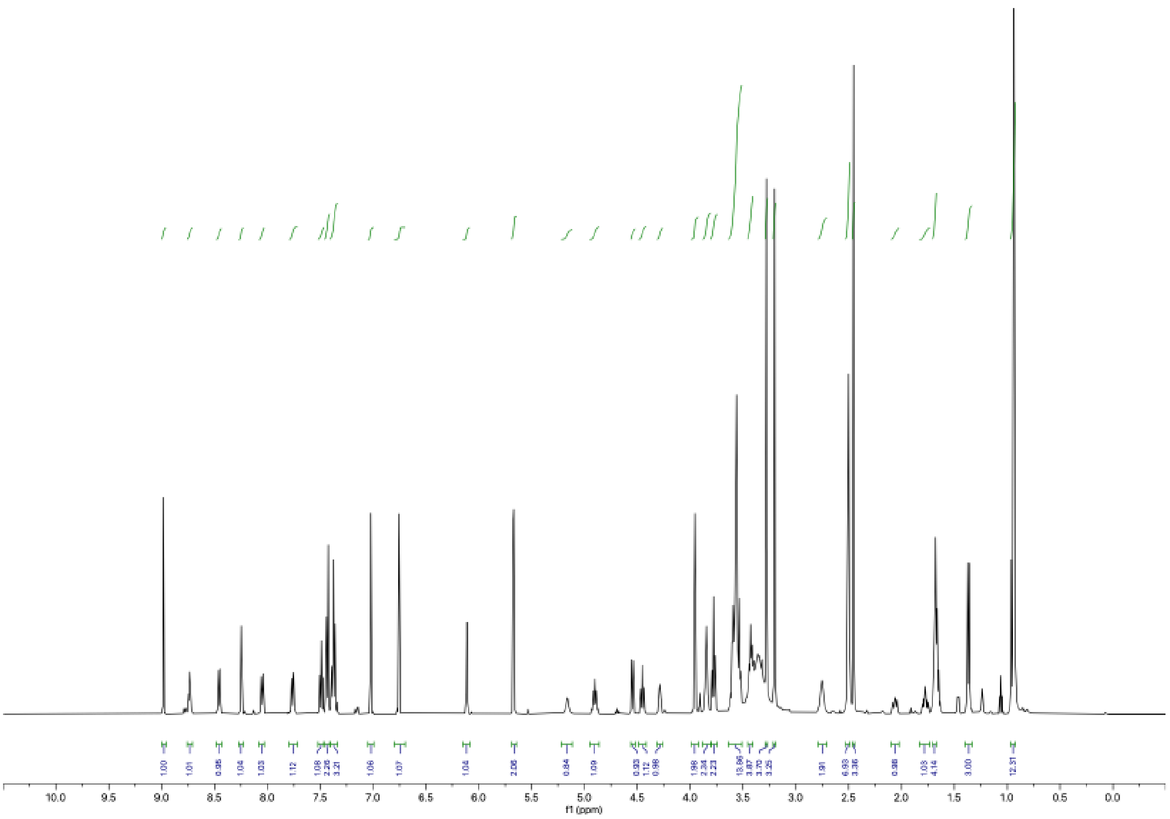


**dTRIM24-2 $^{13}C$ NMR** (126 MHz, DMSO-$d_6$)

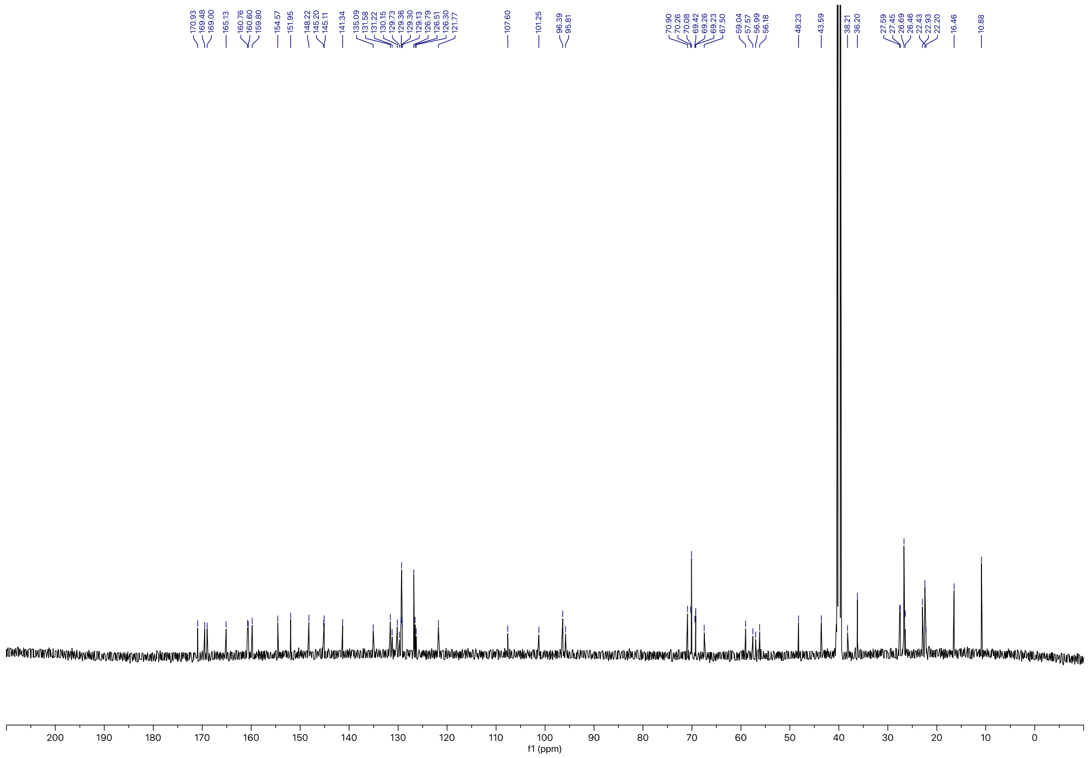
