## Supplemental Table for "TRIM24 Degradation Counteracts Adaptation to Androgen Receptor Inhibition in Prostate Cancer"

**Table S7E Summary of Transcription Factor (TF) Binding Motif Analysis on H3K27ac ChIP-Seq Data f**

| Motif Name | Consensus | P-value |
|---|---|---|
| RORg(NR)/Liver-Rorc-ChIP-Seq(GSE101115)/Homer | WAABTAGGT | 1.00E-99 |
| YY1(Zf)/Promoter/Homer | CAAGATGGC | 1.00E-99 |
| p53(p53)/mES-cMyc-ChIP-Seq(GSE11431)/Homer | ACATGCCCG | 1.00E-57 |
| Tbox:Smad(T-box,MAD)/ESCd5-Smad2_3-ChIP-Seq(GSE29422)/Homer | AGGTGHCAG | 1.00E-46 |
| Elf4(ETS)/BMDM-Elf4-ChIP-Seq(GSE88699)/Homer | ACTTCCKGK | 1.00E-35 |
| Fli1(ETS)/CD8-FLI-ChIP-Seq(GSE20898)/Homer | NRYTTCCGG | 1.00E-35 |
| VDR(NR),DR3/GM10855-VDR+vitD-ChIP-Seq(GSE22484)/Homer | ARAGGTCAN | 1.00E-34 |
| EHF(ETS)/LoVo-EHF-ChIP-Seq(GSE49402)/Homer | AVCAGGAAG | 1.00E-34 |
| Fox:Ebox(Forkhead,bHLH)/Panc1-Foxa2-ChIP-Seq(GSE47459)/Homer | NNNVCTGW( | 1.00E-32 |
| NFAT(RHD)/Jurkat-NFATC1-ChIP-Seq(Jolma_et_al.)/Homer | ATTTTCCATT | 1.00E-32 |
| Foxa3(Forkhead)/Liver-Foxa3-ChIP-Seq(GSE77670)/Homer | BSNTGTTTA( | 1.00E-31 |
| SPDEF(ETS)/VCaP-SPDEF-ChIP-Seq(SRA014231)/Homer | ASWTCCTGE | 1.00E-30 |
| NFkB-p65(RHD)/GM12787-p65-ChIP-Seq(GSE19485)/Homer | WGGGGATT | 1.00E-30 |
| Zfp281(Zf)/ES-Zfp281-ChIP-Seq(GSE81042)/Homer | CCCCTCCCC | 1.00E-30 |
| ETV1(ETS)/GIST48-ETV1-ChIP-Seq(GSE22441)/Homer | AACCGGAAG | 1.00E-30 |
| ELF5(ETS)/T47D-ELF5-ChIP-Seq(GSE30407)/Homer | ACVAGGAAG | 1.00E-30 |
| Elk1(ETS)/Hela-Elk1-ChIP-Seq(GSE31477)/Homer | HACTTCCGG | 1.00E-29 |
| ZFX(Zf)/mES-Zfx-ChIP-Seq(GSE11431)/Homer | AGGCCTRG | 1.00E-29 |
| ETV4(ETS)/HepG2-ETV4-ChIP-Seq(ENCODE)/Homer | ACCGGAAGT | 1.00E-29 |
| BMYB(HTH)/Hela-BMYB-ChIP-Seq(GSE27030)/Homer | NHAACBGYY | 1.00E-28 |
| Elk4(ETS)/Hela-Elk4-ChIP-Seq(GSE31477)/Homer | NRYTTCCGG | 1.00E-27 |
| AMYB(HTH)/Testes-AMYB-ChIP-Seq(GSE44588)/Homer | TGGCAGTTG | 1.00E-27 |
| Sox3(HMG)/NPC-Sox3-ChIP-Seq(GSE33059)/Homer | CCWTTGTY | 1.00E-26 |
| ERG(ETS)/VCaP-ERG-ChIP-Seq(GSE14097)/Homer | ACAGGAAGT | 1.00E-26 |
| GABPA(ETS)/Jurkat-GABPa-ChIP-Seq(GSE17954)/Homer | RACCGGAAG | 1.00E-25 |
| ELF1(ETS)/Jurkat-ELF1-ChIP-Seq(SRA014231)/Homer | AVCCGGAAG | 1.00E-24 |
| Foxa2(Forkhead)/Liver-Foxa2-ChIP-Seq(GSE25694)/Homer | CYTGTTTAC\ | 1.00E-23 |
| FOXP1(Forkhead)/H9-FOXP1-ChIP-Seq(GSE31006)/Homer | NYYTGTTTA( | 1.00E-23 |
| Sox10(HMG)/SciaticNerve-Sox3-ChIP-Seq(GSE35132)/Homer | CCWTTGTYY | 1.00E-23 |
| EBF2(EBF)/BrownAdipose-EBF2-ChIP-Seq(GSE97114)/Homer | NABTCCCWD | 1.00E-22 |
| GRHL2(CP2)/HBE-GRHL2-ChIP-Seq(GSE46194)/Homer | AAACYKGTT\ | 1.00E-22 |
| GFY-Staf(?,Zf)/Promoter/Homer | RACTACAAT | 1.00E-21 |
| Sox4(HMG)/proB-Sox4-ChIP-Seq(GSE50066)/Homer | YCTTTGTTC( | 1.00E-20 |
| MafB(bZIP)/BMM-Mafb-ChIP-Seq(GSE75722)/Homer | WNTGCTGAS | 1.00E-20 |
| ELF3(ETS)/PDAC-ELF3-ChIP-Seq(GSE64557)/Homer | ANCAGGAAG | 1.00E-20 |
| Smad2(MAD)/ES-SMAD2-ChIP-Seq(GSE29422)/Homer | CTGTCTGG | 1.00E-16 |
| ETS1(ETS)/Jurkat-ETS1-ChIP-Seq(GSE17954)/Homer | ACAGGAAGT | 1.00E-16 |
| AR-halfsite(NR)/LNCaP-AR-ChIP-Seq(GSE27824)/Homer | CCAGGAACA | 1.00E-16 |
| CEBP:AP1(bZIP)/ThioMac-CEBPb-ChIP-Seq(GSE21512)/Homer | DRTGTTGCA | 1.00E-16 |
| Etv2(ETS)/ES-ER71-ChIP-Seq(GSE59402)/Homer | NNAYTTCCT( | 1.00E-15 |
| Foxo3(Forkhead)/U2OS-Foxo3-ChIP-Seq(E-MTAB-2701)/Homer | DGTAAACA | 1.00E-15 |
| FOXK2(Forkhead)/U2OS-FOXK2-ChIP-Seq(E-MTAB-2204)/Homer | SCHTGTTTA( | 1.00E-15 |
| EWS:ERG-fusion(ETS)/CADO_ES1-EWS:ERG-ChIP-Seq(SRA014231)/Homer | ATTTCCTGTI | 1.00E-14 |
| FOXM1(Forkhead)/MCF7-FOXM1-ChIP-Seq(GSE72977)/Homer | TRTTTACTTV | 1.00E-14 |
| FoxD3(forkhead)/ZebrafishEmbryo-Foxd3.biotin-ChIP-seq(GSE106676)/Homer | TGTTTAYTTA | 1.00E-13 |
| Rbpj1(?)/Panc1-Rbpj1-ChIP-Seq(GSE47459)/Homer | HTTTCCCAS( | 1.00E-13 |
| FOXA1(Forkhead)/LNCAP-FOXA1-ChIP-Seq(GSE27824)/Homer | WAAGTAAAC | 1.00E-13 |
| Isl1(Homeobox)/Neuron-Isl1-ChIP-Seq(GSE31456)/Homer | CTAATKGV | 1.00E-13 |
| Sox2(HMG)/mES-Sox2-ChIP-Seq(GSE11431)/Homer | BCCATTGTT( | 1.00E-13 |
| FOXA1(Forkhead)/MCF7-FOXA1-ChIP-Seq(GSE26831)/Homer | WAAGTAAAC | 1.00E-12 |
| Zac1(Zf)/Neuro2A-Plagl1-ChIP-Seq(GSE75942)/Homer | HAWGRGGC( | 1.00E-12 |
| Smad4(MAD)/ESC-SMAD4-ChIP-Seq(GSE29422)/Homer | VBSYGTCTG | 1.00E-11 |

| | | |
|---|---|---|
| ZNF711(Zf)/SHSY5Y-ZNF711-ChIP-Seq(GSE20673)/Homer | AGGCCTAG | 1.00E-11 |
| ZNF467(Zf)/HEK293-ZNF467.GFP-ChIP-Seq(GSE58341)/Homer | TGGGGAAG | 1.00E-11 |
| PR(NR)/T47D-PR-ChIP-Seq(GSE31130)/Homer | VAGRACAKN | 1.00E-11 |
| Sox9(HMG)/Limb-SOX9-ChIP-Seq(GSE73225)/Homer | AGGVNCCTT | 1.00E-11 |
| Sox21(HMG)/ESC-SOX21-ChIP-Seq(GSE110505)/Homer | BCCWTTGTB | 1.00E-11 |
| PU.1-IRF(ETS:IRF)/Bcell-PU.1-ChIP-Seq(GSE21512)/Homer | MGGAAGTG | 1.00E-10 |
| FOXK1(Forkhead)/HEK293-FOXK1-ChIP-Seq(GSE51673)/Homer | NVWTGTTTA | 1.00E-10 |
| SCL(bHLH)/HPC7-Scl-ChIP-Seq(GSE13511)/Homer | AVCAGCTG | 1.00E-10 |
| ZNF341(Zf)/EBV-ZNF341-ChIP-Seq(GSE113194)/Homer | GGAACAGC | 1.00E-10 |
| Stat3+il21(Stat)/CD4-Stat3-ChIP-Seq(GSE19198)/Homer | SVYTTCCNG | 1.00E-09 |
| PRDM15(Zf)/ESC-Prdm15-ChIP-Seq(GSE73694)/Homer | YCCDNTCCA | 1.00E-09 |
| ETS(ETS)/Promoter/Homer | AACCGGAAG | 1.00E-09 |
| Tcfcp2l1(CP2)/mES-Tcfcp2l1-ChIP-Seq(GSE11431)/Homer | NRAACCRGT | 1.00E-09 |
| Stat3(Stat)/mES-Stat3-ChIP-Seq(GSE11431)/Homer | CTTCCGGG | 1.00E-09 |
| ZNF692(Zf)/HEK293-ZNF692.GFP-ChIP-Seq(GSE58341)/Homer | GTGGGCCC | 1.00E-09 |
| STAT4(Stat)/CD4-Stat4-ChIP-Seq(GSE22104)/Homer | NYTTCCWGC | 1.00E-08 |
| NF1-halfsite(CTF)/LNCaP-NF1-ChIP-Seq(Unpublished)/Homer | YTGCCAAG | 1.00E-08 |
| Foxo1(Forkhead)/RAW-Foxo1-ChIP-Seq(Fan_et_al.)/Homer | CTGTTTAC | 1.00E-08 |
| IRF2(IRF)/Erythroblas-IRF2-ChIP-Seq(GSE36985)/Homer | GAAASYGAA | 1.00E-08 |
| PU.1(ETS)/ThioMac-PU.1-ChIP-Seq(GSE21512)/Homer | AGAGGAAGT | 1.00E-07 |
| Foxf1(Forkhead)/Lung-Foxf1-ChIP-Seq(GSE77951)/Homer | WWATRTAA | 1.00E-07 |
| EWS:FLI1-fusion(ETS)/SK_N_MC-EWS:FLI1-ChIP-Seq(SRA014231)/Homer | VACAGGAAA | 1.00E-07 |
| AP-2gamma(AP2)/MCF7-TFAP2C-ChIP-Seq(GSE21234)/Homer | SCCTSAGGS | 1.00E-07 |
| FoxL2(Forkhead)/Ovary-FoxL2-ChIP-Seq(GSE60858)/Homer | WWTRTAAA | 1.00E-07 |
| Bcl6(Zf)/Liver-Bcl6-ChIP-Seq(GSE31578)/Homer | NNNCTTTCC | 1.00E-07 |
| Sox17(HMG)/Endoderm-Sox17-ChIP-Seq(GSE61475)/Homer | CCATTGTTY | 1.00E-06 |
| Pitx1(Homeobox)/Chicken-Pitx1-ChIP-Seq(GSE38910)/Homer | TAATCCCN | 1.00E-06 |
| E2F4(E2F)/K562-E2F4-ChIP-Seq(GSE31477)/Homer | GGCGGGAA | 1.00E-06 |
| AP-1(bZIP)/ThioMac-PU.1-ChIP-Seq(GSE21512)/Homer | VTGACTCAT | 1.00E-06 |
| MYB(HTH)/ERMYB-Myb-ChIPSeq(GSE22095)/Homer | GGCVGTTR | 1.00E-06 |
| AP-2alpha(AP2)/Hela-AP2alpha-ChIP-Seq(GSE31477)/Homer | ATGCCCTGA | 1.00E-06 |
| Atf3(bZIP)/GBM-ATF3-ChIP-Seq(GSE33912)/Homer | DATGASTCA | 1.00E-06 |
| IRF3(IRF)/BMDM-Irf3-ChIP-Seq(GSE67343)/Homer | AGTTTCAKT | 1.00E-06 |
| EBF1(EBF)/Near-E2A-ChIP-Seq(GSE21512)/Homer | GTCCCCWG | 1.00E-05 |
| WT1(Zf)/Kidney-WT1-ChIP-Seq(GSE90016)/Homer | MCTCCCMC | 1.00E-05 |
| Sox6(HMG)/Myotubes-Sox6-ChIP-Seq(GSE32627)/Homer | CCATTGTTN | 1.00E-05 |
| RORgt(NR)/EL4-RORgt.Flag-ChIP-Seq(GSE56019)/Homer | AAYTAGGTC | 1.00E-05 |
| RORgt(NR)/EL4-RORgt.Flag-ChIP-Seq(GSE56019)/Homer | AAYTAGGTC | 1.00E-05 |
| TEAD1(TEAD)/HepG2-TEAD1-ChIP-Seq(Encode)/Homer | CYRCATTCC | 1.00E-05 |
| Smad3(MAD)/NPC-Smad3-ChIP-Seq(GSE36673)/Homer | TWGTCTGV | 1.00E-05 |
| BATF(bZIP)/Th17-BATF-ChIP-Seq(GSE39756)/Homer | DATGASTCA | 1.00E-05 |
| Oct4:Sox17(POU,Homeobox,HMG)/F9-Sox17-ChIP-Seq(GSE44553)/Homer | CCATTGTAT | 1.00E-05 |
| Bach2(bZIP)/OCILy7-Bach2-ChIP-Seq(GSE44420)/Homer | TGCTGAGTC | 1.00E-05 |
| JunB(bZIP)/DendriticCells-Junb-ChIP-Seq(GSE36099)/Homer | RATGASTCA | 1.00E-05 |
| Fra1(bZIP)/BT549-Fra1-ChIP-Seq(GSE46166)/Homer | NNATGASTC | 1.00E-04 |
| NF1(CTF)/LNCAP-NF1-ChIP-Seq(Unpublished)/Homer | CYTGGCABN | 1.00E-04 |
| STAT1(Stat)/HelaS3-STAT1-ChIP-Seq(GSE12782)/Homer | NATTTCCNG | 1.00E-04 |
| Zic2(Zf)/ESC-Zic2-ChIP-Seq(SRP197560)/Homer | CHCAGCRG | 1.00E-04 |
| EBF(EBF)/proBcell-EBF-ChIP-Seq(GSE21978)/Homer | DGTCCCYRG | 1.00E-04 |
| p63(p53)/Keratinocyte-p63-ChIP-Seq(GSE17611)/Homer | NNDRCATGY | 1.00E-04 |
| Maz(Zf)/HepG2-Maz-ChIP-Seq(GSE31477)/Homer | GGGGGGGG | 1.00E-04 |
| SpiB(ETS)/OCILY3-SPIB-ChIP-Seq(GSE56857)/Homer | AAAGRGGA | 1.00E-04 |
| NFkB-p65-Rel(RHD)/ThioMac-LPS-Expression(GSE23622)/Homer | GGAAATTCC | 1.00E-04 |
| Fos(bZIP)/TSC-Fos-ChIP-Seq(GSE110950)/Homer | NDATGASTC | 1.00E-04 |

| | | |
|---|---|---|
| Fosl2(bZIP)/3T3L1-Fosl2-ChIP-Seq(GSE56872)/Homer | NATGASTCA | 1.00E-04 |
| IRF8(IRF)/BMDM-IRF8-ChIP-Seq(GSE77884)/Homer | GRAASTGAA | 1.00E-04 |
| Arnt:Ahr(bHLH)/MCF7-Arnt-ChIP-Seq(Lo_et_al.)/Homer | TBGCACGCA | 1.00E-04 |
| Fra2(bZIP)/Striatum-Fra2-ChIP-Seq(GSE43429)/Homer | GGATGACTC | 1.00E-04 |
| HLF(bZIP)/HSC-HLF.Flag-ChIP-Seq(GSE69817)/Homer | RTTATGYAAI | 1.00E-04 |
| NF1:FOXA1(CTF,Forkhead)/LNCAP-FOXA1-ChIP-Seq(GSE27824)/Homer | WNTGTTTRY | 1.00E-04 |
| TEAD3(TEA)/HepG2-TEAD3-ChIP-Seq(Encode)/Homer | TRCATTCCA | 1.00E-04 |
| Nr5a2(NR)/mES-Nr5a2-ChIP-Seq(GSE19019)/Homer | BTCAAGGTC | 1.00E-04 |
| Sox7(HMG)/ESC-Sox7-ChIP-Seq(GSE133899)/Homer | VVRRAACAA | 1.00E-03 |
| TEAD4(TEA)/Tropoblast-Tead4-ChIP-Seq(GSE37350)/Homer | CCWGGAAT | 1.00E-03 |
| ZNF7(Zf)/HepG2-ZNF7.Flag-ChIP-Seq(Encode)/Homer | CTGCCWVC | 1.00E-03 |
| Ets1-distal(ETS)/CD4+-PolII-ChIP-Seq(Barski_et_al.)/Homer | MACAGGAA | 1.00E-03 |
| CArG(MADS)/PUER-Srf-ChIP-Seq(Sullivan_et_al.)/Homer | CCATATATG | 1.00E-03 |
| TATA-Box(TBP)/Promoter/Homer | CCTTTTAWA | 1.00E-03 |
| STAT5(Stat)/mCD4+-Stat5-ChIP-Seq(GSE12346)/Homer | RTTTCTNAG | 1.00E-03 |
| Rfx6(HTH)/Min6b1-Rfx6.HA-ChIP-Seq(GSE62844)/Homer | TGTTKCCTA | 1.00E-03 |
| ISRE(IRF)/ThioMac-LPS-Expression(GSE23622)/Homer | AGTTTCAST | 1.00E-03 |
| Sox15(HMG)/CPA-Sox15-ChIP-Seq(GSE62909)/Homer | RAACAATGG | 1.00E-03 |
| IRF1(IRF)/PBMC-IRF1-ChIP-Seq(GSE43036)/Homer | GAAAGTGAA | 1.00E-03 |
| LRF(Zf)/Erythroblasts-ZBTB7A-ChIP-Seq(GSE74977)/Homer | AAGACCCYY | 1.00E-03 |
| PU.1:IRF8(ETS:IRF)/pDC-Irf8-ChIP-Seq(GSE66899)/Homer | GGAAGTGAA | 1.00E-03 |
| Hand2(bHLH)/Mesoderm-Hand2-ChIP-Seq(GSE61475)/Homer | TGACANARR | 1.00E-03 |
| Foxh1(Forkhead)/hESC-FOXH1-ChIP-Seq(GSE29422)/Homer | NNTGTGGAT | 1.00E-03 |
| E2F6(E2F)/Hela-E2F6-ChIP-Seq(GSE31477)/Homer | GGCGGGAAI | 1.00E-03 |
| ZNF528(Zf)/HEK293-ZNF528.GFP-ChIP-Seq(GSE58341)/Homer | AGAAATGAC | 1.00E-02 |
| IRF4(IRF)/GM12878-IRF4-ChIP-Seq(GSE32465)/Homer | ACTGAAACC | 1.00E-02 |
| Zfp57(Zf)/H1-ZFP57.HA-ChIP-Seq(GSE115387)/Homer | NANTGCSGC | 1.00E-02 |
| Tbr1(T-box)/Cortex-Tbr1-ChIP-Seq(GSE71384)/Homer | AAGGTGTKA | 1.00E-02 |
| RORa(NR)/Liver-Rora-ChIP-Seq(GSE101115)/Homer | AAWCTAGGT | 1.00E-02 |
| MafA(bZIP)/Islet-MafA-ChIP-Seq(GSE30298)/Homer | TGCTGACTC | 1.00E-02 |
| Nr5a2(NR)/Pancreas-LRH1-ChIP-Seq(GSE34295)/Homer | BTCAAGGTC | 1.00E-02 |
| Zic(Zf)/Cerebellum-ZIC1.2-ChIP-Seq(GSE60731)/Homer | CCTGCTGAG | 1.00E-02 |
| STAT6(Stat)/CD4-Stat6-ChIP-Seq(GSE22104)/Homer | ABTTCYYRR | 1.00E-02 |
| E2F1(E2F)/Hela-E2F1-ChIP-Seq(GSE22478)/Homer | CWGGCGGG | 1.00E-02 |
| HRE(HSF)/Striatum-HSF1-ChIP-Seq(GSE38000)/Homer | TTCTAGAABI | 1.00E-02 |
| HRE(HSF)/HepG2-HSF1-ChIP-Seq(GSE31477)/Homer | BSTTCTRGA | 1.00E-02 |
| ARE(NR)/LNCAP-AR-ChIP-Seq(GSE27824)/Homer | RGRACASNS | 1.00E-02 |
| PRDM1(Zf)/Hela-PRDM1-ChIP-Seq(GSE31477)/Homer | ACTTTCACT | 1.00E-02 |
| Tlx?(NR)/NPC-H3K4me1-ChIP-Seq(GSE16256)/Homer | CTGGCAGSC | 1.00E-02 |
| STAT6(Stat)/Macrophage-Stat6-ChIP-Seq(GSE38377)/Homer | TTCCKNAGA | 1.00E-02 |
| TRPS1(Zf)/MCF7-TRPS1-ChIP-Seq(GSE107013)/Homer | AGATAAGAN | 1.00E-02 |
| Oct11(POU,Homeobox)/NCIH1048-POU2F3-ChIP-seq(GSE115123)/Homer | GATTTGCAT | 1.00E-02 |
| E2F7(E2F)/Hela-E2F7-ChIP-Seq(GSE32673)/Homer | VDTTTCCCG | 1.00E-02 |
| THRb(NR)/Liver-NR1A2-ChIP-Seq(GSE52613)/Homer | TRAGGTCA | 1.00E-02 |
| Hoxd11(Homeobox)/ChickenMSG-Hoxd11.Flag-ChIP-Seq(GSE86088)/Homer | VGCCATAAA | 1.00E-02 |
| RXR(NR),DR1/3T3L1-RXR-ChIP-Seq(GSE13511)/Homer | TAGGGCAAA | 1.00E-02 |
| Hoxa13(Homeobox)/ChickenMSG-Hoxa13.Flag-ChIP-Seq(GSE86088)/Homer | CYHATAAAAI | 1.00E-02 |
| SF1(NR)/H295R-Nr5a1-ChIP-Seq(GSE44220)/Homer | CAAGGHCAN | 1.00E-02 |
| Hoxd10(Homeobox)/ChickenMSG-Hoxd10.Flag-ChIP-Seq(GSE86088)/Homer | GGCMATGA | 1.00E-02 |
| Bach1(bZIP)/K562-Bach1-ChIP-Seq(GSE31477)/Homer | AWWNTGCT | 1.00E-02 |
| PGR(NR)/EndoStromal-PGR-ChIP-Seq(GSE69539)/Homer | AAGAACATW | 1.00E-01 |
| Hoxc9(Homeobox)/Ainv15-Hoxc9-ChIP-Seq(GSE21812)/Homer | GGCCATAAA | 1.00E-01 |
| Nkx6.1(Homeobox)/Islet-Nkx6.1-ChIP-Seq(GSE40975)/Homer | GKTAATGR | 1.00E-01 |
| IRF:BATF(IRF:bZIP)/pDC-Irf8-ChIP-Seq(GSE66899)/Homer | CTTTCANTA | 1.00E-01 |

| | | |
|---|---|---|
| ZNF189(Zf)/HEK293-ZNF189.GFP-ChIP-Seq(GSE58341)/Homer | TGGAACAGM | 1.00E-01 |
| DLX1(Homeobox)/BasalGanglia-Dlx1-ChIP-seq(GSE124936)/Homer | NSNNTAATTA | 1.00E-01 |
| Reverb(NR),DR2/RAW-Reverba.biotin-ChIP-Seq(GSE45914)/Homer | GTRGGTCAS | 1.00E-01 |
| ZNF322(Zf)/HEK293-ZNF322.GFP-ChIP-Seq(GSE58341)/Homer | GAGCCTGGT | 1.00E-01 |
| Oct6(POU,Homeobox)/NPC-Pou3f1-ChIP-Seq(GSE35496)/Homer | WATGCAAAT | 1.00E-01 |
| Egr1(Zf)/K562-Egr1-ChIP-Seq(GSE32465)/Homer | TGCGTGGGY | 1.00E-01 |
| MyoG(bHLH)/C2C12-MyoG-ChIP-Seq(GSE36024)/Homer | AACAGCTG | 1.00E-01 |
| Esrrb(NR)/mES-Esrrb-ChIP-Seq(GSE11431)/Homer | KTGACCTTG | 1.00E-01 |
| TEAD(TEA)/Fibroblast-PU.1-ChIP-Seq(Unpublished)/Homer | YCWGGAATC | 1.00E-01 |
| E2F3(E2F)/MEF-E2F3-ChIP-Seq(GSE71376)/Homer | BTKGGCGGC | 1.00E-01 |
| PRDM14(Zf)/H1-PRDM14-ChIP-Seq(GSE22767)/Homer | RGGTCTCTA | 1.00E-01 |
| MYNN(Zf)/HEK293-MYNN.eGFP-ChIP-Seq(Encode)/Homer | TTCAAAWTA | 1.00E-01 |
| Tcf3(HMG)/mES-Tcf3-ChIP-Seq(GSE11724)/Homer | ASWTCAAAG | 1.00E-01 |
| Jun-AP1(bZIP)/K562-cJun-ChIP-Seq(GSE31477)/Homer | GATGASTCA | 1.00E-01 |
| Tcf7(HMG)/GM12878-TCF7-ChIP-Seq(Encode)/Homer | CTTTGATGT | 1.00E-01 |
| Sp2(Zf)/HEK293-Sp2.eGFP-ChIP-Seq(Encode)/Homer | YGGCCCCGC | 1.00E-01 |
| ZNF264(Zf)/HEK293-ZNF264.GFP-ChIP-Seq(GSE58341)/Homer | RGGGCACTA | 1.00E-01 |
| Brn1(POU,Homeobox)/NPC-Brn1-ChIP-Seq(GSE35496)/Homer | TATGCWAAT | 1.00E-01 |
| RARa(NR)/K562-RARa-ChIP-Seq(Encode)/Homer | TTGAMCTTT | 1.00E-01 |
| ZNF652/HepG2-ZNF652.Flag-ChIP-Seq(Encode)/Homer | TTAACCCTT | 1.00E-01 |
| TCFL2(HMG)/K562-TCF7L2-ChIP-Seq(GSE29196)/Homer | ACWTCAAAG | 1.00E-01 |
| Unknown(Homeobox)/Limb-p300-ChIP-Seq/Homer | SSCMATWAA | 1.00E-01 |
| HNF4a(NR),DR1/HepG2-HNF4a-ChIP-Seq(GSE25021)/Homer | CARRGKBCA | 1.00E-01 |
| DLX2(Homeobox)/BasalGanglia-Dlx2-ChIP-seq(GSE124936)/Homer | NNNTAATTAS | 1.00E-01 |
| LEF1(HMG)/H1-LEF1-ChIP-Seq(GSE64758)/Homer | CCTTTGATST | 1.00E-01 |
| TEAD2(TEA)/Py2T-Tead2-ChIP-Seq(GSE55709)/Homer | CCWGGAATC | 1.00E-01 |
| GRE(NR),IR3/A549-GR-ChIP-Seq(GSE32465)/Homer | NRGVACABN | 1.00E-01 |
| Ap4(bHLH)/AML-Tfap4-ChIP-Seq(GSE45738)/Homer | NAHCAGCTG | 1.00E-01 |
| ZNF675(Zf)/HEK293-ZNF675.GFP-ChIP-Seq(GSE58341)/Homer | ARGAGGMCA | 1.00E-01 |
| PPARE(NR),DR1/3T3L1-Pparg-ChIP-Seq(GSE13511)/Homer | TGACCTTTG | 1.00E-01 |
| PPARa(NR),DR1/Liver-Ppara-ChIP-Seq(GSE47954)/Homer | VNAGGKCAA | 1.00E-01 |
| p53(p53)/Saos-p53-ChIP-Seq(GSE15780)/Homer | RRCATGYCY | 1.00E-01 |
| p53(p53)/Saos-p53-ChIP-Seq/Homer | RRCATGYCY | 1.00E-01 |
| ZKSCAN1(Zf)/HepG2-ZKSCAN1-ChIP-Seq(Encode)/Homer | GCACAYAGT | 1.00E-01 |
| PAX6(Paired,Homeobox)/Forebrain-Pax6-ChIP-Seq(GSE66961)/Homer | NGTGTTCAV | 1.00E-01 |
| Six2(Homeobox)/NephronProgenitor-Six2-ChIP-Seq(GSE39837)/Homer | GWAAYHTGA | 1.00E-01 |
| Oct4(POU,Homeobox)/mES-Oct4-ChIP-Seq(GSE11431)/Homer | ATTTGCATAV | 1.00E-01 |
| HINFP(Zf)/K562-HINFP.eGFP-ChIP-Seq(Encode)/Homer | TWVGGTCCC | 1.00E-01 |
| DLX5(Homeobox)/BasalGanglia-Dlx5-ChIP-seq(GSE124936)/Homer | SSTAATTA | 1.00E-01 |
| NFAT:AP1(RHD,bZIP)/Jurkat-NFATC1-ChIP-Seq(Jolma_et_al.)/Homer | SARTGGAAA | 1.00E-01 |
| Hoxa9(Homeobox)/ChickenMSG-Hoxa9.Flag-ChIP-Seq(GSE86088)/Homer | RGCAATNAA | 1.00E+00 |
| RUNX2(Runt)/PCa-RUNX2-ChIP-Seq(GSE33889)/Homer | NWAACCACA | 1.00E+00 |
| Brachyury(T-box)/Mesoendoderm-Brachyury-ChIP-exo(GSE54963)/Homer | ANTTMRCAS | 1.00E+00 |
| NFkB-p50,p52(RHD)/Monocyte-p50-ChIP-Chip(Schreiber_et_al.)/Homer | GGGGGAATC | 1.00E+00 |
| p73(p53)/Trachea-p73-ChIP-Seq(PRJNA310161)/Homer | NRRRCAWGT | 1.00E+00 |
| Tcf21(bHLH)/ArterySmoothMuscle-Tcf21-ChIP-Seq(GSE61369)/Homer | NAACAGCTG | 1.00E+00 |
| Pitx1:Ebox(Homeobox,bHLH)/Hindlimb-Pitx1-ChIP-Seq(GSE41591)/Homer | YTAATTRAW | 1.00E+00 |
| CRX(Homeobox)/Retina-Crx-ChIP-Seq(GSE20012)/Homer | GCTAATCC | 1.00E+00 |
| OCT:OCT(POU,Homeobox,IR1)/NPC-Brn2-ChIP-Seq(GSE35496)/Homer | ATGAATWAT | 1.00E+00 |
| HOXB13(Homeobox)/ProstateTumor-HOXB13-ChIP-Seq(GSE56288)/Homer | TTTTATKRGC | 1.00E+00 |
| RAR:RXR(NR),DR5/ES-RAR-ChIP-Seq(GSE56893)/Homer | RGGTCADNN | 1.00E+00 |
| ETS:RUNX(ETS,Runt)/Jurkat-RUNX1-ChIP-Seq(GSE17954)/Homer | RCAGGATGT | 1.00E+00 |
| Hoxa11(Homeobox)/ChickenMSG-Hoxa11.Flag-ChIP-Seq(GSE86088)/Homer | TTTTATGGCI | 1.00E+00 |
| ZBTB12(Zf)/HEK293-ZBTB12.GFP-ChIP-Seq(GSE58341)/Homer | NGNTCTAGA | 1.00E+00 |

| | | |
|---|---|---|
| RBPJ:Ebox(?,bHLH)/Panc1-Rbpj1-ChIP-Seq(GSE47459)/Homer | GGGRAARR( | 1.00E+00 |
| Nrf2(bZIP)/Lymphoblast-Nrf2-ChIP-Seq(GSE37589)/Homer | HTGCTGAGT | 1.00E+00 |
| PBX2(Homeobox)/K562-PBX2-ChIP-Seq(Encode)/Homer | RTGATTKATI | 1.00E+00 |
| NF-E2(bZIP)/K562-NFE2-ChIP-Seq(GSE31477)/Homer | GATGACTCA | 1.00E+00 |
| Hoxb4(Homeobox)/ES-Hoxb4-ChIP-Seq(GSE34014)/Homer | TGATTRATG | 1.00E+00 |
| ZNF165(Zf)/WHIM12-ZNF165-ChIP-Seq(GSE65937)/Homer | AAGGKGRC( | 1.00E+00 |
| Hoxa10(Homeobox)/ChickenMSG-Hoxa10.Flag-ChIP-Seq(GSE86088)/Homer | GGYAATGAA | 1.00E+00 |
| DMRT6(DM)/Testis-DMRT6-ChIP-Seq(GSE60440)/Homer | YDGHTACAW | 1.00E+00 |
| Lhx3(Homeobox)/Neuron-Lhx3-ChIP-Seq(GSE31456)/Homer | ADBTAATTAI | 1.00E+00 |
| DMRT1(DM)/Testis-DMRT1-ChIP-Seq(GSE64892)/Homer | TWGHWACA\ | 1.00E+00 |
| FOXA1:AR(Forkhead,NR)/LNCAP-AR-ChIP-Seq(GSE27824)/Homer | AGTAAACAA | 1.00E+00 |
| EBNA1(EBV-virus)/Raji-EBNA1-ChIP-Seq(GSE30709)/Homer | GGYAGCAYD | 1.00E+00 |
| OCT4-SOX2-TCF-NANOG(POU,Homeobox,HMG)/mES-Oct4-ChIP-Seq(GSE1 | ATTTGCATA/ | 1.00E+00 |
| Zic3(Zf)/mES-Zic3-ChIP-Seq(GSE37889)/Homer | GGCCYCCTG | 1.00E+00 |
| ZNF768(Zf)/Rajj-ZNF768-ChIP-Seq(GSE111879)/Homer | RHHCAGAGA | 1.00E+00 |
| ERRg(NR)/Kidney-ESRRG-ChIP-Seq(GSE104905)/Homer | GTGACCTTG | 1.00E+00 |
| ETS:E-box(ETS,bHLH)/HPC7-Scl-ChIP-Seq(GSE22178)/Homer | AGGAARCAG | 1.00E+00 |
| PRDM9(Zf)/Testis-DMC1-ChIP-Seq(GSE35498)/Homer | ADGGYAGYA | 1.00E+00 |
| Znf263(Zf)/K562-Znf263-ChIP-Seq(GSE31477)/Homer | CVGTSCTCC | 1.00E+00 |
| MafK(bZIP)/C2C12-MafK-ChIP-Seq(GSE36030)/Homer | GCTGASTCA | 1.00E+00 |
| Tbx20(T-box)/Heart-Tbx20-ChIP-Seq(GSE29636)/Homer | GGTGYTGAC | 1.00E+00 |
| RUNX1(Runt)/Jurkat-RUNX1-ChIP-Seq(GSE29180)/Homer | AAACCACAR | 1.00E+00 |
| Gata2(Zf)/K562-GATA2-ChIP-Seq(GSE18829)/Homer | BBCTTATCTS | 1.00E+00 |
| PAX3:FKHR-fusion(Paired,Homeobox)/Rh4-PAX3:FKHR-ChIP-Seq(GSE1906 | ACCRTGACT | 1.00E+00 |
| Dlx3(Homeobox)/Kerainocytes-Dlx3-ChIP-Seq(GSE89884)/Homer | NDGTAATTA | 1.00E+00 |
| Gata4(Zf)/Heart-Gata4-ChIP-Seq(GSE35151)/Homer | NBWGATAAG | 1.00E+00 |
| Nanog(Homeobox)/mES-Nanog-ChIP-Seq(GSE11724)/Homer | RGCCATTAA | 1.00E+00 |
| COUP-TFII(NR)/Artia-Nr2f2-ChIP-Seq(GSE46497)/Homer | AGRGGTCA | 1.00E+00 |
| Gata1(Zf)/K562-GATA1-ChIP-Seq(GSE18829)/Homer | SAGATAAGR | 1.00E+00 |
| Hoxd13(Homeobox)/ChickenMSG-Hoxd13.Flag-ChIP-Seq(GSE86088)/Homer | NCYAATAAA | 1.00E+00 |
| CTCF-SatelliteElement(Zf?)/CD4+-CTCF-ChIP-Seq(Barski_et_al.)/Homer | TGCAGTTCC | 1.00E+00 |
| NFE2L2(bZIP)/HepG2-NFE2L2-ChIP-Seq(Encode)/Homer | AWWWTGCT | 1.00E+00 |
| ZNF16(Zf)/HEK293-ZNF16.GFP-ChIP-Seq(GSE58341)/Homer | MACCTTCYA | 1.00E+00 |
| Egr2(Zf)/Thymocytes-Egr2-ChIP-Seq(GSE34254)/Homer | NGCGTGGG | 1.00E+00 |
| T1ISRE(IRF)/ThioMac-Ifnb-Expression/Homer | ACTTTCGTT | 1.00E+00 |
| Six4(Homeobox)/MCF7-SIX4-ChIP-Seq(Encode)/Homer | TGWAAYCTG | 1.00E+00 |
| Otx2(Homeobox)/EpiLC-Otx2-ChIP-Seq(GSE56098)/Homer | NYTAATCCYI | 1.00E+00 |
| Pit1(Homeobox)/GCrat-Pit1-ChIP-Seq(GSE58009)/Homer | ATGMATATD | 1.00E+00 |
| Nur77(NR)/K562-NR4A1-ChIP-Seq(GSE31363)/Homer | TGACCTTTN | 1.00E+00 |
| CEBP(bZIP)/ThioMac-CEBPb-ChIP-Seq(GSE21512)/Homer | ATTGCGCAA | 1.00E+00 |
| Oct2(POU,Homeobox)/Bcell-Oct2-ChIP-Seq(GSE21512)/Homer | ATATGCAAA | 1.00E+00 |
| HIF2a(bHLH)/785_O-HIF2a-ChIP-Seq(GSE34871)/Homer | GCACGTACC | 1.00E+00 |
| KLF5(Zf)/LoVo-KLF5-ChIP-Seq(GSE49402)/Homer | DGGGYGKG | 1.00E+00 |
| NFIL3(bZIP)/HepG2-NFIL3-ChIP-Seq(Encode)/Homer | VTTACGTAA | 1.00E+00 |
| Gata6(Zf)/HUG1N-GATA6-ChIP-Seq(GSE51936)/Homer | YCTTATCTBI | 1.00E+00 |
| Myf5(bHLH)/GM-Myf5-ChIP-Seq(GSE24852)/Homer | BAACAGCTG | 1.00E+00 |
| MyoD(bHLH)/Myotube-MyoD-ChIP-Seq(GSE21614)/Homer | RRCAGCTGY | 1.00E+00 |
| KLF14(Zf)/HEK293-KLF14.GFP-ChIP-Seq(GSE58341)/Homer | RGKGGGCGI | 1.00E+00 |
| RUNX(Runt)/HPC7-Runx1-ChIP-Seq(GSE22178)/Homer | SAAACCACA | 1.00E+00 |
| ZFP3(Zf)/HEK293-ZFP3.GFP-ChIP-Seq(GSE58341)/Homer | GGGTTTTGA | 1.00E+00 |
| Mouse_Recombination_Hotspot(Zf)/Testis-DMC1-ChIP-Seq(GSE24438)/Home | ACTYKNATT | 1.00E+00 |
| GRE(NR),IR3/RAW264.7-GRE-ChIP-Seq(Unpublished)/Homer | VAGRACAKW | 1.00E+00 |
| GFX(?)/Promoter/Homer | ATTCTCGCG | 1.00E+00 |
| CHR(?)/Hela-CellCycle-Expression/Homer | SRGTTTCAA | 1.00E+00 |

| | | |
|---|---|---|
| Ascl1(bHLH)/NeuralTubes-Ascl1-ChIP-Seq(GSE55840)/Homer | NNVVCAGCT | 1.00E+00 |
| Eomes(T-box)/H9-Eomes-ChIP-Seq(GSE26097)/Homer | ATTAACACC | 1.00E+00 |
| Meis1(Homeobox)/MastCells-Meis1-ChIP-Seq(GSE48085)/Homer | VGCTGWCA | 1.00E+00 |
| Lhx2(Homeobox)/HFSC-Lhx2-ChIP-Seq(GSE48068)/Homer | TAATTAGN | 1.00E+00 |
| Lhx1(Homeobox)/EmbryoCarcinoma-Lhx1-ChIP-Seq(GSE70957)/Homer | NNYTAATTA | 1.00E+00 |
| Mef2d(MADS)/Retina-Mef2d-ChIP-Seq(GSE61391)/Homer | GCTATTTTT | 1.00E+00 |
| PAX5(Paired,Homeobox),condensed/GM12878-PAX5-ChIP-Seq(GSE32465)/I | GTCACGCTC | 1.00E+00 |
| ZNF382(Zf)/HEK293-ZNF382.GFP-ChIP-Seq(GSE58341)/Homer | GNCTGTAST | 1.00E+00 |
| bZIP:IRF(bZIP,IRF)/Th17-BatF-ChIP-Seq(GSE39756)/Homer | NAGTTTCAB | 1.00E+00 |
| Hnf1(Homeobox)/Liver-Foxa2-Chip-Seq(GSE25694)/Homer | GGTTAAWC | 1.00E+00 |
| ZNF669(Zf)/HEK293-ZNF669.GFP-ChIP-Seq(GSE58341)/Homer | GARTGGTCA | 1.00E+00 |
| Olig2(bHLH)/Neuron-Olig2-ChIP-Seq(GSE30882)/Homer | RCCATMTGT | 1.00E+00 |
| E2F(E2F)/Hela-CellCycle-Expression/Homer | TTSGCGCGA | 1.00E+00 |
| ERE(NR),IR3/MCF7-ERa-ChIP-Seq(Unpublished)/Homer | VAGGTCACN | 1.00E+00 |
| Cdx2(Homeobox)/mES-Cdx2-ChIP-Seq(GSE14586)/Homer | GYMATAAAA | 1.00E+00 |
| HNF1b(Homeobox)/PDAC-HNF1B-ChIP-Seq(GSE64557)/Homer | GTTAATNAT | 1.00E+00 |
| RARg(NR)/ES-RARg-ChIP-Seq(GSE30538)/Homer | AGGTCAAGG | 1.00E+00 |
| Six1(Homeobox)/Myoblast-Six1-ChIP-Chip(GSE20150)/Homer | GKVTCADRT | 1.00E+00 |
| CEBP:CEBP(bZIP)/MEF-Chop-ChIP-Seq(GSE35681)/Homer | NTNATGCAA | 1.00E+00 |
| CDX4(Homeobox)/ZebrafishEmbryos-Cdx4.Myc-ChIP-Seq(GSE48254)/Homer | NGYCATAAA | 1.00E+00 |
| Tcf12(bHLH)/GM12878-Tcf12-ChIP-Seq(GSE32465)/Homer | VCAGCTGYT | 1.00E+00 |
| OCT:OCT-short(POU,Homeobox)/NPC-OCT6-ChIP-Seq(GSE43916)/Homer | ATGCATWAT | 1.00E+00 |
| GATA3(Zf),DR8/iTreg-Gata3-ChIP-Seq(GSE20898)/Homer | AGATSTNDN | 1.00E+00 |
| LHX9(Homeobox)/Hct116-LHX9.V5-ChIP-Seq(GSE116822)/Homer | NGCTAATTA | 1.00E+00 |
| ZNF416(Zf)/HEK293-ZNF416.GFP-ChIP-Seq(GSE58341)/Homer | WDNCTGGG | 1.00E+00 |
| ZNF317(Zf)/HEK293-ZNF317.GFP-ChIP-Seq(GSE58341)/Homer | GTCWGCTG | 1.00E+00 |
| RAR:RXR(NR),DR0/ES-RAR-ChIP-Seq(GSE56893)/Homer | AGGTCAAGG | 1.00E+00 |
| Pbx3(Homeobox)/GM12878-PBX3-ChIP-Seq(GSE32465)/Homer | SCTGTCAMT | 1.00E+00 |
| HIC1(Zf)/Treg-ZBTB29-ChIP-Seq(GSE99889)/Homer | TGCCAGCB | 1.00E+00 |
| Unknown-ESC-element(?)/mES-Nanog-ChIP-Seq(GSE11724)/Homer | CACAGCAGG | 1.00E+00 |
| Pdx1(Homeobox)/Islet-Pdx1-ChIP-Seq(SRA008281)/Homer | YCATYAATC | 1.00E+00 |
| MafF(bZIP)/HepG2-MafF-ChIP-Seq(GSE31477)/Homer | HWWGTCAG | 1.00E+00 |
| X-box(HTH)/NPC-H3K4me1-ChIP-Seq(GSE16256)/Homer | GGTTGCCAT | 1.00E+00 |
| Tbx5(T-box)/HL1-Tbx5.biotin-ChIP-Seq(GSE21529)/Homer | AGGTGTCA | 1.00E+00 |
| Rfx1(HTH)/NPC-H3K4me1-ChIP-Seq(GSE16256)/Homer | KGTTGCCAT | 1.00E+00 |
| Rfx5(HTH)/GM12878-Rfx5-ChIP-Seq(GSE31477)/Homer | SCCTAGCAA | 1.00E+00 |
| HIF-1b(HLH)/T47D-HIF1b-ChIP-Seq(GSE59937)/Homer | RTACGTGC | 1.00E+00 |
| Atoh1(bHLH)/Cerebellum-Atoh1-ChIP-Seq(GSE22111)/Homer | VNRVCAGCT | 1.00E+00 |
| OCT:OCT(POU,Homeobox)/NPC-Brn1-ChIP-Seq(GSE35496)/Homer | ATGAATATT | 1.00E+00 |
| PSE(SNAPc)/K562-mStart-Seq/Homer | WAVTCACC | 1.00E+00 |
| Brn2(POU,Homeobox)/NPC-Brn2-ChIP-Seq(GSE35496)/Homer | ATGAATATT | 1.00E+00 |
| EAR2(NR)/K562-NR2F6-ChIP-Seq(Encode)/Homer | NRBCARRGG | 1.00E+00 |
| Sp5(Zf)/mES-Sp5.Flag-ChIP-Seq(GSE72989)/Homer | RGKGGGCG | 1.00E+00 |
| En1(Homeobox)/SUM149-EN1-ChIP-Seq(GSE120957)/Homer | NDCTAATTA | 1.00E+00 |
| Pknox1(Homeobox)/ES-Prep1-ChIP-Seq(GSE63282)/Homer | SCTGTCAVT | 1.00E+00 |
| ZBTB33(Zf)/GM12878-ZBTB33-ChIP-Seq(GSE32465)/Homer | GGVTCTCGC | 1.00E+00 |
| RUNX-AML(Runt)/CD4+-PolII-ChIP-Seq(Barski_et_al.)/Homer | GCTGTGGTT | 1.00E+00 |
| COUP-TFII(NR)/K562-NR2F1-ChIP-Seq(Encode)/Homer | GKBCARAGG | 1.00E+00 |
| GATA3(Zf)/iTreg-Gata3-ChIP-Seq(GSE20898)/Homer | AGATAASR | 1.00E+00 |
| Atf4(bZIP)/MEF-Atf4-ChIP-Seq(GSE35681)/Homer | MTGATGCAA | 1.00E+00 |
| Zfp809(Zf)/ES-Zfp809-ChIP-Seq(GSE70799)/Homer | GGGGCTYG | 1.00E+00 |
| Tbx6(T-box)/ESC-Tbx6-ChIP-Seq(GSE93524)/Homer | DAGGTGTBA | 1.00E+00 |
| GATA3(Zf),DR4/iTreg-Gata3-ChIP-Seq(GSE20898)/Homer | AGATGKDGA | 1.00E+00 |
| ZNF519(Zf)/HEK293-ZNF519.GFP-ChIP-Seq(GSE58341)/Homer | GAGSCCGA | 1.00E+00 |

| | | |
|---|---|---|
| GLI3(Zf)/Limb-GLI3-ChIP-Chip(GSE11077)/Homer | CGTGGGTG( | 1.00E+00 |
| Tgif2(Homeobox)/mES-Tgif2-ChIP-Seq(GSE55404)/Homer | TGTCANYT | 1.00E+00 |
| ZEB1(Zf)/PDAC-ZEB1-ChIP-Seq(GSE64557)/Homer | VCAGGTRDF | 1.00E+00 |
| GATA(Zf),IR4/iTreg-Gata3-ChIP-Seq(GSE20898)/Homer | NAGATWNBN | 1.00E+00 |
| Pit1+1bp(Homeobox)/GCrat-Pit1-ChIP-Seq(GSE58009)/Homer | ATGCATAAT | 1.00E+00 |
| ZNF415(Zf)/HEK293-ZNF415.GFP-ChIP-Seq(GSE58341)/Homer | GRTGMTRG/ | 1.00E+00 |
| E2A(bHLH)/proBcell-E2A-ChIP-Seq(GSE21978)/Homer | DNRCAGCTG | 1.00E+00 |
| RFX(HTH)/K562-RFX3-ChIP-Seq(SRA012198)/Homer | CGGTTGCCA | 1.00E+00 |
| ZBTB18(Zf)/HEK293-ZBTB18.GFP-ChIP-Seq(GSE58341)/Homer | AACATCTGG | 1.00E+00 |
| Erra(NR)/HepG2-Erra-ChIP-Seq(GSE31477)/Homer | CAAAGGTCA | 1.00E+00 |
| Pax7(Paired,Homeobox),long/Myoblast-Pax7-ChIP-Seq(GSE25064)/Homer | TAATCHGAT | 1.00E+00 |
| Pax7(Paired,Homeobox)/Myoblast-Pax7-ChIP-Seq(GSE25064)/Homer | TAATCAATT/ | 1.00E+00 |
| Rfx2(HTH)/LoVo-RFX2-ChIP-Seq(GSE49402)/Homer | GTTGCCATG | 1.00E+00 |
| HNF6(Homeobox)/Liver-Hnf6-ChIP-Seq(ERP000394)/Homer | NTATYGATCI | 1.00E+00 |
| BHLHA15(bHLH)/NIH3T3-BHLHB8.HA-ChIP-Seq(GSE119782)/Homer | NAMCAGCTG | 1.00E+00 |
| NRF(NRF)/Promoter/Homer | STGCGCATG | 1.00E+00 |
| HEB(bHLH)/mES-Heb-ChIP-Seq(GSE53233)/Homer | VCAGCTGBN | 1.00E+00 |
| TR4(NR),DR1/Hela-TR4-ChIP-Seq(GSE24685)/Homer | GAGGTCAAA | 1.00E+00 |
| FXR(NR),IR1/Liver-FXR-ChIP-Seq(Chong_et_al.)/Homer | AGGTCANTG | 1.00E+00 |
| Ptf1a(bHLH)/Panc1-Ptf1a-ChIP-Seq(GSE47459)/Homer | ACAGCTGTT | 1.00E+00 |
| Pax8(Paired,Homeobox)/Thyroid-Pax8-ChIP-Seq(GSE26938)/Homer | GTCATGCHT | 1.00E+00 |
| ZSCAN22(Zf)/HEK293-ZSCAN22.GFP-ChIP-Seq(GSE58341)/Homer | SMCAGTCW( | 1.00E+00 |
| PAX5(Paired,Homeobox)/GM12878-PAX5-ChIP-Seq(GSE32465)/Homer | GCAGCCAAG | 1.00E+00 |
| DUX(Homeobox)/C2C12-Dux-ChIP-Seq(GSE87279)/Homer | BCWGATTCA | 1.00E+00 |
| ZNF41(Zf)/HEK293-ZNF41.GFP-ChIP-Seq(GSE58341)/Homer | CCTCATGGT | 1.00E+00 |
| DUX4(Homeobox)/Myoblasts-DUX4.V5-ChIP-Seq(GSE75791)/Homer | NWTAAYCYA | 1.00E+00 |
| GATA:SCL(Zf,bHLH)/Ter119-SCL-ChIP-Seq(GSE18720)/Homer | CRGCTGBNG | 1.00E+00 |
| ZEB2(Zf)/SNU398-ZEB2-ChIP-Seq(GSE103048)/Homer | GNMCAGGT( | 1.00E+00 |
| JunD(bZIP)/K562-JunD-ChIP-Seq/Homer | ATGACGTCA | 1.00E+00 |
| Srebp2(bHLH)/HepG2-Srebp2-ChIP-Seq(GSE31477)/Homer | CGGTCACSC | 1.00E+00 |
| Phox2b(Homeobox)/CLBGA-PHOX2B-ChIP-Seq(GSE90683)/Homer | TTAATTNAAT | 1.00E+00 |
| TCF4(bHLH)/SHSY5Y-TCF4-ChIP-Seq(GSE96915)/Homer | SMCATCTGK | 1.00E+00 |
| Tbx21(T-box)/GM12878-TBX21-ChIP-Seq(Encode)/Homer | AGGTGTGAA | 1.00E+00 |
| E2A(bHLH),near_PU.1/Bcell-PU.1-ChIP-Seq(GSE21512)/Homer | NVCACCTGB | 1.00E+00 |
| Hnf6b(Homeobox)/LNCaP-Hnf6b-ChIP-Seq(GSE106305)/Homer | TATTGAYY | 1.00E+00 |
| PBX1(Homeobox)/MCF7-PBX1-ChIP-Seq(GSE28007)/Homer | GSCTGTCAC | 1.00E+00 |
| Barx1(Homeobox)/Stomach-Barx1.3xFlag-ChIP-Seq(GSE69483)/Homer | AAACMATTA | 1.00E+00 |
| PRDM10(Zf)/HEK293-PRDM10.eGFP-ChIP-Seq(Encode)/Homer | TGGTACATT( | 1.00E+00 |
| GATA(Zf),IR3/iTreg-Gata3-ChIP-Seq(GSE20898)/Homer | NNNNNBAGA | 1.00E+00 |
| NeuroG2(bHLH)/Fibroblast-NeuroG2-ChIP-Seq(GSE75910)/Homer | ACCATCTGT | 1.00E+00 |
| Nkx2.5(Homeobox)/HL1-Nkx2.5.biotin-ChIP-Seq(GSE21529)/Homer | RRSCACTYA | 1.00E+00 |
| HIF-1a(bHLH)/MCF7-HIF1a-ChIP-Seq(GSE28352)/Homer | TACGTGCV | 1.00E+00 |
| KLF6(Zf)/PDAC-KLF6-ChIP-Seq(GSE64557)/Homer | MKGGGYGT( | 1.00E+00 |
| Twist2(bHLH)/Myoblast-Twist2.Ty1-ChIP-Seq(GSE127998)/Homer | MCAGCTGBY | 1.00E+00 |
| Tgif1(Homeobox)/mES-Tgif1-ChIP-Seq(GSE55404)/Homer | YTGWCADY | 1.00E+00 |
| Cux2(Homeobox)/Liver-Cux2-ChIP-Seq(GSE35985)/Homer | HNRAATCAA | 1.00E+00 |
| OCT:OCT(POU,Homeobox)/NPC-OCT6-ChIP-Seq(GSE43916)/Homer | YATGCATATI | 1.00E+00 |
| Chop(bZIP)/MEF-Chop-ChIP-Seq(GSE35681)/Homer | ATTGCATCA | 1.00E+00 |
| ZNF143\|STAF(Zf)/CUTLL-ZNF143-ChIP-Seq(GSE29600)/Homer | ATTTCCCAG | 1.00E+00 |
| Gli2(Zf)/GM2-Gli2-ChIP-Chip(GSE112702)/Homer | YSTGGGTGG | 1.00E+00 |
| GFY(?)/Promoter/Homer | ACTACAATT( | 1.00E+00 |
| REST-NRSF(Zf)/Jurkat-NRSF-ChIP-Seq/Homer | GGMGCTGT( | 1.00E+00 |
| NeuroD1(bHLH)/Islet-NeuroD1-ChIP-Seq(GSE30298)/Homer | GCCATCTGT | 1.00E+00 |
| EKLF(Zf)/Erythrocyte-Klf1-ChIP-Seq(GSE20478)/Homer | NWGGGTGT( | 1.00E+00 |

| | | |
|---|---|---|
| BORIS(Zf)/K562-CTCFL-ChIP-Seq(GSE32465)/Homer | CNNBRGCGC | 1.00E+00 |
| CRE(bZIP)/Promoter/Homer | CSGTGACGT | 1.00E+00 |
| CREB5(bZIP)/LNCaP-CREB5.V5-ChIP-Seq(GSE137775)/Homer | VVATGACGT | 1.00E+00 |
| Srebp1a(bHLH)/HepG2-Srebp1a-ChIP-Seq(GSE31477)/Homer | RTCACSCCA | 1.00E+00 |
| Nkx2.1(Homeobox)/LungAC-Nkx2.1-ChIP-Seq(GSE43252)/Homer | RSCACTYRA | 1.00E+00 |
| NRF1(NRF)/MCF7-NRF1-ChIP-Seq(Unpublished)/Homer | CTGCGCATG | 1.00E+00 |
| c-Jun-CRE(bZIP)/K562-cJun-ChIP-Seq(GSE31477)/Homer | ATGACGTCA | 1.00E+00 |
| Bcl11a(Zf)/HSPC-BCL11A-ChIP-Seq(GSE104676)/Homer | TYTGACCAS | 1.00E+00 |
| Phox2a(Homeobox)/Neuron-Phox2a-ChIP-Seq(GSE31456)/Homer | YTAATYNRA | 1.00E+00 |
| SCRT1(Zf)/HEK293-SCRT1.eGFP-ChIP-Seq(Encode)/Homer | GCAACAGGT | 1.00E+00 |
| Nkx3.1(Homeobox)/LNCaP-Nkx3.1-ChIP-Seq(GSE28264)/Homer | AAGCACTTA | 1.00E+00 |
| Nkx2.2(Homeobox)/NPC-Nkx2.2-ChIP-Seq(GSE61673)/Homer | BTBRAGTGS | 1.00E+00 |
| HOXA2(Homeobox)/mES-Hoxa2-ChIP-Seq(Donaldson_et_al.)/Homer | GYCATCMAT | 1.00E+00 |
| Duxbl(Homeobox)/NIH3T3-Duxbl.HA-ChIP-Seq(GSE119782)/Homer | TAAYCYAATC | 1.00E+00 |
| Npas4(bHLH)/Neuron-Npas4-ChIP-Seq(GSE127793)/Homer | NHRTCACGA | 1.00E+00 |
| Pax7(Paired,Homeobox),longest/Myoblast-Pax7-ChIP-Seq(GSE25064)/Homer | NTAATTDGC | 1.00E+00 |
| CTCF(Zf)/CD4+-CTCF-ChIP-Seq(Barski_et_al.)/Homer | AYAGTGCCM | 1.00E+00 |
| TFE3(bHLH)/MEF-TFE3-ChIP-Seq(GSE75757)/Homer | GTCACGTGA | 1.00E+00 |
| Twist(bHLH)/HMLE-TWIST1-ChIP-Seq(Chang_et_al)/Homer | VCAKCTGGN | 1.00E+00 |
| Ascl2(bHLH)/ESC-Ascl2-ChIP-Seq(GSE97712)/Homer | SSRGCAGCT | 1.00E+00 |
| KLF1(Zf)/HUDEP2-KLF1-CutnRun(GSE136251)/Homer | VDGGGYGGC | 1.00E+00 |
| NPAS2(bHLH)/Liver-NPAS2-ChIP-Seq(GSE39860)/Homer | KCCACGTGA | 1.00E+00 |
| Tbet(T-box)/CD8-Tbet-ChIP-Seq(GSE33802)/Homer | AGGTGTGAA | 1.00E+00 |
| Prop1(Homeobox)/GHFT1-PROP1.biotin-ChIP-Seq(GSE77302)/Homer | NTAATBNAA | 1.00E+00 |
| MITF(bHLH)/MastCells-MITF-ChIP-Seq(GSE48085)/Homer | RTCATGTGA | 1.00E+00 |
| CUX1(Homeobox)/K562-CUX1-ChIP-Seq(GSE92882)/Homer | TATCGATNA | 1.00E+00 |
| HOXA1(Homeobox)/mES-Hoxa1-ChIP-Seq(SRP084292)/Homer | TGATKGATG | 1.00E+00 |
| Hoxd12(Homeobox)/ChickenMSG-Hoxd12.Flag-ChIP-Seq(GSE86088)/Homer | HDGYAATGA | 1.00E+00 |
| E-box(bHLH)/Promoter/Homer | SSGGTCACG | 1.00E+00 |
| GSC(Homeobox)/FrogEmbryos-GSC-ChIP-Seq(DRA000576)/Homer | RGGATTAR | 1.00E+00 |
| THRb(NR)/HepG2-THRb.Flag-ChIP-Seq(Encode)/Homer | GGTCACCTG | 1.00E+00 |
| Snail1(Zf)/LS174T-SNAIL1.HA-ChIP-Seq(GSE127183)/Homer | TRCACCTGC | 1.00E+00 |
| NPAS(bHLH)/Liver-NPAS-ChIP-Seq(GSE39860)/Homer | NVCACGTG | 1.00E+00 |
| bHLHE41(bHLH)/proB-Bhlhe41-ChIP-Seq(GSE93764)/Homer | KCACGTGMC | 1.00E+00 |
| Ronin(THAP)/ES-Thap11-ChIP-Seq(GSE51522)/Homer | RACTACAAC | 1.00E+00 |
| BMAL1(bHLH)/Liver-Bmal1-ChIP-Seq(GSE39860)/Homer | GNCACGTG | 1.00E+00 |
| Gfi1b(Zf)/HPC7-Gfi1b-ChIP-Seq(GSE22178)/Homer | MAATCACTG | 1.00E+00 |
| Atf2(bZIP)/3T3L1-Atf2-ChIP-Seq(GSE56872)/Homer | NRRTGACGT | 1.00E+00 |
| ZNF136(Zf)/HEK293-ZNF136.GFP-ChIP-Seq(GSE58341)/Homer | YTKGATAHA | 1.00E+00 |
| KLF10(Zf)/HEK293-KLF10.GFP-ChIP-Seq(GSE58341)/Homer | GGGGGTGTC | 1.00E+00 |
| Mef2b(MADS)/HEK293-Mef2b.V5-ChIP-Seq(GSE67450)/Homer | GCTATTTTTC | 1.00E+00 |
| MNT(bHLH)/HepG2-MNT-ChIP-Seq(Encode)/Homer | DGCACACGT | 1.00E+00 |
| Bapx1(Homeobox)/VertebralCol-Bapx1-ChIP-Seq(GSE36672)/Homer | TTRAGTGSY | 1.00E+00 |
| THRa(NR)/C17.2-THRa-ChIP-Seq(GSE38347)/Homer | GGTCANYTG | 1.00E+00 |
| bHLHE40(bHLH)/HepG2-BHLHE40-ChIP-Seq(GSE31477)/Homer | KCACGTGMC | 1.00E+00 |
| c-Myc(bHLH)/mES-cMyc-ChIP-Seq(GSE11431)/Homer | VVCCACGTG | 1.00E+00 |
| n-Myc(bHLH)/mES-nMyc-ChIP-Seq(GSE11431)/Homer | VRCCACGTG | 1.00E+00 |
| c-Myc(bHLH)/LNCAP-cMyc-ChIP-Seq(Unpublished)/Homer | VCCACGTG | 1.00E+00 |
| Max(bHLH)/K562-Max-ChIP-Seq(GSE31477)/Homer | RCCACGTGG | 1.00E+00 |
| LXRE(NR),DR4/RAW-LXRb.biotin-ChIP-Seq(GSE21512)/Homer | RGGTTACTA | 1.00E+00 |
| Sp1(Zf)/Promoter/Homer | GGCCCCGCC | 1.00E+00 |
| KLF3(Zf)/MEF-Klf3-ChIP-Seq(GSE44748)/Homer | NRGCCCCRC | 1.00E+00 |
| NFY(CCAAT)/Promoter/Homer | RGCCAATSR | 1.00E+00 |
| Klf9(Zf)/GBM-Klf9-ChIP-Seq(GSE62211)/Homer | GCCACRCCC | 1.00E+00 |

| | | |
|---|---|---|
| GLIS3(Zf)/Thyroid-Glis3.GFP-ChIP-Seq(GSE103297)/Homer | CTCCCTGGG | 1.00E+00 |
| Mef2c(MADS)/GM12878-Mef2c-ChIP-Seq(GSE32465)/Homer | DCYAAAAATA | 1.00E+00 |
| Usf2(bHLH)/C2C12-Usf2-ChIP-Seq(GSE36030)/Homer | GTCACGTGG | 1.00E+00 |
| Atf7(bZIP)/3T3L1-Atf7-ChIP-Seq(GSE56872)/Homer | NGRTGACGT | 1.00E+00 |
| Mef2a(MADS)/HL1-Mef2a.biotin-ChIP-Seq(GSE21529)/Homer | CYAAAAATA( | 1.00E+00 |
| Atf1(bZIP)/K562-ATF1-ChIP-Seq(GSE31477)/Homer | GATGACGTC | 1.00E+00 |
| USF1(bHLH)/GM12878-Usf1-ChIP-Seq(GSE32465)/Homer | SGTCACGTG | 1.00E+00 |
| Klf4(Zf)/mES-Klf4-ChIP-Seq(GSE11431)/Homer | GCCACACCC | 1.00E+00 |
| Slug(Zf)/Mesoderm-Snai2-ChIP-Seq(GSE61475)/Homer | SNGCACCTG | 1.00E+00 |
| CLOCK(bHLH)/Liver-Clock-ChIP-Seq(GSE39860)/Homer | GHCACGTG | 1.00E+00 |

**from LuCap-147-CR post treatment with dTRIM24-1**

| Log P-value | q-value (Benj | # of Target S | % of Target S | # of Backgro | % of Background Sequenc |
|---|---|---|---|---|---|
| -2.30E+02 | 0 | 554 | 3.53% | 419 | 1.23% |
| -2.28E+02 | 0 | 871 | 5.55% | 848.1 | 2.50% |
| -1.32E+02 | 0 | 266 | 1.70% | 179.2 | 0.53% |
| -1.08E+02 | 0 | 869 | 5.54% | 1117 | 3.29% |
| -8.24E+01 | 0 | 4431 | 28.24% | 8112.9 | 23.88% |
| -8.10E+01 | 0 | 5357 | 34.14% | 10036.7 | 29.54% |
| -8.02E+01 | 0 | 1060 | 6.76% | 1546.4 | 4.55% |
| -7.94E+01 | 0 | 4910 | 31.30% | 9125.4 | 26.86% |
| -7.50E+01 | 0 | 2511 | 16.00% | 4319 | 12.71% |
| -7.48E+01 | 0 | 2546 | 16.23% | 4388.6 | 12.92% |
| -7.18E+01 | 0 | 868 | 5.53% | 1240.4 | 3.65% |
| -7.04E+01 | 0 | 3943 | 25.13% | 7225.2 | 21.27% |
| -6.97E+01 | 0 | 2438 | 15.54% | 4214.3 | 12.41% |
| -6.96E+01 | 0 | 1931 | 12.31% | 3228.8 | 9.50% |
| -6.95E+01 | 0 | 5839 | 37.22% | 11162.5 | 32.86% |
| -6.94E+01 | 0 | 2956 | 18.84% | 5243 | 15.43% |
| -6.78E+01 | 0 | 3567 | 22.74% | 6485.9 | 19.09% |
| -6.77E+01 | 0 | 6799 | 43.34% | 13212.5 | 38.89% |
| -6.68E+01 | 0 | 5654 | 36.04% | 10803.1 | 31.80% |
| -6.45E+01 | 0 | 4589 | 29.25% | 8610.5 | 25.35% |
| -6.36E+01 | 0 | 3553 | 22.65% | 6496 | 19.12% |
| -6.26E+01 | 0 | 4791 | 30.54% | 9048.5 | 26.64% |
| -6.08E+01 | 0 | 4445 | 28.33% | 8351.7 | 24.58% |
| -6.01E+01 | 0 | 6154 | 39.22% | 11933.1 | 35.13% |
| -5.83E+01 | 0 | 4204 | 26.80% | 7881.5 | 23.20% |
| -5.72E+01 | 0 | 3270 | 20.84% | 5982 | 17.61% |
| -5.45E+01 | 0 | 1881 | 11.99% | 3237.2 | 9.53% |
| -5.37E+01 | 0 | 1134 | 7.23% | 1811.2 | 5.33% |
| -5.37E+01 | 0 | 4225 | 26.93% | 7974.3 | 23.47% |
| -5.12E+01 | 0 | 4453 | 28.38% | 8475.8 | 24.95% |
| -5.09E+01 | 0 | 1114 | 7.10% | 1791 | 5.27% |
| -5.06E+01 | 0 | 465 | 2.96% | 621.5 | 1.83% |
| -4.78E+01 | 0 | 2464 | 15.71% | 4447.6 | 13.09% |
| -4.72E+01 | 0 | 1282 | 8.17% | 2131.2 | 6.27% |
| -4.64E+01 | 0 | 2631 | 16.77% | 4796.8 | 14.12% |
| -3.82E+01 | 0 | 5884 | 37.50% | 11653.3 | 34.30% |
| -3.80E+01 | 0 | 4394 | 28.01% | 8518.3 | 25.07% |
| -3.77E+01 | 0 | 10282 | 65.54% | 21175.8 | 62.33% |
| -3.74E+01 | 0 | 1857 | 11.84% | 3332.4 | 9.81% |
| -3.60E+01 | 0 | 3751 | 23.91% | 7207 | 21.21% |
| -3.58E+01 | 0 | 1681 | 10.71% | 2997.1 | 8.82% |
| -3.49E+01 | 0 | 1658 | 10.57% | 2960.8 | 8.72% |
| -3.45E+01 | 0 | 2255 | 14.37% | 4160.2 | 12.25% |
| -3.34E+01 | 0 | 2401 | 15.30% | 4467.6 | 13.15% |
| -3.22E+01 | 0 | 2109 | 13.44% | 3890.2 | 11.45% |
| -3.19E+01 | 0 | 5011 | 31.94% | 9903.7 | 29.15% |
| -3.18E+01 | 0 | 2672 | 17.03% | 5039 | 14.83% |
| -3.13E+01 | 0 | 4412 | 28.12% | 8653.7 | 25.47% |
| -3.02E+01 | 0 | 2297 | 14.64% | 4293 | 12.64% |
| -2.92E+01 | 0 | 2262 | 14.42% | 4234.1 | 12.46% |
| -2.81E+01 | 0 | 11162 | 71.15% | 23281.9 | 68.53% |
| -2.72E+01 | 0 | 5940 | 37.86% | 11953.3 | 35.19% |

| | | | | | |
|---|---|---|---|---|---|
| -2.66E+01 | 0 | 9438 | 60.16% | 19514.3 | 57.44% |
| -2.62E+01 | 0 | 5349 | 34.09% | 10714.4 | 31.54% |
| -2.62E+01 | 0 | 5775 | 36.81% | 11620.1 | 34.21% |
| -2.58E+01 | 0 | 2396 | 15.27% | 4549.2 | 13.39% |
| -2.54E+01 | 0 | 4482 | 28.57% | 8896.1 | 26.19% |
| -2.47E+01 | 0 | 4610 | 29.38% | 9178.7 | 27.02% |
| -2.44E+01 | 0 | 2340 | 14.91% | 4453.3 | 13.11% |
| -2.40E+01 | 0 | 11727 | 74.75% | 24609.1 | 72.44% |
| -2.32E+01 | 0 | 3032 | 19.33% | 5899.6 | 17.37% |
| -2.30E+01 | 0 | 2390 | 15.23% | 4576 | 13.47% |
| -2.29E+01 | 0 | 4003 | 25.51% | 7932 | 23.35% |
| -2.21E+01 | 0 | 1910 | 12.17% | 3605.9 | 10.61% |
| -2.21E+01 | 0 | 598 | 3.81% | 997.3 | 2.94% |
| -2.20E+01 | 0 | 1980 | 12.62% | 3749.6 | 11.04% |
| -2.17E+01 | 0 | 1146 | 7.30% | 2070.4 | 6.09% |
| -2.07E+01 | 0 | 2678 | 17.07% | 5204.2 | 15.32% |
| -1.93E+01 | 0 | 6043 | 38.52% | 12330.2 | 36.30% |
| -1.89E+01 | 0 | 5210 | 33.21% | 10562.7 | 31.09% |
| -1.85E+01 | 0 | 394 | 2.51% | 636 | 1.87% |
| -1.84E+01 | 0 | 1797 | 11.45% | 3424.1 | 10.08% |
| -1.78E+01 | 0 | 2009 | 12.81% | 3867.2 | 11.38% |
| -1.77E+01 | 0 | 2327 | 14.83% | 4525.4 | 13.32% |
| -1.70E+01 | 0 | 6663 | 42.47% | 13710.4 | 40.36% |
| -1.68E+01 | 0 | 1844 | 11.75% | 3543.9 | 10.43% |
| -1.66E+01 | 0 | 4018 | 25.61% | 8083.1 | 23.79% |
| -1.60E+01 | 0 | 1693 | 10.79% | 3245.9 | 9.55% |
| -1.59E+01 | 0 | 9949 | 63.41% | 20863.8 | 61.42% |
| -1.58E+01 | 0 | 3518 | 22.42% | 7047.5 | 20.75% |
| -1.57E+01 | 0 | 1663 | 10.60% | 3187.5 | 9.38% |
| -1.56E+01 | 0 | 5323 | 33.93% | 10877.6 | 32.02% |
| -1.51E+01 | 0 | 5472 | 34.88% | 11205.4 | 32.98% |
| -1.49E+01 | 0 | 1466 | 9.34% | 2796.5 | 8.23% |
| -1.39E+01 | 0 | 977 | 6.23% | 1817.4 | 5.35% |
| -1.30E+01 | 0 | 5588 | 35.62% | 11508.8 | 33.88% |
| -1.30E+01 | 0 | 4053 | 25.83% | 8239.5 | 24.25% |
| -1.29E+01 | 0 | 3651 | 23.27% | 7389.6 | 21.75% |
| -1.27E+01 | 0 | 349 | 2.22% | 588.8 | 1.73% |
| -1.27E+01 | 0 | 349 | 2.22% | 588.8 | 1.73% |
| -1.27E+01 | 0 | 2667 | 17.00% | 5324.3 | 15.67% |
| -1.25E+01 | 0 | 8271 | 52.72% | 17302.6 | 50.93% |
| -1.22E+01 | 0 | 1397 | 8.90% | 2694.8 | 7.93% |
| -1.20E+01 | 0 | 254 | 1.62% | 413.5 | 1.22% |
| -1.19E+01 | 0 | 514 | 3.28% | 915.7 | 2.70% |
| -1.17E+01 | 0 | 1230 | 7.84% | 2360.4 | 6.95% |
| -1.13E+01 | 0.0001 | 1194 | 7.61% | 2293.7 | 6.75% |
| -1.10E+01 | 0.0001 | 1846 | 11.77% | 3643.8 | 10.73% |
| -1.10E+01 | 0.0001 | 797 | 5.08% | 1489.4 | 4.38% |
| -1.09E+01 | 0.0001 | 3026 | 19.29% | 6118 | 18.01% |
| -1.08E+01 | 0.0001 | 1173 | 7.48% | 2257.5 | 6.65% |
| -1.04E+01 | 0.0001 | 1156 | 7.37% | 2228.3 | 6.56% |
| -1.03E+01 | 0.0002 | 8364 | 53.31% | 17572 | 51.73% |
| -1.01E+01 | 0.0002 | 879 | 5.60% | 1667.5 | 4.91% |
| -1.01E+01 | 0.0002 | 248 | 1.58% | 414.1 | 1.22% |
| -1.00E+01 | 0.0002 | 1281 | 8.16% | 2491 | 7.33% |

| | | | | | |
|---|---|---|---|---|---|
| -1.00E+01 | 0.0002 | 781 | 4.98% | 1469.7 | 4.33% |
| -9.93E+00 | 0.0002 | 1108 | 7.06% | 2137.2 | 6.29% |
| -9.87E+00 | 0.0002 | 2650 | 16.89% | 5350.6 | 15.75% |
| -9.80E+00 | 0.0002 | 1066 | 6.79% | 2053.3 | 6.04% |
| -9.68E+00 | 0.0002 | 1595 | 10.17% | 3147.7 | 9.27% |
| -9.67E+00 | 0.0002 | 129 | 0.82% | 195.8 | 0.58% |
| -9.49E+00 | 0.0003 | 2935 | 18.71% | 5960.5 | 17.55% |
| -9.37E+00 | 0.0003 | 1861 | 11.86% | 3707.9 | 10.91% |
| -9.14E+00 | 0.0004 | 785 | 5.00% | 1489.2 | 4.38% |
| -9.01E+00 | 0.0005 | 2491 | 15.88% | 5037 | 14.83% |
| -8.96E+00 | 0.0005 | 1288 | 8.21% | 2524.3 | 7.43% |
| -8.60E+00 | 0.0007 | 931 | 5.93% | 1795.6 | 5.29% |
| -8.36E+00 | 0.0009 | 728 | 4.64% | 1385 | 4.08% |
| -8.30E+00 | 0.0009 | 3102 | 19.77% | 6345.2 | 18.68% |
| -8.24E+00 | 0.001 | 917 | 5.84% | 1772.5 | 5.22% |
| -8.16E+00 | 0.001 | 4097 | 26.11% | 8464.9 | 24.92% |
| -8.10E+00 | 0.0011 | 214 | 1.36% | 363.5 | 1.07% |
| -7.97E+00 | 0.0012 | 2792 | 17.80% | 5698.8 | 16.78% |
| -7.96E+00 | 0.0012 | 452 | 2.88% | 832.4 | 2.45% |
| -7.90E+00 | 0.0013 | 8639 | 55.06% | 18249.2 | 53.72% |
| -7.23E+00 | 0.0025 | 613 | 3.91% | 1166.2 | 3.43% |
| -7.11E+00 | 0.0028 | 1515 | 9.66% | 3033.1 | 8.93% |
| -7.11E+00 | 0.0028 | 1328 | 8.46% | 2643.9 | 7.78% |
| -6.99E+00 | 0.0031 | 4063 | 25.90% | 8430.5 | 24.82% |
| -6.81E+00 | 0.0037 | 21 | 0.13% | 21 | 0.06% |
| -6.77E+00 | 0.0038 | 1142 | 7.28% | 2263.6 | 6.66% |
| -6.66E+00 | 0.0043 | 2682 | 17.09% | 5503.2 | 16.20% |
| -6.61E+00 | 0.0044 | 3426 | 21.84% | 7085.2 | 20.86% |
| -6.54E+00 | 0.0047 | 385 | 2.45% | 714.8 | 2.10% |
| -6.43E+00 | 0.0052 | 2254 | 14.37% | 4604.9 | 13.56% |
| -6.39E+00 | 0.0054 | 2385 | 15.20% | 4882.9 | 14.37% |
| -6.38E+00 | 0.0054 | 3514 | 22.40% | 7280.1 | 21.43% |
| -6.25E+00 | 0.0061 | 1299 | 8.28% | 2601.2 | 7.66% |
| -6.12E+00 | 0.0069 | 2121 | 13.52% | 4332.9 | 12.75% |
| -6.01E+00 | 0.0077 | 569 | 3.63% | 1094.9 | 3.22% |
| -5.99E+00 | 0.0078 | 513 | 3.27% | 980.4 | 2.89% |
| -5.95E+00 | 0.008 | 724 | 4.61% | 1413.9 | 4.16% |
| -5.89E+00 | 0.0085 | 1625 | 10.36% | 3293.4 | 9.69% |
| -5.75E+00 | 0.0097 | 1638 | 10.44% | 3324.9 | 9.79% |
| -5.69E+00 | 0.0102 | 1332 | 8.49% | 2683.5 | 7.90% |
| -5.68E+00 | 0.0102 | 3883 | 24.75% | 8093.4 | 23.82% |
| -5.61E+00 | 0.0109 | 585 | 3.73% | 1133.6 | 3.34% |
| -5.38E+00 | 0.0137 | 1031 | 6.57% | 2061.2 | 6.07% |
| -5.31E+00 | 0.0145 | 11002 | 70.13% | 23499.6 | 69.17% |
| -4.96E+00 | 0.0204 | 4997 | 31.85% | 10510.6 | 30.94% |
| -4.96E+00 | 0.0204 | 4526 | 28.85% | 9499.2 | 27.96% |
| -4.75E+00 | 0.0249 | 4911 | 31.30% | 10335.4 | 30.42% |
| -4.67E+00 | 0.0269 | 1500 | 9.56% | 3063.3 | 9.02% |
| -4.66E+00 | 0.0269 | 2372 | 15.12% | 4910.6 | 14.46% |
| -4.64E+00 | 0.0272 | 116 | 0.74% | 200.2 | 0.59% |
| -4.45E+00 | 0.0326 | 478 | 3.05% | 932.8 | 2.75% |
| -4.29E+00 | 0.038 | 901 | 5.74% | 1814.9 | 5.34% |
| -4.23E+00 | 0.0402 | 4544 | 28.96% | 9571.4 | 28.17% |
| -4.21E+00 | 0.041 | 243 | 1.55% | 456.6 | 1.34% |

| | | | | | |
|---|---|---|---|---|---|
| -4.19E+00 | 0.0415 | 2910 | 18.55% | 6074.2 | 17.88% |
| -4.12E+00 | 0.0442 | 2581 | 16.45% | 5375.9 | 15.82% |
| -4.12E+00 | 0.0442 | 696 | 4.44% | 1390.5 | 4.09% |
| -4.11E+00 | 0.0442 | 1425 | 9.08% | 2921 | 8.60% |
| -4.11E+00 | 0.0442 | 731 | 4.66% | 1463.3 | 4.31% |
| -4.11E+00 | 0.0442 | 4543 | 28.96% | 9575.5 | 28.19% |
| -4.11E+00 | 0.0442 | 4064 | 25.90% | 8547.9 | 25.16% |
| -4.08E+00 | 0.0444 | 2094 | 13.35% | 4340.4 | 12.78% |
| -4.01E+00 | 0.0471 | 1649 | 10.51% | 3398.1 | 10.00% |
| -4.00E+00 | 0.0475 | 4514 | 28.77% | 9518.8 | 28.02% |
| -3.88E+00 | 0.0531 | 880 | 5.61% | 1780.2 | 5.24% |
| -3.88E+00 | 0.0531 | 506 | 3.23% | 1000.9 | 2.95% |
| -3.88E+00 | 0.0531 | 533 | 3.40% | 1056.6 | 3.11% |
| -3.86E+00 | 0.0532 | 556 | 3.54% | 1104.2 | 3.25% |
| -3.85E+00 | 0.0535 | 699 | 4.46% | 1402.4 | 4.13% |
| -3.80E+00 | 0.0562 | 9053 | 57.70% | 19332.7 | 56.91% |
| -3.76E+00 | 0.0577 | 2465 | 15.71% | 5142 | 15.14% |
| -3.68E+00 | 0.0621 | 536 | 3.42% | 1067 | 3.14% |
| -3.65E+00 | 0.0642 | 8421 | 53.67% | 17970.2 | 52.90% |
| -3.64E+00 | 0.0644 | 574 | 3.66% | 1146.7 | 3.38% |
| -3.42E+00 | 0.0793 | 167 | 1.06% | 312.6 | 0.92% |
| -3.41E+00 | 0.0802 | 1327 | 8.46% | 2736.7 | 8.06% |
| -3.10E+00 | 0.1085 | 1340 | 8.54% | 2774.9 | 8.17% |
| -3.05E+00 | 0.1137 | 2808 | 17.90% | 5907.7 | 17.39% |
| -2.98E+00 | 0.1213 | 1418 | 9.04% | 2944.8 | 8.67% |
| -2.85E+00 | 0.1367 | 1468 | 9.36% | 3055.6 | 8.99% |
| -2.82E+00 | 0.1398 | 371 | 2.36% | 740.4 | 2.18% |
| -2.76E+00 | 0.1481 | 4249 | 27.08% | 9016.5 | 26.54% |
| -2.76E+00 | 0.1481 | 403 | 2.57% | 809 | 2.38% |
| -2.69E+00 | 0.1574 | 3587 | 22.86% | 7597.9 | 22.37% |
| -2.69E+00 | 0.1574 | 3693 | 23.54% | 7825.7 | 23.04% |
| -2.65E+00 | 0.161 | 232 | 1.48% | 455.8 | 1.34% |
| -2.65E+00 | 0.161 | 232 | 1.48% | 455.8 | 1.34% |
| -2.65E+00 | 0.161 | 121 | 0.77% | 228.2 | 0.67% |
| -2.63E+00 | 0.1625 | 227 | 1.45% | 445.4 | 1.31% |
| -2.53E+00 | 0.1784 | 2426 | 15.46% | 5115.5 | 15.06% |
| -2.49E+00 | 0.1847 | 860 | 5.48% | 1777.7 | 5.23% |
| -2.41E+00 | 0.199 | 2757 | 17.57% | 5831.4 | 17.17% |
| -2.34E+00 | 0.2132 | 1423 | 9.07% | 2980.1 | 8.77% |
| -2.31E+00 | 0.2193 | 383 | 2.44% | 776.1 | 2.28% |
| -2.29E+00 | 0.2209 | 5568 | 35.49% | 11890.8 | 35.00% |
| -2.29E+00 | 0.2209 | 1998 | 12.74% | 4211.2 | 12.40% |
| -2.27E+00 | 0.2251 | 618 | 3.94% | 1272.6 | 3.75% |
| -2.21E+00 | 0.2362 | 799 | 5.09% | 1657.9 | 4.88% |
| -2.15E+00 | 0.2496 | 121 | 0.77% | 234.2 | 0.69% |
| -2.13E+00 | 0.2545 | 3566 | 22.73% | 7587.9 | 22.34% |
| -2.12E+00 | 0.2545 | 330 | 2.10% | 669.3 | 1.97% |
| -2.12E+00 | 0.2545 | 6349 | 40.47% | 13590.8 | 40.01% |
| -2.10E+00 | 0.2591 | 24 | 0.15% | 40.2 | 0.12% |
| -2.02E+00 | 0.2793 | 2153 | 13.72% | 4558.5 | 13.42% |
| -1.98E+00 | 0.2887 | 115 | 0.73% | 224.1 | 0.66% |
| -1.92E+00 | 0.3055 | 433 | 2.76% | 891.9 | 2.63% |
| -1.88E+00 | 0.3156 | 4548 | 28.99% | 9721.4 | 28.62% |
| -1.84E+00 | 0.3254 | 1366 | 8.71% | 2881.8 | 8.48% |

| | | | | | |
|---|---|---|---|---|---|
| -1.83E+00 | 0.3296 | 1281 | 8.16% | 2700.1 | 7.95% |
| -1.81E+00 | 0.3333 | 110 | 0.70% | 216.6 | 0.64% |
| -1.81E+00 | 0.3333 | 1389 | 8.85% | 2932.5 | 8.63% |
| -1.80E+00 | 0.3333 | 123 | 0.78% | 243.8 | 0.72% |
| -1.79E+00 | 0.3352 | 313 | 2.00% | 641.3 | 1.89% |
| -1.77E+00 | 0.3424 | 1006 | 6.41% | 2115.2 | 6.23% |
| -1.76E+00 | 0.3434 | 1084 | 6.91% | 2282.4 | 6.72% |
| -1.74E+00 | 0.3472 | 356 | 2.27% | 733.5 | 2.16% |
| -1.73E+00 | 0.3489 | 2971 | 18.94% | 6334.7 | 18.65% |
| -1.64E+00 | 0.3814 | 441 | 2.81% | 916.8 | 2.70% |
| -1.59E+00 | 0.4006 | 152 | 0.97% | 307.2 | 0.90% |
| -1.58E+00 | 0.4006 | 61 | 0.39% | 119 | 0.35% |
| -1.55E+00 | 0.4121 | 268 | 1.71% | 552.9 | 1.63% |
| -1.44E+00 | 0.4589 | 3371 | 21.49% | 7219.7 | 21.25% |
| -1.40E+00 | 0.4727 | 202 | 1.29% | 416.2 | 1.23% |
| -1.36E+00 | 0.4896 | 2800 | 17.85% | 5994.1 | 17.64% |
| -1.33E+00 | 0.5061 | 293 | 1.87% | 611.4 | 1.80% |
| -1.32E+00 | 0.5088 | 1413 | 9.01% | 3011.6 | 8.87% |
| -1.26E+00 | 0.5342 | 8065 | 51.41% | 17384.8 | 51.17% |
| -1.22E+00 | 0.556 | 644 | 4.10% | 1365.8 | 4.02% |
| -1.21E+00 | 0.5611 | 556 | 3.54% | 1177.9 | 3.47% |
| -1.20E+00 | 0.5626 | 2596 | 16.55% | 5568.4 | 16.39% |
| -1.20E+00 | 0.5626 | 1251 | 7.97% | 2670.2 | 7.86% |
| -1.16E+00 | 0.5808 | 358 | 2.28% | 755.5 | 2.22% |
| -1.14E+00 | 0.5865 | 1156 | 7.37% | 2469.9 | 7.27% |
| -1.10E+00 | 0.6106 | 1931 | 12.31% | 4142.2 | 12.19% |
| -1.08E+00 | 0.6182 | 8967 | 57.15% | 19359.8 | 56.99% |
| -1.07E+00 | 0.6254 | 6484 | 41.33% | 13985.7 | 41.17% |
| -1.06E+00 | 0.6257 | 1122 | 7.15% | 2401.5 | 7.07% |
| -1.04E+00 | 0.6369 | 3107 | 19.80% | 6686.5 | 19.68% |
| -1.03E+00 | 0.6413 | 40 | 0.25% | 81.7 | 0.24% |
| -1.00E+00 | 0.6553 | 80 | 0.51% | 166.3 | 0.49% |
| -9.78E-01 | 0.67 | 43 | 0.27% | 88.3 | 0.26% |
| -9.75E-01 | 0.67 | 1467 | 9.35% | 3151.5 | 9.28% |
| -9.68E-01 | 0.6712 | 23 | 0.15% | 46.7 | 0.14% |
| -9.57E-01 | 0.6761 | 133 | 0.85% | 280.2 | 0.82% |
| -9.53E-01 | 0.6761 | 1947 | 12.41% | 4189.1 | 12.33% |
| -9.48E-01 | 0.6767 | 1548 | 9.87% | 3328.1 | 9.80% |
| -9.37E-01 | 0.6811 | 369 | 2.35% | 787.4 | 2.32% |
| -9.12E-01 | 0.6959 | 1636 | 10.43% | 3521 | 10.36% |
| -9.02E-01 | 0.7002 | 548 | 3.49% | 1174.7 | 3.46% |
| -8.96E-01 | 0.7015 | 1471 | 9.38% | 3166.7 | 9.32% |
| -8.95E-01 | 0.7015 | 8038 | 51.23% | 17373 | 51.14% |
| -8.92E-01 | 0.7015 | 1243 | 7.92% | 2674.8 | 7.87% |
| -8.43E-01 | 0.7313 | 1685 | 10.74% | 3633.6 | 10.70% |
| -8.31E-01 | 0.7369 | 2474 | 15.77% | 5340.7 | 15.72% |
| -8.13E-01 | 0.7475 | 3136 | 19.99% | 6774.7 | 19.94% |
| -8.05E-01 | 0.7509 | 9884 | 63.00% | 21383.6 | 62.95% |
| -7.85E-01 | 0.7634 | 1811 | 11.54% | 3911.5 | 11.51% |
| -7.79E-01 | 0.7652 | 8 | 0.05% | 17 | 0.05% |
| -7.74E-01 | 0.7658 | 134 | 0.85% | 287.2 | 0.85% |
| -7.74E-01 | 0.7658 | 548 | 3.49% | 1181.1 | 3.48% |
| -7.60E-01 | 0.7707 | 106 | 0.68% | 227.3 | 0.67% |
| -7.33E-01 | 0.7885 | 1316 | 8.39% | 2845.7 | 8.38% |

| | | | | | |
|---|---|---|---|---|---|
| -7.22E-01 | 0.7945 | 6273 | 39.98% | 13577 | 39.97% |
| -6.98E-01 | 0.8113 | 5165 | 32.92% | 11182.1 | 32.92% |
| -6.90E-01 | 0.8142 | 4801 | 30.60% | 10395.7 | 30.60% |
| -6.73E-01 | 0.8252 | 1849 | 11.79% | 4005.2 | 11.79% |
| -6.55E-01 | 0.8372 | 1957 | 12.47% | 4241.1 | 12.48% |
| -6.52E-01 | 0.8372 | 420 | 2.68% | 911.9 | 2.68% |
| -6.40E-01 | 0.8437 | 416 | 2.65% | 903.1 | 2.66% |
| -6.25E-01 | 0.8534 | 45 | 0.29% | 98.1 | 0.29% |
| -6.10E-01 | 0.8633 | 960 | 6.12% | 2085.3 | 6.14% |
| -6.04E-01 | 0.8655 | 212 | 1.35% | 462.9 | 1.36% |
| -5.60E-01 | 0.9006 | 354 | 2.26% | 773.7 | 2.28% |
| -5.32E-01 | 0.923 | 5136 | 32.74% | 11148.3 | 32.82% |
| -5.32E-01 | 0.923 | 367 | 2.34% | 803.8 | 2.37% |
| -4.89E-01 | 0.9567 | 819 | 5.22% | 1790 | 5.27% |
| -4.89E-01 | 0.9567 | 1244 | 7.93% | 2714.8 | 7.99% |
| -4.81E-01 | 0.9574 | 224 | 1.43% | 494.1 | 1.45% |
| -4.80E-01 | 0.9574 | 98 | 0.62% | 219 | 0.64% |
| -4.44E-01 | 0.9874 | 537 | 3.42% | 1180.5 | 3.48% |
| -4.25E-01 | 1 | 323 | 2.06% | 714.8 | 2.10% |
| -4.22E-01 | 1 | 1654 | 10.54% | 3614.8 | 10.64% |
| -3.79E-01 | 1 | 3870 | 24.67% | 8435.5 | 24.83% |
| -3.73E-01 | 1 | 1054 | 6.72% | 2315.8 | 6.82% |
| -3.72E-01 | 1 | 111 | 0.71% | 251.6 | 0.74% |
| -3.71E-01 | 1 | 2611 | 16.64% | 5703.5 | 16.79% |
| -3.31E-01 | 1 | 5965 | 38.02% | 12991.6 | 38.24% |
| -3.20E-01 | 1 | 215 | 1.37% | 484.3 | 1.43% |
| -3.14E-01 | 1 | 416 | 2.65% | 927.1 | 2.73% |
| -3.00E-01 | 1 | 614 | 3.91% | 1363.8 | 4.01% |
| -2.80E-01 | 1 | 8193 | 52.22% | 17833.4 | 52.50% |
| -2.64E-01 | 1 | 2431 | 15.49% | 5335.2 | 15.71% |
| -2.57E-01 | 1 | 1537 | 9.80% | 3388.9 | 9.98% |
| -2.57E-01 | 1 | 510 | 3.25% | 1140.2 | 3.36% |
| -2.37E-01 | 1 | 295 | 1.88% | 668.2 | 1.97% |
| -2.34E-01 | 1 | 9695 | 61.79% | 21098.5 | 62.11% |
| -2.30E-01 | 1 | 662 | 4.22% | 1478.8 | 4.35% |
| -2.29E-01 | 1 | 960 | 6.12% | 2132.7 | 6.28% |
| -2.14E-01 | 1 | 4039 | 25.74% | 8848.8 | 26.05% |
| -2.07E-01 | 1 | 3669 | 23.39% | 8046.3 | 23.69% |
| -2.05E-01 | 1 | 11 | 0.07% | 30.6 | 0.09% |
| -1.90E-01 | 1 | 1033 | 6.58% | 2300.3 | 6.77% |
| -1.88E-01 | 1 | 139 | 0.89% | 325.4 | 0.96% |
| -1.78E-01 | 1 | 5253 | 33.48% | 11499.2 | 33.85% |
| -1.76E-01 | 1 | 7022 | 44.76% | 15337.3 | 45.15% |
| -1.72E-01 | 1 | 3539 | 22.56% | 7776.8 | 22.89% |
| -1.68E-01 | 1 | 658 | 4.19% | 1480.4 | 4.36% |
| -1.47E-01 | 1 | 342 | 2.18% | 784.2 | 2.31% |
| -1.41E-01 | 1 | 1783 | 11.36% | 3957.2 | 11.65% |
| -1.38E-01 | 1 | 5630 | 35.89% | 12337.9 | 36.32% |
| -1.34E-01 | 1 | 2872 | 18.31% | 6339.3 | 18.66% |
| -1.26E-01 | 1 | 517 | 3.30% | 1177.9 | 3.47% |
| -1.15E-01 | 1 | 1607 | 10.24% | 3581.4 | 10.54% |
| -1.14E-01 | 1 | 3421 | 21.81% | 7546.2 | 22.21% |
| -1.10E-01 | 1 | 117 | 0.75% | 283.5 | 0.83% |
| -1.01E-01 | 1 | 1591 | 10.14% | 3552.5 | 10.46% |

| | | | | | |
|---|---|---|---|---|---|
| -8.25E-02 | 1 | 520 | 3.31% | 1195.4 | 3.52% |
| -7.70E-02 | 1 | 7383 | 47.06% | 16182 | 47.63% |
| -7.20E-02 | 1 | 7146 | 45.55% | 15672.8 | 46.14% |
| -6.96E-02 | 1 | 98 | 0.62% | 245.8 | 0.72% |
| -6.66E-02 | 1 | 484 | 3.08% | 1120.7 | 3.30% |
| -6.63E-02 | 1 | 2403 | 15.32% | 5352.8 | 15.76% |
| -5.92E-02 | 1 | 6617 | 42.18% | 14538.2 | 42.80% |
| -5.39E-02 | 1 | 312 | 1.99% | 738.1 | 2.17% |
| -5.31E-02 | 1 | 1532 | 9.76% | 3449 | 10.15% |
| -5.24E-02 | 1 | 7566 | 48.22% | 16603.8 | 48.88% |
| -4.41E-02 | 1 | 25 | 0.16% | 74.9 | 0.22% |
| -4.25E-02 | 1 | 141 | 0.90% | 351.7 | 1.04% |
| -4.25E-02 | 1 | 329 | 2.10% | 781.8 | 2.30% |
| -3.58E-02 | 1 | 926 | 5.90% | 2122.5 | 6.25% |
| -3.20E-02 | 1 | 4351 | 27.73% | 9647 | 28.40% |
| -2.89E-02 | 1 | 1014 | 6.46% | 2324.4 | 6.84% |
| -2.65E-02 | 1 | 7835 | 49.94% | 17227.3 | 50.71% |
| -2.32E-02 | 1 | 347 | 2.21% | 833.1 | 2.45% |
| -2.24E-02 | 1 | 1501 | 9.57% | 3413 | 10.05% |
| -2.15E-02 | 1 | 9107 | 58.05% | 19990 | 58.84% |
| -1.91E-02 | 1 | 1411 | 8.99% | 3218.6 | 9.47% |
| -1.75E-02 | 1 | 383 | 2.44% | 920.5 | 2.71% |
| -1.74E-02 | 1 | 1578 | 10.06% | 3591.2 | 10.57% |
| -1.58E-02 | 1 | 1 | 0.01% | 9.6 | 0.03% |
| -1.52E-02 | 1 | 67 | 0.43% | 186.3 | 0.55% |
| -1.23E-02 | 1 | 47 | 0.30% | 138.6 | 0.41% |
| -1.15E-02 | 1 | 236 | 1.50% | 589.2 | 1.73% |
| -1.01E-02 | 1 | 4716 | 30.06% | 10501.9 | 30.91% |
| -8.78E-03 | 1 | 213 | 1.36% | 539.4 | 1.59% |
| -8.15E-03 | 1 | 660 | 4.21% | 1563.8 | 4.60% |
| -7.74E-03 | 1 | 294 | 1.87% | 729.8 | 2.15% |
| -6.94E-03 | 1 | 4486 | 28.59% | 10016.5 | 29.49% |
| -6.78E-03 | 1 | 2655 | 16.92% | 6002.6 | 17.67% |
| -6.75E-03 | 1 | 6383 | 40.68% | 14150.9 | 41.66% |
| -5.75E-03 | 1 | 1414 | 9.01% | 3261.1 | 9.60% |
| -5.65E-03 | 1 | 189 | 1.20% | 488.9 | 1.44% |
| -5.52E-03 | 1 | 787 | 5.02% | 1858.6 | 5.47% |
| -4.88E-03 | 1 | 1967 | 12.54% | 4495 | 13.23% |
| -4.49E-03 | 1 | 238 | 1.52% | 606.1 | 1.78% |
| -4.48E-03 | 1 | 4438 | 28.29% | 9930.7 | 29.23% |
| -2.79E-03 | 1 | 6578 | 41.93% | 14614.6 | 43.02% |
| -2.76E-03 | 1 | 999 | 6.37% | 2351.1 | 6.92% |
| -2.31E-03 | 1 | 6723 | 42.85% | 14937.2 | 43.97% |
| -2.19E-03 | 1 | 5237 | 33.38% | 11705.5 | 34.46% |
| -2.14E-03 | 1 | 6610 | 42.13% | 14695.1 | 43.26% |
| -1.99E-03 | 1 | 694 | 4.42% | 1668.2 | 4.91% |
| -1.82E-03 | 1 | 134 | 0.85% | 368.2 | 1.08% |
| -1.57E-03 | 1 | 367 | 2.34% | 921.9 | 2.71% |
| -1.24E-03 | 1 | 1334 | 8.50% | 3122.7 | 9.19% |
| -1.22E-03 | 1 | 1153 | 7.35% | 2716.3 | 8.00% |
| -1.05E-03 | 1 | 270 | 1.72% | 699.6 | 2.06% |
| -9.16E-04 | 1 | 26 | 0.17% | 97.6 | 0.29% |
| -6.50E-04 | 1 | 2590 | 16.51% | 5936 | 17.47% |
| -5.49E-04 | 1 | 883 | 5.63% | 2122 | 6.25% |

| | | | | | |
|---|---|---|---|---|---|
| -3.24E-04 | 1 | 1292 | 8.24% | 3058.8 | 9.00% |
| -3.19E-04 | 1 | 815 | 5.19% | 1977.6 | 5.82% |
| -2.89E-04 | 1 | 767 | 4.89% | 1869.1 | 5.50% |
| -2.64E-04 | 1 | 793 | 5.05% | 1930.5 | 5.68% |
| -1.84E-04 | 1 | 8218 | 52.38% | 18275.2 | 53.80% |
| -1.31E-04 | 1 | 1001 | 6.38% | 2417.5 | 7.12% |
| -1.08E-04 | 1 | 776 | 4.95% | 1906.4 | 5.61% |
| -8.20E-05 | 1 | 2520 | 16.06% | 5838.3 | 17.19% |
| -6.30E-05 | 1 | 555 | 3.54% | 1403.1 | 4.13% |
| -4.80E-05 | 1 | 1005 | 6.41% | 2444.6 | 7.20% |
| -3.50E-05 | 1 | 6597 | 42.05% | 14817.7 | 43.62% |
| -2.00E-05 | 1 | 6578 | 41.93% | 14794.2 | 43.55% |
| -1.90E-05 | 1 | 105 | 0.67% | 330.8 | 0.97% |
| -1.90E-05 | 1 | 87 | 0.55% | 284 | 0.84% |
| -1.80E-05 | 1 | 3251 | 20.72% | 7500 | 22.08% |
| -1.70E-05 | 1 | 26 | 0.17% | 114.2 | 0.34% |
| -1.60E-05 | 1 | 426 | 2.72% | 1117.7 | 3.29% |
| -1.60E-05 | 1 | 210 | 1.34% | 596.7 | 1.76% |
| -1.10E-05 | 1 | 354 | 2.26% | 949.3 | 2.79% |
| -9.00E-06 | 1 | 5079 | 32.37% | 11544.9 | 33.98% |
| -7.00E-06 | 1 | 6173 | 39.35% | 13945.5 | 41.05% |
| -5.00E-06 | 1 | 3330 | 21.23% | 7706.3 | 22.68% |
| -3.00E-06 | 1 | 2549 | 16.25% | 5978.4 | 17.60% |
| -1.00E-06 | 1 | 922 | 5.88% | 2309.3 | 6.80% |
| -1.00E-06 | 1 | 2464 | 15.71% | 5813 | 17.11% |
| 0.00E+00 | 1 | 948 | 6.04% | 2386 | 7.02% |
| 0.00E+00 | 1 | 319 | 2.03% | 901.6 | 2.65% |
| 0.00E+00 | 1 | 3939 | 25.11% | 9139.8 | 26.90% |
| 0.00E+00 | 1 | 301 | 1.92% | 863 | 2.54% |
| 0.00E+00 | 1 | 3653 | 23.28% | 8536.1 | 25.13% |
| 0.00E+00 | 1 | 3030 | 19.31% | 7157 | 21.07% |
| 0.00E+00 | 1 | 4172 | 26.59% | 9727.3 | 28.63% |
| 0.00E+00 | 1 | 3897 | 24.84% | 9129.7 | 26.87% |
| 0.00E+00 | 1 | 4395 | 28.01% | 10233.6 | 30.12% |
| 0.00E+00 | 1 | 145 | 0.92% | 492.1 | 1.45% |
| 0.00E+00 | 1 | 4306 | 27.45% | 10067.8 | 29.64% |
| 0.00E+00 | 1 | 1964 | 12.52% | 4837 | 14.24% |
| 0.00E+00 | 1 | 812 | 5.18% | 2168.5 | 6.38% |
| 0.00E+00 | 1 | 213 | 1.36% | 694.9 | 2.05% |
| 0.00E+00 | 1 | 2670 | 17.02% | 6470.7 | 19.05% |
| 0.00E+00 | 1 | 2253 | 14.36% | 5546.8 | 16.33% |
| 0.00E+00 | 1 | 2321 | 14.79% | 5705.6 | 16.80% |
| 0.00E+00 | 1 | 6494 | 41.39% | 14988.9 | 44.12% |
| 0.00E+00 | 1 | 2027 | 12.92% | 5137.2 | 15.12% |
| 0.00E+00 | 1 | 921 | 5.87% | 2535.3 | 7.46% |
| 0.00E+00 | 1 | 1380 | 8.80% | 3639.3 | 10.71% |
| 0.00E+00 | 1 | 2063 | 13.15% | 5260.2 | 15.48% |
| 0.00E+00 | 1 | 1759 | 11.21% | 4549.9 | 13.39% |
| 0.00E+00 | 1 | 1678 | 10.70% | 4366.4 | 12.85% |
| 0.00E+00 | 1 | 463 | 2.95% | 1457.5 | 4.29% |
| 0.00E+00 | 1 | 2587 | 16.49% | 6897.9 | 20.31% |
| 0.00E+00 | 1 | 3427 | 21.84% | 8489.5 | 24.99% |
| 0.00E+00 | 1 | 1886 | 12.02% | 5069.8 | 14.92% |
| 0.00E+00 | 1 | 2491 | 15.88% | 6550.7 | 19.28% |

| | | | | | |
|---|---|---|---|---|---|
| 0.00E+00 | 1 | 8439 | 53.79% | 19467.7 | 57.31% |
| 0.00E+00 | 1 | 1486 | 9.47% | 4098.9 | 12.07% |
| 0.00E+00 | 1 | 904 | 5.76% | 2736.5 | 8.06% |
| 0.00E+00 | 1 | 1374 | 8.76% | 3656.8 | 10.76% |
| 0.00E+00 | 1 | 1455 | 9.27% | 3899.7 | 11.48% |
| 0.00E+00 | 1 | 1908 | 12.16% | 4929.5 | 14.51% |
| 0.00E+00 | 1 | 1283 | 8.18% | 3490 | 10.27% |
| 0.00E+00 | 1 | 2119 | 13.51% | 5488.4 | 16.16% |
| 0.00E+00 | 1 | 2677 | 17.06% | 6756.6 | 19.89% |
| 0.00E+00 | 1 | 1421 | 9.06% | 3841.9 | 11.31% |

**es with Motif**
